# No global collapse of food webs across the Permian–Triassic Mass Extinction

**DOI:** 10.64898/2026.02.24.707709

**Authors:** Baran Karapunar, Tanya Strydom, Andrew P. Beckerman, Andy Ridgwell, Paul B. Wignall, Jennifer A. Dunne, Crispin T. S. Little, Pincelli Hull, Catalina Pimiento, Alexander M. Dunhill

## Abstract

The Permian–Triassic mass extinction (PTME), the Earth’s most severe biotic crisis associated with extreme environmental perturbations, eliminated >80% of marine species^1^. However, whether it triggered a globally pervasive top-down collapse of marine food webs, and whether recovery proceeded through bottom-up reassembly, remain unresolved^2-4^. Here we reconstruct spatially explicit metacommunity food webs from seven regions spanning equatorial to high latitudes to test how extinction dynamics and ecosystem reorganization varied geographically. By integrating estimates of community structure and species interactions, we provide direct inference on trophic disruption across the PTME. Despite catastrophic species loss and flattening of the latitudinal diversity gradient^5^, trophic collapse was not globally uniform, and higher trophic levels were not globally truncated. Instead, extinction selectivity was spatially heterogenous and tracked environmental severity. Benthic, low-motility herbivores with limited respiratory capacity were disproportionately lost, consistent with intensified warming, deoxygenation and disruption of primary productivity under elevated pCO_2_. Mid-to high-latitude communities became top-heavy and structurally complex, whereas tropical systems remained bottom-heavy and less robust to secondary extinction. These results demonstrate that trophic disruption and recovery were geographically structured, mediated by environmental forcing, species traits and pre-extinction food-web architecture, with implications for predicting marine ecosystem responses to ongoing climate change.

## Introduction

Life on Earth has undergone multiple mass extinction events, and the Permian–Triassic mass extinction (PTME; ∼252 Ma) was the most catastrophic, eliminating an estimated 80% of marine animal species^1^. The PTME is linked to volcanic-driven extreme environmental disturbances^6^, including extreme global warming of 10–15°C^7,8^, widespread ocean deoxygenation^9^ and euxinia^10^, changes in ocean productivity^11,12^, and ocean acidification^13^. Due to these global environmental change features, the PTME is often viewed as a partial analogue for how the marine biosphere might respond to present and future climate change^14^.

Most PTME analyses have drawn their conclusions from global scale analyses^15-19^, averaging extinction drivers across environmentally heterogeneous regions. Spatial gradients in warming and deoxygenation, e.g., more intense thermal stress at low latitudes and expanded oxygen limitation at higher latitudes^10,20^, would have interacted with species composition, physiological tolerances and life-history strategies to produce regionally distinct extinction pressures, contributing to among-clade variation in extinction rates and potentially the collapse of the latitudinal diversity gradient^5^. How this environmental and ecological heterogeneity translated into spatial variation in extinction dynamics remains insufficiently tested.

Analyses of the PTME have typically quantified taxonomic and functional losses and often treated all extinctions as primary extinctions, ignoring community structure that may buffer losses or drive secondary extinctions. Accounting for community structure is a way to define ecological interactions that can strongly modulate extinction risk, influencing which taxa disappear, when losses occur, and how cascading secondary extinctions can arise^21-23^. Ecological community structure also underpins ecological evolutionary dynamics that are central to understanding how biodiversity, ecological structure, and ecosystem function recover after mass extinctions^22,23^. Ignoring this structure and these dynamics risks obscuring mechanisms such as habitat loss, physiological or thermal stress, prey depletion, and increased predation that may have mediated extinction trajectories^21^.

A key prediction about the PTME is the truncation of upper trophic levels, with recovery proceeding stepwise from primary producers upward over as much as eight million years^2^. This prediction derives from the fossil record in South China^2^ and was partly confirmed by a further study from this region, which proposed trophic collapse following the second extinction pulse and heightened vulnerability to secondary extinction cascades^4^. However, growing fossil evidence suggests this view may be overly generalised^3,24^. Nektonic clades occupying higher trophic levels, including fishes^25^ and ammonoids^26^, appear to have been comparatively less affected, recovered rapidly, and became proportionally more abundant in the immediate aftermath^27^. A shift in dominance from low-to high-motility organisms^17^, together with limited body-size reduction in motile nektonic taxa compared to selective losses among smaller, non-motile benthic groups such as foraminifers and brachiopods^16,25^, further indicates uneven ecological impacts and recovery trajectories. However, this counter evidence for a lack of complete trophic collapse derives either from global diversity curves^25,26^ or from later Early Triassic intervals^3,24,27,28^ and limited attention has been given to geographic variation in trophic structure across the extinction interval itself.

Whether such trophic disruption^4^ was globally pervasive or spatially variable, and whether recovery followed a bottom-up sequence^2,3^, remains unresolved^24^. Here, we explicitly integrate estimates of community structure and species interactions into a spatially resolved palaeoecological analysis of the PTME to deliver strong inference on how extinction dynamics and ecosystem reorganization varied across regions. We reconstruct seven geographically distinct marine metacommunities spanning equatorial to high palaeolatitudes and infer trophic networks before, during, and after the extinction interval using the Paleo Food Web Inference Model (PFIM)^23,29^. We address three primary questions: (i) Did the PTME produce a global, top-down collapse of marine food webs?; (ii) How heterogeneous was extinction selectivity among different regions and palaeolatitudes?; and (iii) Were post-extinction trophic networks reorganized in a uniform or spatially variable manner?

To address these questions, we reconstructed food webs from fossil trait data and simulated 17 trait-informed extinction scenarios based on selectivity against key biological traits. We used these simulations to estimate both direct species losses and the cascading effects that followed^23^, i.e., secondary extinctions that were triggered when consumers lost all prey items. We compared these simulations with observed outcomes of extinction in the fossil record to estimate the most plausible extinction targets and sequence in each region. We further quantified spatial variation in warming, deoxygenation, and productivity change using the coupled climate–Earth system model EcoGEnIE^30^ to deliver parallel insight into bottom-up processes across the PTME. Together, this analysis framework enables a spatially explicit assessment of how environmental forcing and community structure interacted to shape marine ecosystem extinction dynamics, collapse, and reorganisation during the most severe mass extinction in Earth history.

## Results and discussion

### No global collapse of trophic structure

Species loss and turnover, estimated by comparing pre- and post-extinction community composition across the PTME, was extensive at all sites, with 68–92% of pre-extinction taxa lost during the Changhsingian and 89–96% lost by the Griesbachian (Table S4). In Russia, all pre-extinction taxa were eliminated. Despite this profound taxonomic loss and species turnover, trophic structure did not collapse globally. Most regions retained four trophic levels across the extinction interval (Figure 1), indicating persistence of higher trophic levels even as species richness declined sharply. Complete loss of the highest trophic level occurred only in the tropical Meishan and Türkiye communities, and even there, trophic truncation was partial, with ecological roles comparable to apex predators persisting in Meishan. Mid- and high-latitude communities consistently retained upper trophic levels, providing no support for a globally pervasive, top-down collapse of marine predators^2^.

**Figure 1.**
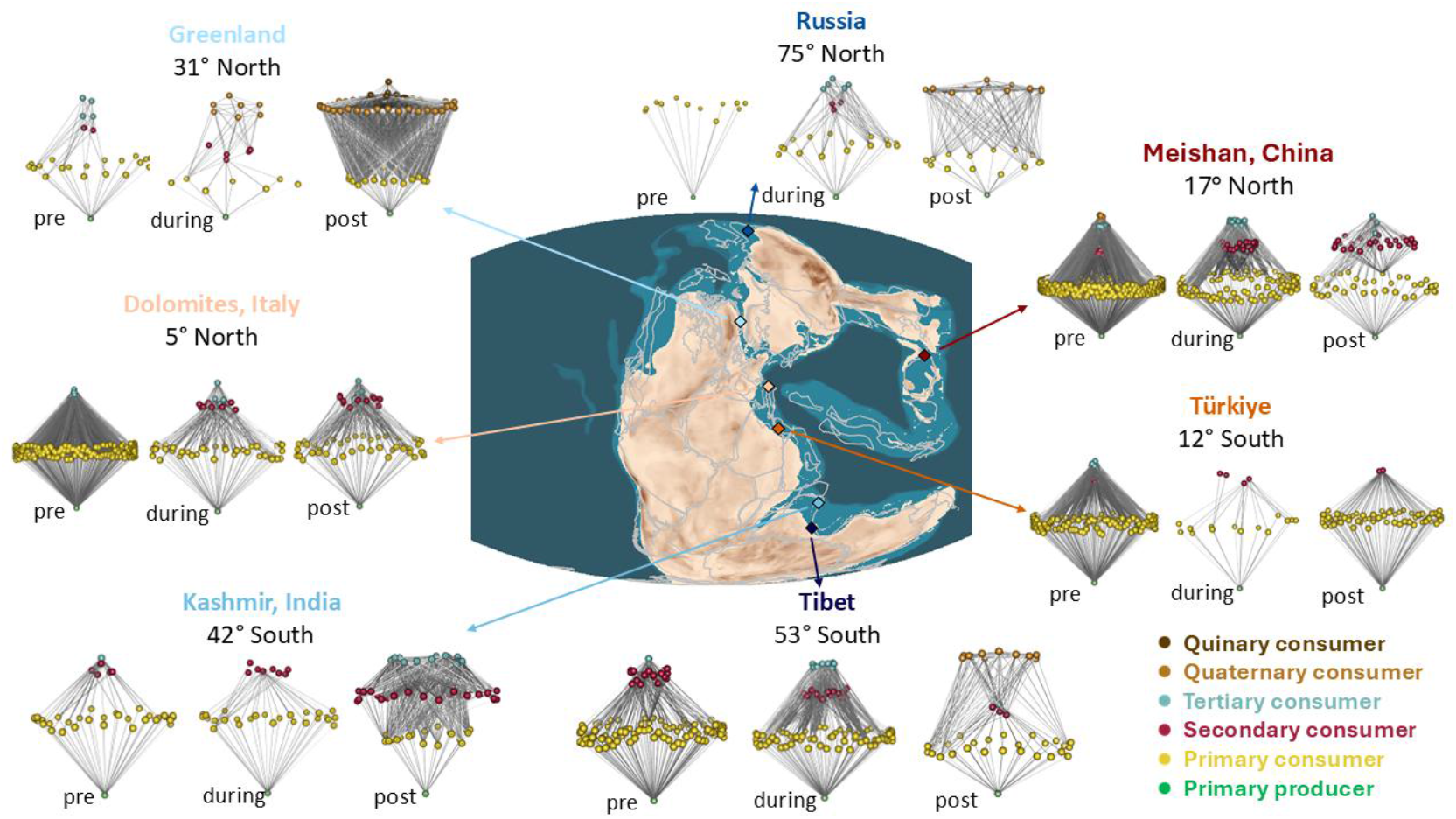
Paleo Food web Inference Model (PFIM)^23,29^ reconstructed food webs at seven regions across the PTME. **pre:** Changhsingian pre-extinction interval, **during:** mass extinction interval (Changhsingian or Changhsingian–earliest Griesbachian), **post:** Griesbachian post-extinction interval.

Post-extinction food webs were nonetheless reorganised, and this reorganisation varied with palaeolatitude. In most regions, primary consumers (e.g., filter feeders, deposit feeders, herbivores), which are largely non-motile and benthic, declined disproportionately relative to higher trophic guilds (Fig. 2k). A regional exception occurs in Türkiye, where elevated ostracod diversity likely reflects a taxonomic over splitting effect. Overall, the PTME was characterized not by uniform, top-down trophic collapse, but by spatially variable restructuring, underscoring the risk of inferring global dynamics from single localities^4^.

**Figure 2.**
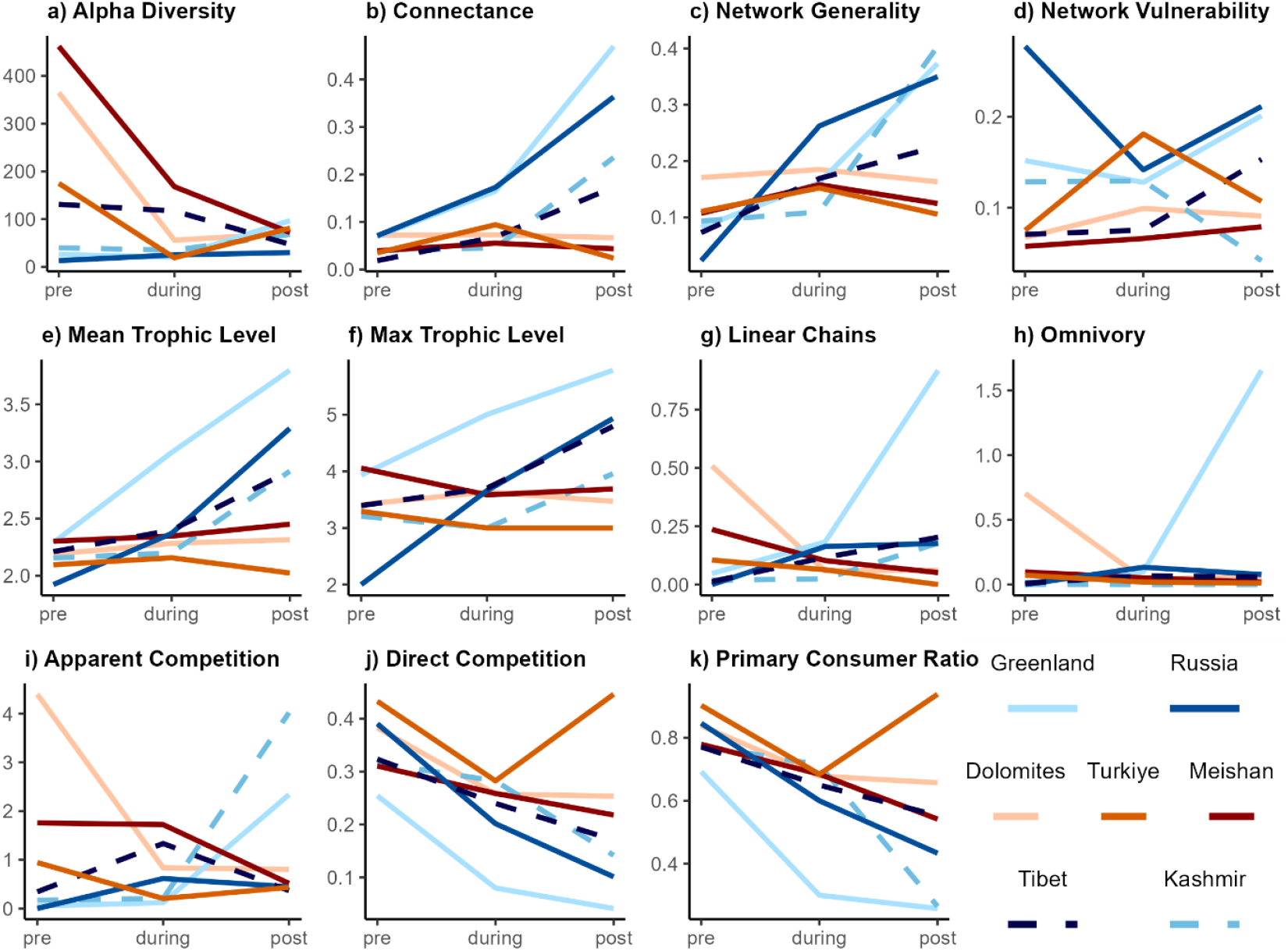
Food web metrics of species-level metacommunities including a primary producer node across the PTME. Low latitudinal regions are coloured in red tones; mid-high latitude regions are coloured in blue tones.

### Spatially varied extinction selectivity

Simulation of multiple primary extinction scenarios (see Extinction selectivity analysis in Supplementary Methods) reveal strong spatial heterogeneity in trait-based selectivity from the pre-extinction to mass extinction interval (Fig. 3a-b). Different communities experienced selective losses along distinct ecological and physiological axes, contrary to multiple and conflicting previous suggestions of a single global axis such as low physiological buffering capability^15^, limited motility^17^, low respiratory capacity^19^, or large size^16^. In high-latitude Greenland and Tibet, non-motile benthic taxa were preferentially lost, whereas in Meishan and Kashmir, extinction disproportionately affected organisms with low respiratory capacity, including foraminifera, echinoderms, and bryozoans, though extinction in Kashmir also included a strong random component. In Russia, the entire fauna was comprised of non-motile primary consumers, all of which were eliminated, while in Türkiye, unbuffered specialists such as brachiopods were selectively removed. Conversely, in the tropical Dolomites, taxa with high respiratory capacity, including vertebrates and cephalopods, were disproportionately lost. These patterns highlight that primary extinction pressures varied substantially across regions and palaeolatitudes.

**Figure 3.**
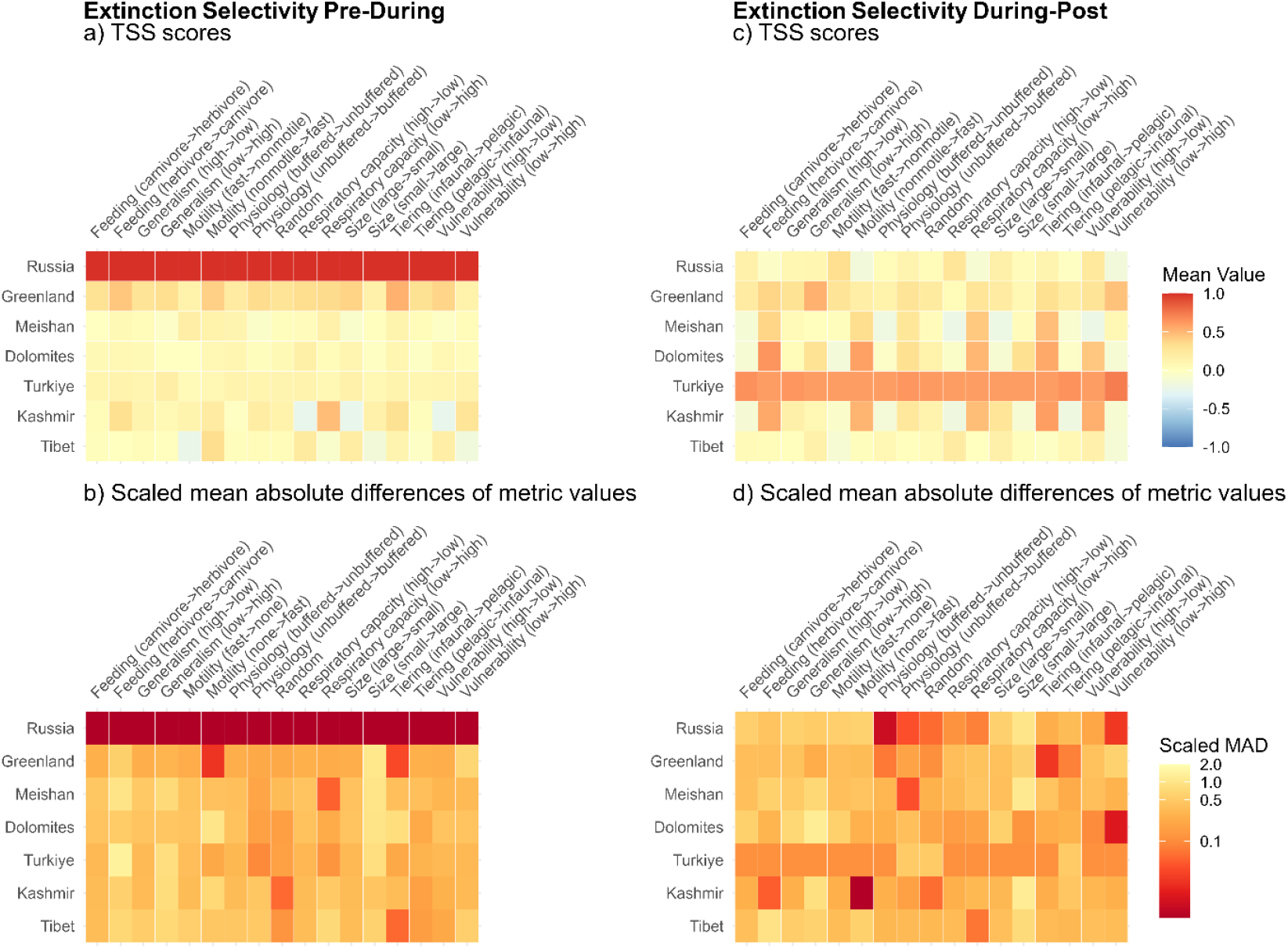
Trait-based extinction cascade simulations (17 scenarios) across the PTME, comparing pre-to during-extinction (a,b) and during-to post-extinction (c,d) intervals. Scenarios (x-axis) directionally targeted ecological or physiological traits, or were random, with secondary extinctions emerging from primary losses. The alignment of predicted outcome versus empirical networks was assessed using node identity (a,c) and eight network metrics (b,d); red indicates stronger fit. In Russia, complete taxon loss makes all scenarios equally supported in the first interval.

During the extinction interval itself, selectivity remained heterogeneous (Fig. 3c-d) but primary selectivity varied compared to the previous interval at the same locality, due to changes in community composition and environmental conditions. For example, benthic, non-motile herbivores were most affected in Kashmir and the Dolomites, benthic taxa with low buffering capacity against acidification in Meishan, benthic specialist feeders in Greenland, fast-moving buffered taxa in Russia, and large, high-respiratory capacity taxa in Tibet. In Türkiye, extremely low survivorship limited robust inference of selectivity patterns.

Despite the pronounced regional variability, is the network perspective and high-resolution data make it possible to identify some broad themes in primary extinction selectivity throughout the mass extinction interval. Primary extinction preferentially impacted taxa with limited motility and physiological constraints, particularly non-motile benthic primary consumers with low respiratory capacity (e.g., Foraminifera, Brachiopoda, Bryozoa, Cnidaria), supporting some consistency with global trait analyses^15,17,19^. In contrast, nektonic predators occupying higher trophic levels in mid-high latitude communities, such as fishes and conodonts, were generally less affected, potentially facilitating the apparent rapid post-extinction diversification of these groups seen in the fossil record^25-27,31^.

Overall, our results demonstrate that primary extinction selectivity across the PTME was spatially and temporally heterogeneous, shaped by interactions between organismal traits and temporal changes in community compositions and regional environmental stressors. While the primary target in high latitudes was benthic non-motile organisms across the onset of the extinction interval, the same functional group was targeted in low latitude communities during the extinction interval itself. Rather than a single global extinction filter, distinct palaeolatitudinal and environmental settings imposed distinct selective pressures, producing geographically structured extinction patterns that influenced subsequent ecosystem reassembly.

### Spatially varied community restructuring

Following the PTME, species richness converged across palaeolatitudes as tropical diversity declined sharply and some mid-to high-latitude regions experienced modest increases (Fig. 2), thus supporting the idea of a flattened Early Triassic latitudinal diversity gradient^5^. Despite this convergence in alpha diversity and a marked decrease in beta diversity (Figure S10, S11) driven by increased cosmopolitanism^32^, trophic restructuring was spatially variable. Low-latitude communities (i.e., Meishan, Türkiye, Dolomites) retained comparatively bottom-heavy trophic structure into the Griesbachian, even after disproportionate losses among benthic primary consumers. In contrast, mid-to high-latitude communities (Tibet, Kashmir, Greenland) became increasingly top-heavy, dominated by higher trophic levels as newly originated and immigrating taxa were disproportionately predatory (Fig. 1; Supplementary Material).

In these higher-latitude systems, origination and immigration of predatory fishes, temnospondyls, and cephalopods generated quaternary and quinary trophic levels, increasing both mean and maximum trophic level (Fig. 2e–f). Network metrics indicate that these food webs developed greater vertical complexity, with higher connectance, elevated proportions of omnivory, and increased generalism relative to pre-extinction configurations (Fig. 2). Higher connectance at mid-to high latitudes was driven primarily by the proliferation of generalist predators, which lengthened food chains and increased the number of potential energy pathways. In contrast, tropical networks remained comparatively shorter and less vertically expanded, maintaining more bottom-heavy structures despite overall biodiversity loss. Thus, although trophic levels were not globally truncated, the balance of energy flow shifted latitudinally, with high-latitude communities becoming predator-rich and vertically complex, and low-latitude systems remaining more benthos-dominated.

These structural differences were accompanied by shifts in interaction patterns estimated via motif distributions in the networks. Apparent competition among prey increased in several higher-latitude communities where top and intermediate predatory taxa dominate both predator and prey roles, while direct competition within networks generally declined, reflecting selective loss of primary consumers (Fig. 2). However, there was an increase in direct competition among the few remaining higher trophic groups in low latitude communities (Fig. S12), due to declines in prey. At higher latitudes, the proliferation of generalist predators increased interaction overlap at upper trophic levels, potentially promoting niche partitioning, and thus potentially driving increases in diversity and disparity among nektonic groups. This interpretation is consistent with fossil evidence for rapid Early Triassic radiations of fishes^25^, ammonoids^26^, predatory gastropods^33^, and marine tetrapods^31^, and with documented expansion of morphospace among secondary and tertiary consumers (such as conodonts and ammonoids)^18^. Our modelled food webs support limited diversification among low-trophic benthic groups in tropical settings, likely associated with reduced interspecific competition^34^, but indicate that this pattern was not globally uniform nor consistent across all trophic levels.

Denser networks with increased connectance and higher levels of generalism in post-extinction communities are consistent with patterns seen across other extinction events in the terrestrial ^35,36^ and marine fossil record^23^. These patterns reflect a high proportion of generalist predators and cannot be attributed to differential preservation of nektonic taxa. Although local benthic deoxygenation may have improved preservation in some regions, including high latitudes, it was not globally uniform^20,37^ (Figure 5). Moreover, benthic deoxygenation would have reinforced the observed trophic restructuring by selectively reducing benthic diversity. A similar but weaker top-heavy reorganization during the Early Toarcian extinction^23^ suggests a broader post-extinction tendency for increased richness at higher trophic levels. The strongly inverse-pyramidal trophic structure following the PTME likely reflects the exceptional environmental severity of the Permian–Triassic crisis, including extreme low-latitude warming, intensified high-latitude deoxygenation, and reduced productivity^27^.

Changes in trophic structure are also reflected in contrasting patterns of intrinsic robustness of each community. Characterising robustness simulations under random species removal show that pre-extinction communities were uniformly fragile: removal of a relatively small fraction of taxa could trigger extensive secondary extinctions (Figure 4). This low intrinsic robustness suggests that late Palaeozoic marine ecosystems may have been more susceptible to cascading secondary collapse, potentially amplifying biodiversity loss under extreme environmental stress and contributing to the exceptional severity of the PTME. Following the extinction, intrinsic robustness appear to increase in mid-to high-latitude communities as networks became more top-heavy, vertically complex, and highly connected, whereas tropical communities exhibited slight declines in robustness and thus may have remained vulnerable to further secondary extinction cascades^4^ (Fig. 4). Overall, robustness was positively associated with longer food chains, greater omnivory, and higher connectance, network properties known to buffer systems against secondary cascades^38^.

**Figure 4.**
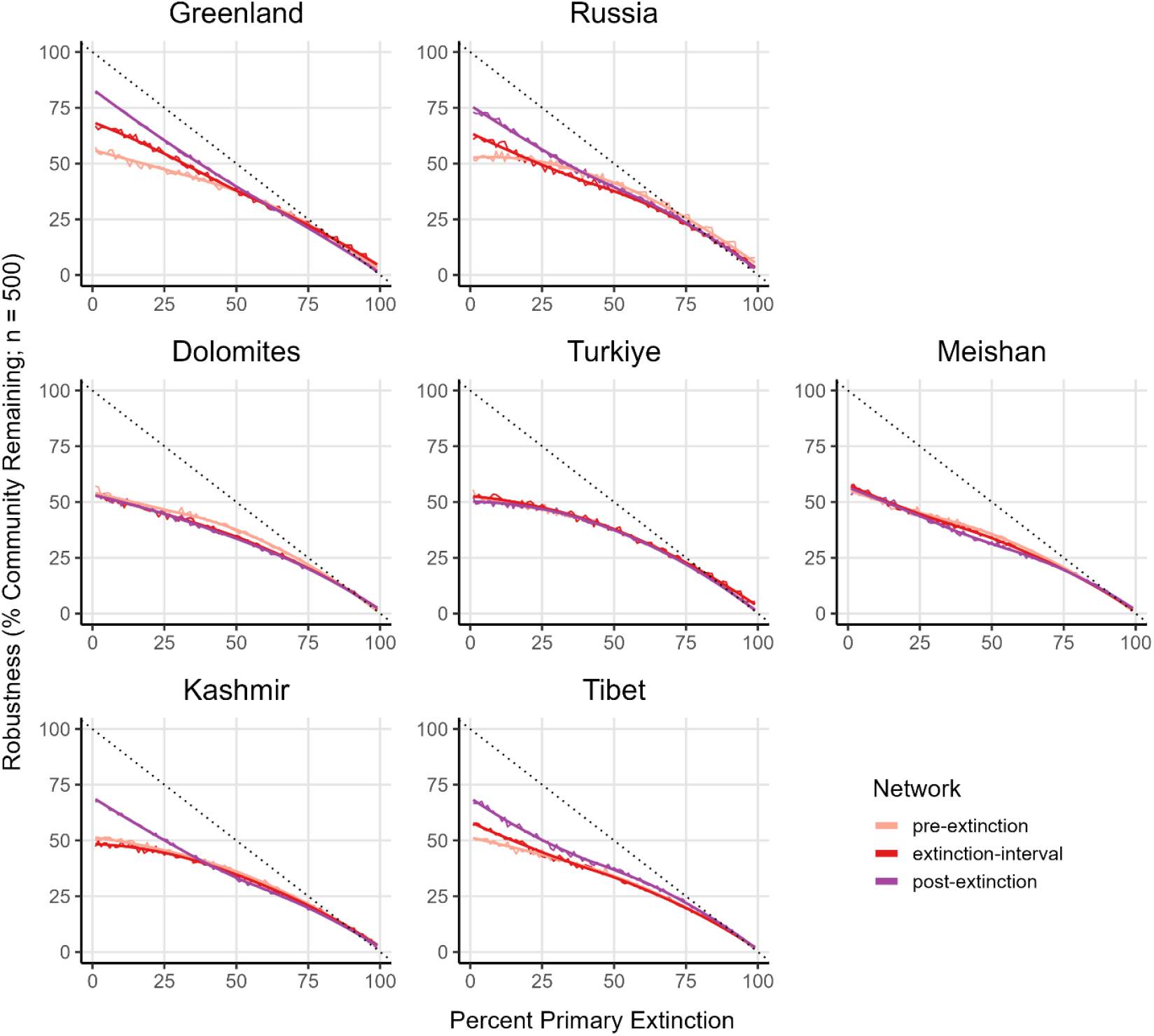
Robustness against random primary extinctions for the three communities across the PTME at each site, drawn by LOESS fit of mean values after 500 trials. Robustness curves were produced by simulating random primary extinctions in three consecutive communities in each region and measuring secondary extinctions (i.e., extinctions due to all prey loss) because of primary extinctions.

Although, all pre-extinction communities showed similar bottom-heavy trophic structure, post-extinction ecosystems did not recover towards a single trophic structure. Instead, mid-to high-latitude regions reorganized into predator-dominated, vertically complex, and more robust food webs, whereas tropical communities retained more bottom-heavy and comparatively less robust configurations that remained vulnerable to secondary collapse^4^. The PTME therefore neither reflects a global trophic collapse, nor a uniform recovery of pre-extinction community structure^24^, but the PTME is characterised by spatially divergent restructuring. Its extreme severity may have arisen from the combination of intense environmental forcing and the comparatively low pre-extinction robustness of Palaeozoic marine ecosystems, followed by the emergence of more vertically complex and interaction-rich Triassic food webs.

### Environmental drivers of the variance

The spatially divergent extinction and recovery patterns identified in our network analyses are consistent with geographically heterogeneous environmental forcing during the PTME. The extreme environmental perturbations including warming, deoxygenation, acidification, and decline in POC quantity and quality, interacted with trait-mediated metabolic and ecological filters to generate spatially heterogeneous extinction selectivity and divergent community restructuring. Rather than producing uniform trophic collapse, environmental gradients reshaped communities differently across latitudes, enabling mid-to high-latitude ecosystems to reorganize into more vertically complex and robust configurations while tropical systems remained comparatively bottom-heavy and vulnerable to further collapse^4^.

EcoGEnIE simulations indicate pervasive ocean acidification^13^ and deoxygeneation^9,10,20^ in all study regions (Figure S17), globally elevated sea-surface temperatures with extreme thermal stress in the tropics^7,8^, and highest rates of warming at mid-to high latitudes^20^ (Fig. 5, S17). High metabolic stress imposed by combination of high temperatures and low oxygen conditions in low latitudes and decline in POC export in northern high latitudes likely interacted with species’ ecological traits and community structure to produce the regionally distinct selectivity regimes and trophic reorganizations described in this study.

**Figure 5.**
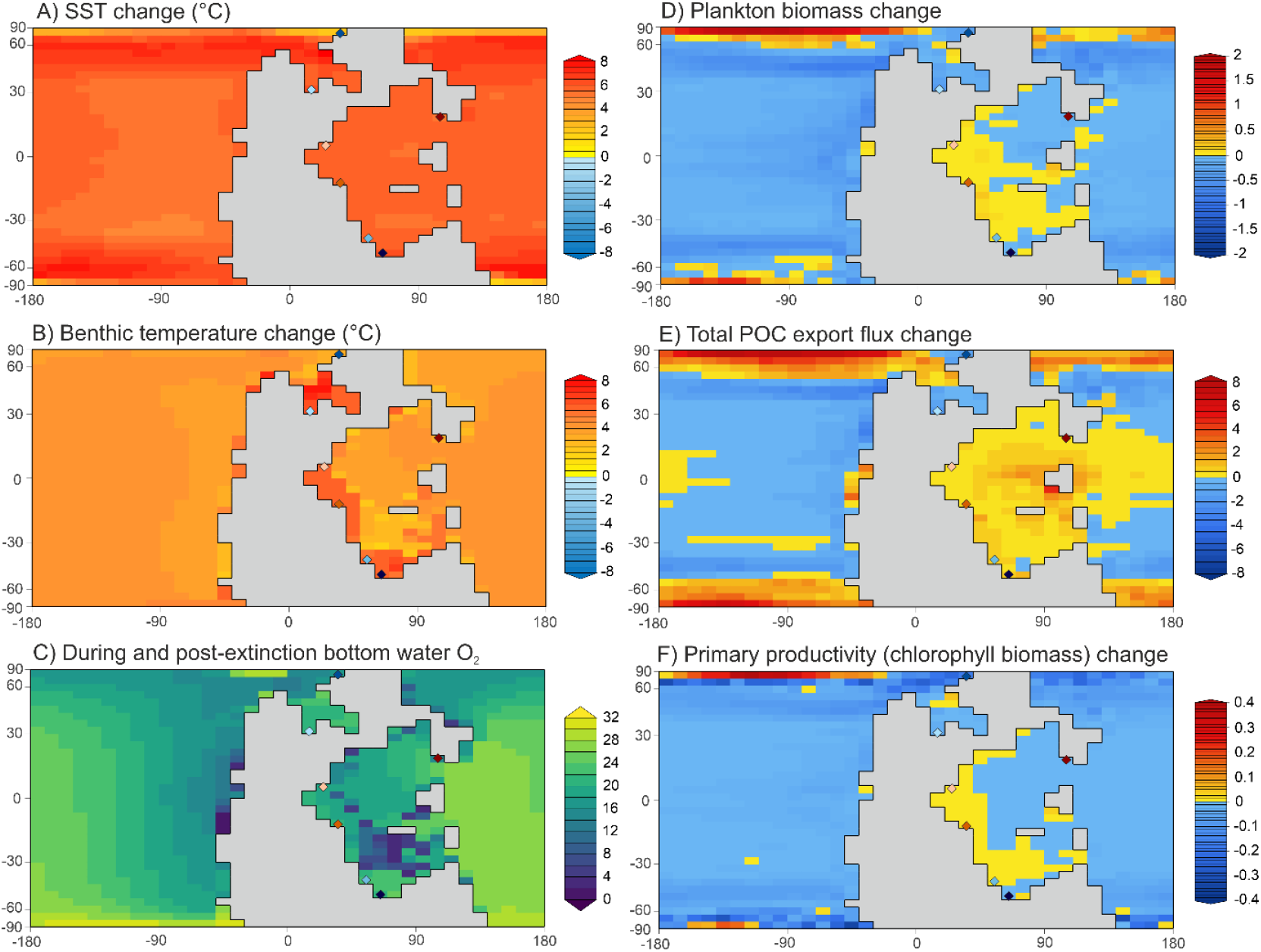
Modelled environmental and ecological variables across the PTME using EcoGEnIE under increase from a 2-fold modern *p*CO2 in the pre-extinction interval to 10-fold modern *p*CO2 during and in the post-extinction interval. **A)** Sea surface temperature change (°C); **B)** benthic temperature change (°C); **C)** benthic oxygen during and post-extinction (mol/kg); **D)** plankton biomass (total Carbon mmol/m^3^); **E)** POC (10^11^ mol/year); **F)** change in primary productivity measured as chlorophyll biomass (mg chl/m^3^).

Intense global warming at the PTME^7,8^ would have driven poleward range shifts among mobile nektonic taxa^25^ and planktonic consumers^39^, consistent with post-extinction proliferation of predatory fishes, ammonoids and tetrapods at higher latitudes^3,25-27,31^ and the emergence of top-heavy, vertically complex food webs in these regions. Local disappearances of taxa such as *Bobasatrania* and *Acrodus* in the tropical Dolomites but persistence at higher latitudes^40^ support redistribution rather than global extinction. In contrast, benthic primary consumers with limited motility and low respiratory capacity, particularly filter and deposit feeders, were selectively lost across all latitudes (Figure 2), reflecting dispersal limitations of planktonic larvae and vulnerability to extreme environmental change (warming, oxygen limitation, reduced food supply and quality).

Although hypoxia has been proposed as a unifying kill mechanism^20^, our simulations do not support a single global extinction filter. Large body size, for example, was not universally targeted, despite metabolic expectations under warming and deoxygenation^41^. Instead, size selectivity appears restricted to certain benthic herbivores (i.e. brachiopods and bivalves)^16^, reinforcing that extinction risk was mediated by combinations of physiology, ecology, life-history, and regional environmental context rather than any single trait.

Food-web restructuring may also have been shaped by spatially variable primary productivity responses, as a bottom-up disruption of energy flow could explain selective extinction of benthic herbivores and suppression of higher trophic-level recovery. Geochemical proxies^11,12^ and EcoGEnIE model outputs indicate reduced productivity in some regions (e.g., Meishan, Greenland) but stability or increases in productivity elsewhere. Moreover, shifts from eukaryotic to prokaryotic-dominated primary production likely reduced food quality, disproportionately affecting suspension-feeding benthos even where total productivity did not decline. Beyond temperature-driven range shifts of pre-extinction tropical taxa to mid–high latitudes, higher-latitude siliciclastic settings may have been partly buffered against acidification^42^ by silicate weathering and plankton blooms^43^, potentially acting as refugia that supported predator-rich networks, while energetically demanding predators were extirpated at low latitudes through secondary extinction cascades driven by prey declines during the mass extinction interval^15,44,45^.

## Conclusion

Our analyses centring community structure, species identity, and functional traits suggest that the PTME caused catastrophic biodiversity loss but did not trigger a global collapse of marine food webs. Instead, extinction selectivity and its impact on community structure was spatially heterogenous and aligned with predicted geographic variation in environmental forcing. Benthic primary consumers were disproportionately eliminated globally, whereas mobile, generalist predators persisted and diversified, particularly at mid-to high latitudes, where environmental buffering facilitated the development of more vertically complex networks that are robust against secondary extinction cascades. Tropical systems remained comparatively bottom-heavy and vulnerable to secondary extinction cascades^4^. The severity of the PTME thus reflects the interaction between extreme environmental forcing, clade-specific susceptibilities, and the low intrinsic robustness of Palaeozoic ecosystems, rather than indiscriminate or globally uniform trophic collapse. Our findings transform the prevailing selective narrative of the PTME drawn by global level analyses, which masked spatially heterogeneous, network and trait-mediated ecological reorganisation. It further shows that the long standing “food webs collapsed” narrative derived from findings in a single region^2,4^ cannot be extrapolated to other regions. Accounting for spatial variation in community composition, species traits, and trophic structure is essential for interpreting past mass extinctions and anticipating modern marine ecosystem responses to ongoing environmental change.

## Supporting information

Karapunar_et_al_PT food web_Supplementary_Material_1_Data_and_Results

