## Supplementary material for "No global collapse of food webs across the Permian–Triassic Mass Extinction": Method

### Methods

#### Metacommunity Datasets

The metacommunity datasets were collected from seven regions: Dolomites [Italy], Türkiye, Meishan [China], Kashmir [India], Tibet, Greenland, Russia. We compiled lists of species reported within a region either from a single geological formation in an interval (Italy, China, Russia, Kashmir, Russia) or from several formations (Tibet, Türkiye, Greenland) spanning the Permian Triassic mass extinction (PTME) interval. Since the Meishan Section (Zhejiang, China) is the most heavily sampled Permian–Triassic section among the studied regions, and therefore yields the most-species rich community, we did not supplement the Meishan dataset by including species from other locations in the same region, to balance the sampling effort among regions. A total of 1,783 species occurrences were compiled using our own collections stored in public institutes and publications from these seven regions across the Changhsingian–Griesbachian [Dolomites: 439 species, Türkiye: 263 species, Meishan: 566 species, Kashmir: 115 species, Tibet: 220 species, Greenland: 129 species, Russia: 51 species]. After matching the species names in all lists, the final dataset includes 1,526 from the Changhsingian–Griesbachian (Table S2). References used to compile the occurrence datasets are listed in the Supplementary Material 1, and each occurrence is linked to its specific source in a dedicated column of the dataset.

The metacommunity composition is based solely on the sampled fossil record, which is subject to several filters, such as fossilization, preservation, and sampling. Although, the metacommunity compositions inferred from the fossil record of the study regions is not fully comparable to living communities in a modern ecological sense, they are considered representative of the once living communities.

#### Division into three intervals

Each metacommunity dataset was divided into three intervals (pre-extinction interval, extinction interval and post-extinction interval). The PTME is a diachronous event whereby, in some locations, the main extinction pulse is in the latest Changhsingian (Greenland[1], Türkiye[2]). Whereas, in other sections, the main phase is assumed to cross the Permian/Triassic boundary with a second major pulse in extinction rate in the earliest Griesbachian (Tibet[3], Meishan[4], Italy[5], Türkiye[2]). In Tibet, Meishan and Dolomites sections, the mass extinction interval was placed following these previous studies. However, to place the mass extinction interval in the sections where the main extinction phase is in the latest Changhsingian, we used combination of geochronologic and formation boundaries. In Greenland, Kashmir and Russia, the formation boundary is below the Changhsingian–Griesbachian stage boundaries. In these regions, the interval between the formation and stage boundaries were assigned to extinction interval (i.e., the Changhsingian parts of the younger formations are assigned to as extinction interval). Palaeoenvironmental setting of studied regions and the stratigraphic correlations of the three intervals are provided in the Supplementary Material 1.

#### Ecological traits

Each species was assigned four ecological traits which are used to parameterise the food web model: tiering, motility, feeding, following Bambach's Ecospace model[6], and a categorical body-size category. To avoid unnecessary complexity, categories were simplified where finer distinctions did not affect trophic interpretations: surficial and erect forms were grouped as epifaunal; cemented, byssate, and other attached taxa were treated as "attached"; deposit-, filter-, and grazer-herbivores were combined as "herbivores"; and grazer-carnivores (such as pleurotomariid gastropods), active hunters, and passive predators (such as cuspidariid bivalves) were all assigned to a single "carnivore" category without differentiating the predatory habit (such as crushing, chipping, sucking, tearing, piercing).

Ecological traits were interpreted based on morphological features, phylogenetic affinity, direct evidence (such as gut contents, isotopes), and predation traces (such as drill holes, scratch or bite marks) derived from the literature[7-9] for different groups (Bivalvia[10,11], Gastropoda[12-16], Cephalopoda[17], Echinodermata[18-20], Brachiopoda[21-23], Ostracoda[24], Bryozoa[25], Microconchida[26], Foraminifera[27]). Following Bambach et al.[6], Bryozoa, Foraminifera, Ophiuroidea, Crinoidea, and sponges were classified as herbivores, and Echinoidea as omnivores. Ecological assignments for fish, temnospondyls, and ichnofossils follow the sources[28,29] used to compile their occurrences, with species-level references provided in the dataset. Isotopic data indicate that conodonts occupied low trophic levels and preyed on zooplankton[30,31], consistent with functional-morphological evidence for predation[32], although some early groups may not have been predators[33]. Accordingly, conodonts were assigned to a "microcarnivorous" category.

In addition to the ecological traits used to model trophic interaction, we also compiled the physiological traits of respiratory protein and physiological buffering capability (against decreasing carbonate ions [$CO_3^{2-}$]) to inform primary extinction selectivity in the secondary extinction cascade models. Respiratory protein data for each taxonomic group was gathered from the literature[34-37]. Physiological buffering of each taxonomic group was assigned according to the division given by Bambach et al.[38] and Knoll et al.[39]. The assigned ecological traits of each species are given in the datasets in the Supplementary Material 2. The unique ecological trait combinations within each clade are given in the Table S1.

#### Body size data

Size data were collected from the literature, either from the measurements given in the publication or by measuring from the published drawings/photographs or by digitizing individual datapoints from the scatter plots. A total of 7,205 unique specimen size measurements were used to calculate the mean size for each of the 1,526 species in all datasets (Table S2). The measurements of the same specimens used in multiple datasets, hence the total number of measurements in all datasets, totals 12,848 measurements. Most of the duplicate measurements are for the Foraminifera (48% of the duplicates) and Bivalvia (48%

of the duplicates), mainly *Claraia* and *Unionites*. The references used to compile the size datasets are given in the Supplementary Material 1. Specific references for each size measurement are given in a separate column in the datasets in the Supplementary Material 2.

We used linear measurements, i.e. height and length, as proxies for size as they generally represent the primary axes of body size. As the measurement terminology differs among clades, the height and length variables used in this study for each clade are illustrated in Figure S8.

In some groups (e.g. those with radial symmetry i.e. Ophiuroidea and Anthozoa; as well as some ichnofossils), a single measurement is used as both height and length. Body size categories were assigned using species mean size, calculated as $\sqrt{(\text{height} \times \text{length})}$ for each specimen. Species were classified as tiny (<1 mm), small (1–10 mm), medium (10–100 mm), large (100–1000 mm), or huge (>1000 mm). Shell size was used as a proxy for body size in molluscs, arthropods, foraminifera, and ostracods, burrow diameter for ichnofossils, and individual polyp or zooid size for colonial taxa (bryozoans, corals). For articulated groups (e.g., vertebrates and echinoderms), size estimates were based on disarticulated elements (e.g., conodonts, teeth) following the approach described in the Supplementary Material 1.

#### Paleo Food web Inference Model (PFIM)

We used the Paleo Food web Inference Model (PFIM[40,41]) to reconstruct marine metacommunities. Food webs were modelled based on four ecological trait categories (i.e. feeding, motility, tiering, body size) and feeding rules that dictate whether an organism is able to consume another organism (Figure S9). We reconstructed feasible metawebs (all potential interactions) for all communities across three intervals at seven sites. After reconstructing all potential interactions, we excluded cannibalistic interactions from the reconstructed webs, as they produce unrealistic trophic web metric values, and decrease the trophic level of cannibalistic species.

The food webs are reconstructed at the species-level, and we added one primary producer node and a zooplankton node. All primary consumers have a single feeding link (edge) with the primary producer node. As generality (standard deviation normalized in-degree), vulnerability (standard deviation normalized out-degree) and motifs (apparent and direct competition, omnivory, linear chain) can be biased by changes in the relative proportion of the primary consumers through time, we reconstructed the food webs both with (Figures 1–4) and without a primary node (Figure S11). The R scripts used for the PFIM can be found in the Supplementary Material 2. Species compositions in each time bin at each locality and the trophic levels estimated by PFIM are given as html files in the Supplementary Material 2.

#### Extinction selectivity analysis

Extinction cascades were simulated by subjecting species to primary extinction scenarios based on ecological and trophic traits that correspond to known sensitivities to environmental changes (e.g. high temperatures, anoxia) that have been hypothesised as plausible drivers of mass extinction across the PTME. Specifically, we explored 13 different extinction scenarios. We have simulated the primary extinction order under the following scenarios: 1) random, 2) body size (large to small/small to large), 3) tiering (infaunal to pelagic/pelagic to infaunal), 4) motility (fast to non-motile/non-motile to fast), 5) physiology (buffered to unbuffered/unbuffered to buffered), 6) feeding (carnivorous to herbivorous/herbivorous to carnivorous), 7) respiratory capacity (low to high/high to low), 8) generality (low to high/high to low), and 9) vulnerability (low to high/high to low). Generality and vulnerability of taxa were estimated based on the food webs reconstructed with the PFIM.

For each replicate, we catalogued the primary extinction and any secondary extinctions that arose. Secondary extinctions were treated as being cascading, whereby after the removal of (a) species, all species that had lost all their prey were deemed to have gone extinct and were also removed. This process was repeated until no more species could be removed. Extinctions were stopped when the diversity of the simulated post-extinction community reached the richness that equalled or less than that of the empirical post-extinction community. We generated 50 replicates for each scenario by sampling randomly among species from within each traits' levels in the sequence. Simulated post-extinction food webs were then compared to the empirical post-extinction community using the following approaches.

First, we compared four structural metrics (Connectance, Network generality, Network vulnerability, Maximum trophic level) as well as the frequency of four motifs (Linear chains, Omnivory, Apparent competition, Direct competition) between the empirical and simulated networks. This was done by using the mean absolute difference between the empirical and simulated networks. We then applied z-scoring to mean absolute differences to scale metric values, in order to prevent a single metric from being weighted disproportionately. A value of zero would indicate that the simulated network perfectly matches the network metrics of the empirical network.

Additionally, we also calculated the True Skill Statistic[42] to compare the node level similarities of identity between the empirical and simulated networks:

$$\mathrm{TSS} = \frac{\mathrm{TP} * \mathrm{TN} - \mathrm{FP} * \mathrm{FN}}{(\mathrm{TP} + \mathrm{FN}) * (\mathrm{FP} + \mathrm{TN})}$$

True Positives (TP): Correctly predicted presences (simulated survivors in empirical survivors)
True Negatives (TN): Correctly predicted absence (simulated extinctions in empirical extinction)
False Positives (FP): Incorrectly predicted presences (empirical extinctions in simulated survivors)
False Negatives (FN): Incorrectly predicted absences (simulated extinctions in empirical survivors)

The TSS can range between -1 and 1, where one indicates a perfect match between empirical and simulated network compositions, and anything below zero indicates a performance no better than random.

TSS calculations consider only the identity of species (i.e., has the extinction scenario removed/retained the species known to survive into empirical post-extinction community). We calculated the number of extinct species as the difference between the number of boundary crossers (survivors) and the pre-extinction diversity. We then used this value as the simulated extinction threshold and compared the simulated survivors with the empirical survivors – this means that the TSS calculations will only reflect the removal or retainment of the 'wrong species'. If there is a complete extinction, TSS scores cannot be calculated. We combined the inference from these comparisons to identify the most plausible set of extinction scenarios that could deliver a community that most closely resembles the post-extinction community. The R scripts used for the extinction selectivity analyses can be found in the Supplementary Material 2.

#### Robustness simulations

Robustness ($R_x$) was calculated for each network using the framework introduced in Jonsson et al. (2015) where robustness is defined as the proportion of primary extinctions that will result in x% of all species in the network going extinct (from both primary and secondary cascading extinctions). Here, we estimate $R_x$ for x in 1% to 99% in steps of 1%. This was repeated 500 times using a random primary extinction sequence, this was then used to calculate the mean robustness at each x% value.

A low value of $R_x$ for a particular sequence (relative to other sequences) implies a lower robustness as only a small proportion of primary extinction will lead to an x% collapse of the trophic network. An $R_x$ value of x/100 indicates that no secondary extinctions have occurred and 1/S shows that only one primary species deletion is needed to cause an x% collapse of the network. Robustness simulation scripts written in the Julia language can be found in the Supplementary Material 2.

#### EcoGEnIE

EcoGEnIE is a is 3D ocean circulation model that is an extension of widely used cGEnIE (carbon-centric Grid ENabled Integrated Earth system model) with an ecological component (ECOGEM[43]). Unlike cGEnIE, EcoGEnIE incorporates plankton populations and plankton ecological dynamics by explicitly accounting for the growth and interaction of an arbitrary number of plankton species[43]. EcoGEnIE is composed of three models. The physical model (extension of c-goldstein[44]) is a 3D ocean circulation model coupled with energy moisture balance model and a dynamic thermodynamic sea-ice model[45]. The atmospheric chemistry model (atchem) is a 2D 36×36 atmospheric grid storing atmospheric composition[45]. The biogeochemistry model (biogem) redistributes the essential biogeochemical compounds and isotopes via biological processes[45].

The ocean circulation model calculates the horizontal and vertical transport of heat, salinity, and biogeochemical tracers via the combined parameterisation for isoneutral diffusion and eddy-induced advection[43,44]. The ocean model is configured on a 36x36 equal-area horizontal grid with 16 logarithmically spaced depth levels. The horizontal grid is uniform in longitude (10°) and varying in latitudes (from 3.2° at the Equator to 19.2° near the poles). The thickness of the vertical grid increases with depth, from 40.4 m at the surface to as much as 710.2 m near sea floor.

In the EcoGEnIE, the inorganic nutrients are assimilated by the plankton groups and exported as DOM and POM. The remineralization of POM with depth is predicted by globally fixed exponential function or variable temperature-dependent function[46]. The EcoGEnIE resolves plankton populations by functional groups (e.g., phytoplankton, zooplankton) and organism size (spherical diameter)[47]. In the model, the biological productivity is controlled by a single nutrient (phosphate), and the organic matter production is limited by light and nutrient availability, but biomass of different plankton groups is subjected to resource competition and grazing by zooplankton[45,47]. The nutrient uptake by phytoplankton is governed by a function that is dependent on plankton size, with the smaller plankton have higher nutrient uptake than larger ones[47]. The carbon assimilation is also dependent on size, but with photosynthesis rate decreasing towards largest and smallest size cohorts[47]. Additionally, the phytoplankton growth depends on temperature. The zooplankton grazing depends on available prey biomass and size-dependent maximum grazing rate, and the grazing rate is temperature dependent[47]. The availability of prey biomass is determined as a log-normal function of zooplankton-to-phytoplankton size ratios, with zooplankton predominantly grazing on prey that are 10 times smaller than themselves[47]. The production of organic matter is a function of the mortality of all plankton (linearly related to biomass) and the inefficient assimilation of biomass by zooplankton grazing (at least 30% is lost to organic matter as messy feeding)[47].

To model the carbon climate and plankton biomass, we use the ECOGEM with 8 phytoplankton and 8 zooplankton size categories and the temperature-independent POM. To simulate the Changhsingian pre-extinction interval climate, we used two times the $pCO_2$. To simulate the latest Changhsingian to Griesbachian (extinction interval) and the Griesbachian (post-extinction interval) we used ten times the $pCO_2$ following previous geochemical studies[48,49]. Previously, Hülse et al.[46] and Song et al.[50] simulated Griesbachian climate by doubling the $PO_4$, but there is no geochemical evidence for such increase[51]; hence we kept the $PO_4$ unchanged throughout the Permian–Triassic in our model. The EcoGEnIE output was illustrated using Panoply (NASA Goddard Institute for Space Studies) and MATLAB. EcoGENIE uses a topographic model with coarser grid system, hence the paleocoordinates of locations extracted in R were adjusted to extract environmental variables in EcoGENIE. The EcoGEnIE files used to run the model, the output files (including 2D biogem, 3D biogem, and ecogem netCDF files), and the MATLAB scripts used to generate the figures (3_PermTrias_plots.m, 4_make_extractlocations_pre.m, and 5_make_extractlocations_post.m) can be found in the Supplementary Material 2.

#### Paleogeographic map and paleolatitude inference

The paleogeographic topography map (Figure 1) is based on the PALEOMAP paleodigital elevation models (PaleoDEMS)[52] and is created using the chronosphere[53] R package. Paleolatitude estimates are done with rgplates R package using the PALEOMAP plate model[54]. The R scripts can be found in the Supplementary Material 2.
