## Supplementary material for "No global collapse of food webs across the Permian–Triassic Mass Extinction": Karapunar_et_al_PT food web_Supplementary_Material_1_Data_and_Results

### Metacommunity Datasets

**Taxonomic vetting**

An extensive literature review was conducted to capture all previously reported species from the formations within the target locations/regions. The same species are commonly reported under different names in publications, either due to species misidentifications (heterochresonym), or usage of different species names for the same species (synonyms), or varying taxonomic opinions on generic classifications (combinations). To overcome multiple inclusion of the same species reported under different names (artificial inflation of diversity), synonyms, chresonyms and combination of species names were checked and accounted for by using the last published taxonomic opinion in each instance. In some cases, we revised the generic attribution of taxa (e.g., *Spirorbis* to *Microconchus*) but we refrained from revision of species misidentifications in the latest published opinions. We included all open nomenclature names from the latest taxonomic work on any given systematic group (e.g., Prinoth & Posenato 2023 for Bivalvia from the Dolomites). We included all taxa identified to species level. Fossil occurrences reported in open nomenclature were evaluated to avoid inflating diversity. Indeterminate taxa were counted as distinct species only when no species-level identification existed for the same taxonomic group (genus or higher; e.g., Neoselachii indet.; Perleididae gen. et sp. indet. in Meishan by Wang et al. 2007). When multiple indeterminate names for a given group appeared across publications, we retained a single genus-level record and excluded additional ones; however, multiple indeterminate records reported within the same publication were each treated as distinct species. Nomina nuda (names lacking description or figures e.g., *Dicosmos nana* by Caneva 1906) were excluded.

#### Stratigraphic and palaeoenvironmental setting of studied regions

**Dolomites, northern Italy**

The Dolomites dataset was compiled from 81 publications. Data were collected mostly from the Dolomites (Trentino-South Tyrol), but also from the Carnic Alps (Friuli-Venezia Giulia; 29 out of 419 occurrences). Our analyses only focus on the pre-extinction to immediate post-extinction, only species from the Changhsingian and Griesbachian were included and thus, species occurrences that were not determined at substage level were excluded (i.e., Induan, Early Triassic).

The Changhsingian Bellerophon Formation is regarded as the pre-extinction interval. Following Posenato (2019), the latest Changhsingian to Griesbachian Tesero Member was regarded to be deposited during the mass extinction interval. The post-extinction fauna includes the species from the Griesbachian Mazzin and lower Siusi members of the Werfen Formation. Airaghi's (1907) material was excluded because his specimens, assigned to Permian species, actually derive from the Lower Triassic Werfen

Formation, and Tommasi's (1895) study was excluded because its broad Early Triassic attribution lacks the necessary stratigraphic resolution.

The Dolomites dataset was derived from a palaeolatitude of 5°N. The fossils of the pre-extinction Bellerophon Formation are derived from shallow-shelf carbonates deposited in marginal and restricted low-energy marine environments to shore-face/foreshore open shallow sea near the fair-weather wave base (Farabegoli et al. 2007; Posenato 2010). The mass extinction interval of the Changhsingian to Griesbachian Tesero Oolite was deposited in a shallow water above fair wave base (Farabegoli et al. 2007). The post-extinction Griesbachian marlstones and siltstones of the Mazzin Member, Andraz Member, and lower Siusi Member (late Griesbachian) range from supra-tidal to deep subtidal deposits (Broglio Loriga et al. 1990; Wignall & Hallam 1992).

<table>
<tr><th colspan="5">Dolomites, Italy</th></tr>
<tr><th>Age</th><th>Formation</th><th>Member</th><th>Zone</th><th>Interval</th></tr>
<tr><td rowspan="5">Griesbachian</td><td rowspan="6">Werfen</td><td>lower Siusi</td><td></td><td rowspan="4">post-extinction interval</td></tr>
<tr><td rowspan="3">Mazzin</td><td>I. isarcica</td></tr>
<tr><td>I. staeschei</td></tr>
<tr><td rowspan="2">H. parvus</td></tr>
<tr><td rowspan="2">Tesero</td><td rowspan="2">mass extinction interval</td></tr>
<tr><td rowspan="4">Changhsingian</td><td>H. changxingensis</td></tr>
<tr><td rowspan="3">Bellerophon</td><td rowspan="2">Bulla</td><td>H. praeparvus</td><td rowspan="3">pre-extinction interval</td></tr>
<tr><td rowspan="2"></td></tr>
<tr><td></td></tr>
</table>

**Figure S1.** Generalized stratigraphy and their assignments to pre-, during and post-extinction intervals in Dolomites, Italy.

**Meishan, China**

The Meishan dataset was compiled from 72 publications. As both the Permian-Triassic boundary and bed numbers differ among the Meishan sections and therefore, different publications, we used the latest framework used by Foster et al. (2024) which also provides the most comprehensive occurrence data of benthic invertebrates from the Meishan sections (Meishan A, B, C, D, E, Z; see Yin et al. 1996 for more details about the sections). Foster et al. (2024) updated the stratigraphic data based on latest stratigraphic knowledge (e.g., mixed bed 1 in Sheng et al. 1984 represents Changhsingian following Yin et al. 1996) and discarded any occurrences with unreliable stratigraphic information. We checked all stratigraphic ranges and occurrences of each taxon again for our study by referring back to the original references, and then we updated these accordingly based on the most recently published records. We used the original dataset used by Foster et al. (2024), which consists of 603 species. We excluded Wuchiapingian and Dienerian species occurrences and 77 taxa not assigned to species level as well as any occurrence that could not be validated back to an original publication. From the remaining dataset, a total of 152 species were either renamed, or synonymized, or reassigned to another genus based on recent taxonomic knowledge. Finally, the dataset was further augmented by adding 102 species from the literature, mostly nektonic taxa, that were missing in the initial compilation. The Meishan dataset compiled by Huang et al. (2023) includes the fish taxa *Lissodus xiushuiensis* Wang et al. 2007a, *Polyacrodus jiangxiensis* Wang et al. 2007a and cf. *Caturus* in Wang et al. 2007b. There are no published records of these fossils from the Meishan sections, Changxing County, Zhejiang Province but from Xiushui County, Jiangxi Province (Wang et al. 2007a, 2007b). Therefore, these taxa were not included in our analyses. The current compilation exceeded the previous compilations made from Meishan sections (Jin et al. 2000: 333 species; Song et al. 2013: 168 species; Huang et al. 2023: 172) and comprises 567 Changhsingian–Griesbachian species.

Following Song et al. (2013) and Foster et al. (2024), the Changhsingian Beds from 4b to 24 are regarded as the pre-extinction interval; the Changhsingian to Griesbachian Beds 25–28 (changxingensis and parvus conodont Zones) are regarded as the mass extinction interval, and the Griesbachian Beds 29 to 60 are regarded as post-extinction interval.

<table>
<tr><th colspan="6">Meishan, China</th></tr>
<tr><th>Age</th><th>Formation</th><th>Bed</th><th>Stage</th><th>Zone</th><th>Interval</th></tr>
<tr><td rowspan="4">Griesbachian</td><td rowspan="5">Yinkeng</td><td rowspan="2">29-60</td><td rowspan="2">39-71</td><td></td><td rowspan="2">post-extinction interval</td></tr>
<tr><td>I. isarcica</td></tr>
<tr><td rowspan="3">25-28</td><td rowspan="3">32-38</td><td>I. staeschei</td><td rowspan="3">mass extinction interval</td></tr>
<tr><td>H. parvus</td></tr>
<tr><td rowspan="3">Changhsingian</td><td>H. changxingensis</td></tr>
<tr><td rowspan="2">Changxing</td><td rowspan="2">4b-24</td><td rowspan="2">5-31</td><td>C. meishanensis</td><td rowspan="2">pre-extinction interval</td></tr>
<tr><td>N. yini</td></tr>
</table>

**Figure S2.** Generalized stratigraphy and their assignments to pre-, during and post-extinction intervals in Meishan, China.

The Meishan dataset was derived from a palaeolatitude of 17°N. The Meishan deposits are interpreted to have been deposited on slope of a carbonate platform (Kexin et al. in Yin et al. 1996; Song et al. 2013). The environment became shallower towards the main extinction horizon (subtidal, below fair-weather wave base) and deepened afterwards into the post-extinction interval (Kexin et al. in Yin et al. 1996; Chen et al. 2015).

**Southwest Türkiye**

The Türkiye dataset was compiled from 16 publications and is derived from different sections in Antalya (Çürük Dağ, Köpük Dağ, Demirtaş and Öznurtepe) and Konya (Taşkent). The data were collected from the Changhsingian beds of the Pamucak Formation (Çürük Dağ), Yüğlük Tepe Limestone (Demirtaş, Öznurtepe), Çekiç Dağı Formation (Taşkent), and the Lower Triassic Kokarkuyu Formation (Çürük Dağ), Sapadere Formation (Demirtaş, Öznurtepe), and the İspatlı Member of the Gevne Formation (Taşkent). Although the uppermost Permian formation and the lowermost Triassic formation are named differently in each section, they have the same facies and similar faunal content, suggesting a connected palaeoenvironmental setting (Karapunar et al. 2025a). At Çürük Dağ, the Changhsingian is 12 m thick (Richoz 2006), and the nodular limestones measured by Marcoux et al. (1986; beginning at sample 180/17) and TK47 in Crasquin-Soleau et al. (2002) also fall within this interval based on Richoz's (2006) age assignments. Units E and F at Köpük Dağ and Demirtaş (Richoz 2006) are likewise treated as Changhsingian. In Taşkent, the Changhsingian is 50 m thick (Altıner et al. 2021). Although Groves et al. (2005) did not assign samples to substages, lithology indicates that samples D20–26 and T14–17 derive from the Changhsingian oolite; we further interpret D27–39 and T18–56 as Griesbachian, D40–51 and T57–92 as Dienerian, T93–97 as Smithian, and T97–123 as Spathian. Additional occurrences follow Karapunar et al. (in prep.).

The Changhsingian nodular limestones (wackestone) are regarded as pre-extinction interval. The Changhsingian transitional oolite bed is regarded as the mass extinction interval following Karapunar et al. (2025a), the alternating earliest Triassic (Griesbachian) stromatolite, microbialite and oolite beds are regarded as post-extinction interval. The overlying Dienerian siliciclastic and oolitic deposits are regarded as early recovery interval. Smithian and Spathian data was also collected and regarded as late recovery intervals. These intervals are not included in the analysis, as they are not the primary focus.

<table>
<tr><th colspan="5">Türkiye</th></tr>
<tr><th>Age</th><th>Formation</th><th>Bed</th><th>Zone</th><th>Interval</th></tr>
<tr><td rowspan="2">Griesbachian</td><td rowspan="3">Kokarkuyu/ Sapadere/ Gevne</td><td rowspan="2">microbialites/ oolites</td><td></td><td rowspan="2">post-extinction interval</td></tr>
<tr><td>H. parvus</td></tr>
<tr><td rowspan="3">Changhsingian</td><td>transitional oolites</td><td>H. praeparvus</td><td>mass extinction interval</td></tr>
<tr><td rowspan="2">Pamucak/ Yüğlük Tepe/ Çekiç Dağı</td><td rowspan="2">wackestone</td><td>Paradagmarita monodi</td><td rowspan="2">pre-extinction interval</td></tr>
<tr><td></td></tr>
</table>

**Figure S3.** Generalized stratigraphy and their assignments to pre-, during and post-extinction intervals in southwest Türkiye.

The Türkiye dataset was derived from a palaeolatitude of 12°S. The pre-extinction wackestones are interpreted to be deposited at a shallow shelf environment in the photic zone, the extinction interval with oolites indicates an agitated, shallow environment, and the post-extinction interval is represented by shallow water oolites and microbialites (Karapunar et al. 2025a), which might have been deposited in episodically higher salinity environment (Heindel et al. 2018).

**South Tibet**

The Tibet dataset was compiled from 25 publications. The data were collected from the Selong (Jin et al. 1996; Shen et al. 2006; Wang et al. 2017), Tulong (Xu et al. 2018), Gongda (Zhang & Jin 1976) and Qubu (Shen et al. 2003) sections, which yield the upper Permian Selong and Qubuerga Formations and the lower Triassic Kangshare and Tulong Formations. The species list was primarily compiled from the literature. Additionally, thin sections used by Wignall & Newton (2003), which are stored at the University of Leeds, were re-studied for additional occurrences. Following Shen et al. (2003) and Pan & Shen (2008), the Changhsingian begins at the base of bed 23 at Qubu, with underlying beds treated as Wuchiapingian. At Selong, the Coral Bed (bed 15; Shen et al. 2000), the Caliche Bed, and the Waagenites beds are considered Changhsingian (Jin et al. 1996; Wang et al. 2017), and the lowest two sampled horizons of Wignall & Newton (2003) are reassigned accordingly. Using the zonation of Shen et al. (2003) and taxon ranges from Xu et al. (2018), only the Nimaluoshenza Bed at Tulong is regarded as Changhsingian. From Gongda, the few corals reported from the upper Selong Group (Zhang & Jin 1976) are likewise treated as Changhsingian.

<table>
<tr><th colspan="5">Tibet</th></tr>
<tr><th>Age</th><th>Formation</th><th>Bed</th><th>Zone</th><th>Interval</th></tr>
<tr><td rowspan="2">Griesbachian</td><td rowspan="3">Tulong/ Kangshare</td><td>Ophiceras bed</td><td>O. tibeticum</td><td>post-extinction interval</td></tr>
<tr><td>Otoceras bed</td><td>H. parvus / O. latilobatum</td><td rowspan="2">mass extinction interval</td></tr>
<tr><td rowspan="3">Changhsingian</td><td>Waagenites bed</td><td>C. meishanensis</td></tr>
<tr><td rowspan="2">Qubuerga/ Selong</td><td>Caliche bed</td><td>M. sheni</td><td rowspan="2">pre-extinction interval</td></tr>
<tr><td>Coral bed</td><td></td></tr>
</table>

**Figure S4.** Generalized stratigraphy and their assignments to pre-, during and post-extinction intervals in Tibet.

Shen et al. (2006) placed the extinction horizon in the Changhsingian, at the top of the Selong Formation and base of Kangshare Formation (base of Waagenites bed). Wignall & Newton (2003) suggested an extinction horizon in the Griesbachian. Therefore, we consider the pre-extinction interval as the Qubuerga and Selong formations, the mass extinction interval is assigned from the top of the Selong Formation to the top of the Otoceras bed of the Tulong and Kangshare formations, and the post-extinction interval as the Ophiceras bed of the Tulong and Kangshare formations.

The Tibet dataset was derived from a palaeolatitude of 53°S. The Selong and Kangshare formations are interpreted to have been deposited in a shallow marine, low to medium energy setting on the continental shelf (Jin et al. 1996; Shen et al. 2003; Wignall & Newton 2003). The Qubuerga Formation is regarded to have been deposited in a high-energy, inner shelf shoal setting (Shen et al. 2003) and the Tulong Formation consists of shale and dolostone (Shen et al. 2006), the latter containing ammonoids, foraminifera, conodonts and *Claraia*, and is regarded to have been deposited on the continental shelf.

**Kashmir, India**

The Kashmir dataset was compiled from 17 publications. The data from the Kashmir come from the collections of the Permian Zewan Formation and Changhsingian–Early Triassic Khunamuh Formation at the Guryul Ravine and Barus Spur sections (Baud & Bhat 2014).

Following the zonation of Lyu et al. (2021) and the correlations given in Nakazawa et al. (1975, table 3), we regard Unit D of the Zewan Formation as the pre-extinction interval, Unit $E_1$ of the Khunamuh Formation (Lamellibranch zone of Diener 1915; 26-8-08 in Middlemiss 1910) as the mass extinction interval, and Units $E_2$ and $E_3$ (excluding Bed 70) of the Khunamuh Formation (parvus, isarcica, carinata Zones) as the post-extinction interval at the Guryul Ravine section. At Barus Spur (also known as the Spur 3 km north of Barus), we regard Unit e as the pre-extinction, Unit $f_1$ as the mass extinction interval, and Unit $f_2$ as the post-extinction interval. The brachiopods reported by Diener (1915) from the Zewan Formation were excluded, as the correlation with sampled horizon with Unit D cannot be made (Nakazawa et al. 1975, table 3, for correlation), but the fossils from "Lamellibranch Zone" were included.

The Kashmir dataset was derived from a palaeolatitude of 42°S. The Zewan Formation is considered to have been deposited in a shallow neritic environment ("offshore mixed clastic carbonate deltaic complex" Baud et al. 1996), and the overlying limestones, turbidites and shales of the Khunamuh Formation are interpreted to have been deposited in a deeper shelf environment (Nakazawa & Kapoor 1981; Algeo et al. 2007).

| Kashmir, India | | | | |
|---|---|---|---|---|
| **Age** | **Formation** | **Bed** | **Zone** | **Interval** |
| Griesbachian | Khunamuh | E3 | S. kummeli<br>C. carinata | post-extinction interval |
| | | E2 | I. isarcica<br>H. parvus | |
| Changhsingian | | E1 | H. praeparvus | mass extinction interval |
| | Zewan | D | | pre-extinction interval |

**Figure S5.** Generalized stratigraphy and their assignments to pre-, during and post-extinction intervals in Kashmir, India.

**East Greenland**

The Greenland dataset was compiled from 37 publications. The data were collected from the Changhsingian Schuchert Dal Formation and the Changhsingian–Dienerian Wordie Creek Formation in East Greenland.

The pre-extinction interval was interpreted as the Schuhert Dal Formation, the mass extinction interval as the Changhsingian part of the Wordie Creek Formation (H. triviale Zone in Bjerager et al. 2006, lower Glyphophiceras Bed in Spath 1935), and the post-extinction interval as the Griesbachian levels of the Wordie Creek Formation, below the *Bukkenites rosenkrantzi* ammonite Zone (*Proptychites* Beds in Spath 1935) following the correlation given by Trümpy (1969) and ages given by Bjerager et al. (2006) and Surlyk et al. (2017). The reworked fossils (e.g., brachiopods, bryozoans) in the Triassic horizons (Spath 1935; Teichert & Kummel 1972) were not included in the dataset.

| Greenland | | | | |
|---|---|---|---|---|
| **Age** | **Formation** | **Bed** | **Zone** | **Interval** |
| Griesbachian | Wordie Creek/<br>Kap Stosch | Vishnuites bed | W. decipiens | post-extinction<br>interval |
| | | Ophiceras bed | O. commune | |
| | | | M. subdemissum | |
| | | upper<br>Glyphophiceras | H. martini | |
| Changhsingian | | lower<br>Glyphophiceras | H. triviale | mass extinction<br>interval |
| | Schuhert Dal | | Paramexicoceras/<br>Changhsingoceras | pre-extinction<br>interval |

**Figure S6.** Generalized stratigraphy and their assignments to pre-, during and post-extinction intervals in Greenland.

The Greenland dataset was derived from a palaeolatitude of 31°N. The Changhsingian Schuchert Dal Formation is considered to have been deposited in a shallow shelf setting (Wignall & Twitchett 2002). The overlying Wordie Creek Formation consists of deep marine shales and mudstones, with sandy and conglomeratic turbidites, and are considered as deep marine basinal deposits (Bjerager et al. 2006; Surlyk et al. 2017).

**Siberia, Russia**

The Russia dataset was compiled from 12 publications. The data was collected from the Wuchiapingian–Changhsingian Imtachan Formation and the Changhsingian–Induan Nekuchan Formation in the Verkhoyansk Region, Russia (Setorym River, Kobyume River, Dyby River; Zakharov 2002; Biakov et al. 2018). Following the zonation of Biakov 2024, the upper Intomodesma evenicum Zone is regarded as Changhsingian, and the Changhsingian–Griesbachian boundary was placed on top of the Otoceras concavum Zone, which corresponds to a NCIE (Zakharov et al. 2014, Biakov et al. 2018).

The pre-extinction interval is regarded as the Imtachan Formation, the mass extinction interval is interpreted as cotemporaneous with the uppermost Changhsingian Otoceras concavum Zone, and the post-extinction interval is the remainder of the Nekuchan Formation that falls above the Otoceras concavum Zone.

<table>
<tr><th colspan="4">Russia</th></tr>
<tr><th>Age</th><th>Formation</th><th>Zone</th><th>Interval</th></tr>
<tr><td>Griesbachian</td><td rowspan="2">Nekuchan</td><td></td><td>post-extinction interval</td></tr>
<tr><td rowspan="3">Changhsingian</td><td>Otoceras concavum</td><td>mass extinction interval</td></tr>
<tr><td rowspan="2">Imtachan</td><td>Intomodesma postevenicum</td><td rowspan="2">pre-extinction interval</td></tr>
<tr><td>Intomodesma evenicum</td></tr>
</table>

**Figure S7.** Generalized stratigraphy and their assignments to pre-, during and post-extinction intervals in Greenland.

The Greenland dataset was derived from a palaeolatitude of 75°N. The Imtachan and Nekuchan formations deposited across the Permian–Triassic transition yield successions of sandstone mudstone alternations with carbonaceous clay nodules in the mudstones (Kutygin et al. 2023). The dominant lithology changes from sandstone to mudstone, and the environment is interpreted to change from a coastal marine environment to a slightly deeper shelf environment.

#### Ecological traits

**Table S1.** The unique ecospace occupancy of species within each clade/group. This table contains only unique ecospace of the species studied herein.

| motility | tiering | feeding | size | respiratory protein | physiological buffering |
|---|---|---|---|---|---|
| **Bivalvia** | | | | | |
| facultative | shallow infaunal | herbivore | medium | hemocyanin | buffered |
| facultative | shallow infaunal | herbivore | medium | hemoglobin | buffered |
| nonmotile attached | epifaunal | herbivore | medium | hemoglobin | buffered |
| nonmotile attached | epifaunal | herbivore | small | hemoglobin | buffered |
| nonmotile unattached | semi infaunal | herbivore | medium | hemoglobin | buffered |
| nonmotile unattached | epifaunal | herbivore | medium | hemoglobin | buffered |
| facultative | deep infaunal | herbivore | medium | hemoglobin | buffered |
| nonmotile attached | semi infaunal | herbivore | medium | hemoglobin | buffered |
| facultative | shallow infaunal | herbivore | small | hemoglobin | buffered |
| facultative | shallow infaunal | herbivore | small | hemocyanin | buffered |
| nonmotile attached | semi infaunal | herbivore | small | hemoglobin | buffered |
| facultative | deep infaunal | herbivore | medium | hemocyanin | buffered |
| nonmotile attached | epifaunal | herbivore | large | hemoglobin | buffered |
| facultative | shallow infaunal | herbivore | large | hemoglobin | buffered |
| **Gastropoda** | | | | | |
| slow moving | epifaunal | carnivore | small | hemocyanin | buffered |
| slow moving | epifaunal | omnivore | medium | hemocyanin | buffered |
| slow moving | epifaunal | omnivore | small | hemocyanin | buffered |
| slow moving | epifaunal | herbivore | medium | hemocyanin | buffered |
| slow moving | epifaunal | herbivore | small | hemocyanin | buffered |
| slow moving | shallow infaunal | herbivore | small | hemocyanin | buffered |
| slow moving | shallow infaunal | herbivore | medium | hemocyanin | buffered |
| slow moving | epifaunal | carnivore | medium | hemocyanin | buffered |
| slow moving | epifaunal | herbivore | tiny | hemocyanin | buffered |
| slow moving | epifaunal | carnivore | tiny | hemocyanin | buffered |
| **Cephalopoda** | | | | | |
| fast moving | benthopelagic | carnivore | medium | hemocyanin | buffered |
| fast moving | benthopelagic | carnivore | large | hemocyanin | buffered |
| **Scaphopoda** | | | | | |
| facultative | shallow infaunal | carnivore | medium | hemocyanin | buffered |

| | | | | | |
|---|---|---|---|---|---|
| slow moving | shallow infaunal | carnivore | small | hemocyanin | buffered |
| **Chondrichthyes** | | | | | |
| fast moving | benthopelagic | carnivore | large | hemoglobin | buffered |
| fast moving | benthopelagic | carnivore | medium | hemoglobin | buffered |
| fast moving | benthopelagic | carnivore | small | hemoglobin | buffered |
| fast moving | nektopelagic | microcarnivore | large | hemoglobin | buffered |
| fast moving | benthopelagic | carnivore | huge | hemoglobin | buffered |
| **Actinopterygii** | | | | | |
| fast moving | benthopelagic | carnivore | medium | hemoglobin | buffered |
| fast moving | benthopelagic | carnivore | large | hemoglobin | buffered |
| fast moving | benthopelagic | microcarnivore | large | hemoglobin | buffered |
| fast moving | benthopelagic | microcarnivore | medium | hemoglobin | buffered |
| fast moving | benthopelagic | omnivore | large | hemoglobin | buffered |
| fast moving | benthopelagic | omnivore | medium | hemoglobin | buffered |
| **Sarcopterygii** | | | | | |
| fast moving | benthopelagic | carnivore | large | hemoglobin | buffered |
| fast moving | benthopelagic | carnivore | medium | hemoglobin | buffered |
| **Conodonta** | | | | | |
| fast moving | nektopelagic | microcarnivore | small | hemoglobin | buffered |
| fast moving | nektopelagic | microcarnivore | medium | hemoglobin | buffered |
| fast moving | nektopelagic | microcarnivore | tiny | hemoglobin | buffered |
| **Amphibia** | | | | | |
| fast moving | benthopelagic | carnivore | large | hemoglobin | buffered |
| **Echinoidea** | | | | | |
| slow moving | epifaunal | omnivore | medium | diffusion | unbuffered |
| slow moving | epifaunal | omnivore | small | diffusion | unbuffered |
| **Ophiuroidea** | | | | | |
| slow moving | epifaunal | herbivore | medium | diffusion | unbuffered |
| **Crinoidea** | | | | | |
| nonmotile attached | epifaunal | herbivore | medium | diffusion | unbuffered |
| **Crustacea** | | | | | |
| fast moving | epifaunal | herbivore | medium | hemocyanin | buffered |
| slow moving | epifaunal | herbivore | tiny | hemocyanin | buffered |
| slow moving | epifaunal | herbivore | small | hemocyanin | buffered |
| nonmotile attached | semi infaunal | herbivore | tiny | hemocyanin | buffered |
| **Trilobita** | | | | | |
| fast moving | epifaunal | carnivore | medium | hemocyanin | unbuffered |
| **Rhynchonelliformea** | | | | | |
| nonmotile attached | epifaunal | herbivore | small | hemerythrin | unbuffered |
| nonmotile attached | epifaunal | herbivore | medium | hemerythrin | unbuffered |
| **Linguliformea** | | | | | |
| nonmotile attached | shallow infaunal | herbivore | small | hemerythrin | buffered |
| nonmotile attached | epifaunal | herbivore | small | hemerythrin | buffered |

| | | | | | |
|---|---|---|---|---|---|
| **Bryozoa** | | | | | |
| nonmotile attached | epifaunal | herbivore | tiny | diffusion | unbuffered |
| **Microconchida** | | | | | |
| nonmotile attached | epifaunal | herbivore | tiny | diffusion | unbuffered |
| nonmotile attached | epifaunal | herbivore | small | diffusion | unbuffered |
| **Annelida** | | | | | |
| nonmotile attached | epifaunal | herbivore | small | hemoglobin | buffered |
| **Porifera** | | | | | |
| nonmotile attached | epifaunal | herbivore | small | diffusion | unbuffered |
| nonmotile attached | epifaunal | herbivore | medium | diffusion | unbuffered |
| **Cnidaria** | | | | | |
| nonmotile attached | epifaunal | microcarnivore | medium | hemerythrin | unbuffered |
| nonmotile attached | epifaunal | microcarnivore | small | hemerythrin | unbuffered |
| nonmotile attached | epifaunal | herbivore | tiny | hemerythrin | unbuffered |
| nonmotile attached | epifaunal | herbivore | small | hemerythrin | unbuffered |
| **Foraminifera** | | | | | |
| nonmotile unattached | epifaunal | herbivore | tiny | diffusion | unbuffered |
| nonmotile unattached | epifaunal | herbivore | small | diffusion | unbuffered |
| nonmotile unattached | epifaunal | herbivore | tiny | diffusion | buffered |
| nonmotile attached | epifaunal | herbivore | tiny | diffusion | unbuffered |
| nonmotile unattached | semi infaunal | herbivore | tiny | diffusion | buffered |
| nonmotile unattached | semi infaunal | herbivore | tiny | diffusion | unbuffered |
| nonmotile unattached | shallow infaunal | herbivore | tiny | diffusion | unbuffered |
| **Annelida ichnofossil** | | | | | |
| slow moving | shallow infaunal | herbivore | small | hemoglobin | buffered |
| facultative | shallow infaunal | herbivore | small | hemoglobin | buffered |
| slow moving | shallow infaunal | herbivore | medium | hemoglobin | buffered |
| facultative | deep infaunal | herbivore | small | hemoglobin | buffered |
| slow moving | deep infaunal | herbivore | medium | hemoglobin | buffered |
| slow moving | shallow infaunal | herbivore | tiny | hemoglobin | buffered |
| facultative | deep infaunal | herbivore | medium | hemoglobin | buffered |
| slow moving | deep infaunal | herbivore | small | hemoglobin | buffered |
| **Crustacea ichnofossil** | | | | | |
| fast moving | deep infaunal | herbivore | medium | hemocyanin | buffered |
| facultative | shallow infaunal | herbivore | small | hemocyanin | buffered |
| fast moving | shallow infaunal | microcarnivore | small | hemocyanin | buffered |
| fast moving | deep infaunal | herbivore | small | hemocyanin | buffered |
| facultative | shallow infaunal | herbivore | tiny | hemocyanin | buffered |

| | | | | | |
|---|---|---|---|---|---|
| facultative | shallow infaunal | herbivore | medium | hemocyanin | buffered |
| fast moving | shallow infaunal | carnivore | small | hemocyanin | buffered |
| facultative | deep infaunal | herbivore | tiny | hemocyanin | buffered |
| **Priapulida ichnofossil** | | | | | |
| slow moving | semi infaunal | herbivore | small | hemerythrin | buffered |
| **Nemertea ichnofossil** | | | | | |
| slow moving | semi infaunal | carnivore | medium | hemoglobin | buffered |
| **Nematoda ichofossil** | | | | | |
| slow moving | epifaunal | herbivore | small | hemoglobin | buffered |
| **Mollusca ichnofossil** | | | | | |
| facultative | shallow infaunal | herbivore | medium | hemocyanin | buffered |

#### Body size data

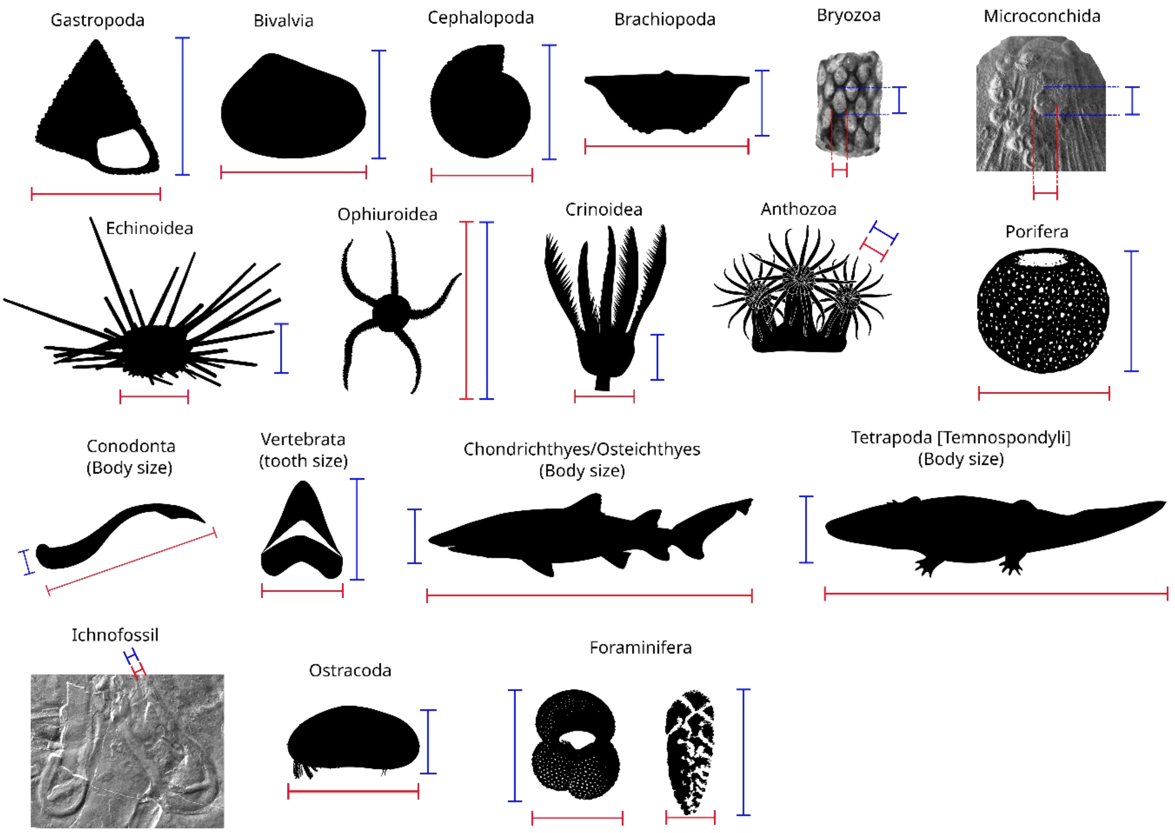


**Figure S8.** Size measurements in different taxonomic groups. Red line indicates length; blue line indicates height measurement. Silhouettes are taken from www.phylopic.org (Katie S. Collins, Tauana J. Cunha, Seth Finnegan, Matthew E. Clapham, Margot Michaud, Aaron O'Dea, DW Bapst, Lauren Sumner-Rooney, F.H. Mollen, R.K. Engelman, Didier Descouens, Nobu Tamura, T. Michael Keesey, Jaime Headden, Joseph P. Botting, Yuandong Zhang, Lucy A. Muir, Maxime Dahirel, Kurtis Wothe, Emily Clark are acknowledged for creating the silhouettes).

The mean size of a species might differ at different locations or through time due to several factors (e.g., high premature mortality, size reduction due to environmental stress such as temperature or dysoxia, sampling bias, reporting bias, preservation bias). To minimize a potential measurement bias, we collected size data of the specimens collected from the studied localities and horizons when possible. In a few cases (e.g., Hofmann et al. 2015), species range in a studied section across multiple horizons, and authors provide an image, but either fail stating from which horizon the specimen was collected, or report it from another horizon (i.e., Dienerian) than the interval of interest (i.e., Changhsingian–Griesbachian). Then we took the measurement for a species from the same locality but from different horizon then the interval of interest. There were also several instances, where authors mentioned a taxon but fail to provide an image or size measurement. In such cases, we took measurement of that species from another publication, although it may be from another location and or stratigraphic horizon. For taxa reported in open nomenclature without any figure or measurement (e.g., *Entalis*? sp. in Caneva 1906; *Zoophycos* isp. in Twitchett 1999), we applied the same procedure and sized using congeneric specimens (e.g., used the genus measurements provided in Monarrez et al. 2021 or the dataset used by Foster et al. 2022, mentioned as Geo.X dataset). In many cases, authors fail to provide the author of the mentioned species (i.e., the person who described the species). In such cases, the author of the species was found with literature research, then the species measurement was taken from other publications. If species identification was uncertain due to homonymy (e.g., *Nodosaria angusta* in Zhao et al. 1978; *Hemigordius regularis* in Song et al. 2009); these records were therefore treated as open nomenclature and sized using congeneric specimens. Such cases are restricted to foraminifera from Meishan and involve taxa with minimal size variation. In a few instances, we reoriented specimens for size measurements, hence our measurements may differ from previously reported sizes but can be reproduced from the published images (e.g., *Aucella* cf. *hausmanni* in Stache, 1878 = *Tambanella*? *stetteneckensis* Prinoth & Posenato, 2023). All data sources and measurement notes are documented in the size dataset.

Possible size biases might be related to collection bias (e.g. preferential collection of larger specimens from surface exposures or limited extraction of smaller fossils due to strong lithification), preservation bias (preservation of certain size groups due to the nature of preservation), reporting bias (preferentially figuring the well-preserved specimens or incomplete reporting of the full size ranges). Size might also be biased due to removal of certain size of species from the population by predation (size selective predation). Although the size-frequency distribution of collected fossils may be biased by sampling, preservation, or predation, we assume that the available measurements approximate the mean body size of each species in life. As the feeding rules allow animals in certain size categories to feed on a size category below their category (e.g., medium is allowed to feed on small), we do not expect size bias due to predation to impact our model. Furthermore, we used size categories (ranges of sizes) rather than absolute measurement, which is expected to mitigate the potential impact of the measurement bias.

**Table S2.** The number of species in each clade/group used in this study, and the number of measured specimens used for size estimates. The entire dataset from 7 sites includes 1,526 unique taxon from the Changhsingian–Griesbachian (and additional 46 taxa from the Dienerian–Spathian from Türkiye) and size measurements of 7,205 specimens.

| Clade/Group | Number of species | Number of measurements |
|---|---|---|
| Foraminifera | 426 | 2428 |
| Crustacea | 273 | 416 |
| Bivalvia | 189 | 2637 |
| Rhynchonelliformea | 156 | 795 |
| Cephalopoda | 155 | 293 |
| Gastropoda | 99 | 171 |
| Conodonta | 93 | 160 |
| Actinopterygii | 32 | 42 |
| Bryozoa | 29 | 34 |
| Cnidaria | 24 | 26 |
| Annelida (ichnofossil) | 19 | 61 |
| Chondrichthyes | 15 | 24 |
| Crustacea (ichnofossil) | 12 | 27 |
| Linguliformea | 7 | 37 |
| Echinoidea | 6 | 9 |
| Ophiuroidea | 6 | 7 |
| Sarcopterygii | 6 | 6 |
| Crinoidea | 4 | 7 |
| Microconchida | 4 | 6 |
| Porifera | 4 | 5 |
| Amphibia | 2 | 2 |
| Ichnofossil | 2 | 3 |
| Scaphopoda | 2 | 2 |
| Trilobita | 2 | 2 |
| Annelida | 1 | 1 |
| Mollusca (ichnofossil) | 1 | 1 |
| Nematoda (ichnofossil) | 1 | 1 |
| Nemertea (ichnofossil) | 1 | 1 |
| Priapulida (ichnofossil) | 1 | 2 |

**Conodonta:** There are examples of conodont fossils with soft-bodied preservation (e.g. Briggs et al. 1983: S-element: 0.76 mm; trunk H: 1.8 mm; L: 40.5 mm; Gabbott et al. 1995: S-element: 10 mm; trunk H: 18 mm; preserved L: 109 mm, estimated L: 400 mm). Herein we assume the ratio of trunk height to conodont element size is assumed as 2.3, body length to tooth element size is assumed as 50.

**Osteichthyans:** For Osteichthyes, if available, the size range of body fossils belonging to the same genus was used as a proxy (e.g., *Bobasatrania*), but in some cases, there are no size measurements for particular genera and only a tooth is available. For Perleididae indet., *Baoqingichthys* and *Zhejiangichthys*, the body height is assumed as 50 times the tooth height, and the body length is assumed 200 times the tooth height, based on scaling in *Teffichthys madagascariensis* (in Marrama et al. 2017). The body size estimates based on these assumptions are within the body size range of *Perleidus* (given in Romano et al. 2016). The body height/length ratio of *Saurichthys* species is estimated as 1/10, based on *Saurichthys spinosa* Wu et al. 2017. The body height/length ratio of *Pteronisculus* species is estimated as 1/5, based on based on the ratio in holotype of *Pteronisculus aldingeri* (Nielsen 1942, pl. 24). The body height/length ratio of *Platysomus* is assumed 1/3. The body height/length ratio of *Australosomus* is interpreted based on scaling in the holotype of *Australosomus kochi* Nielsen, 1949.

**Chondrichthyans**: Because body fossils of Chondrichthyes are rare, body size was estimated using tooth-to-body size ratios derived from taxa with similar tooth morphology, an approach widely applied in the literature (Koot 2013). Owing to the scarcity of articulated specimens and body fossils, such dental-based extrapolations provide the primary basis for size estimation.

Hybodontiformes

The hybodontiform body fossils reported in the literature listed below were used in size estimations.

*Asteracanthus ornatissimus* Agassiz, 1837 (Hybodontiformes, Acrodontidae), Lower Tithonian; tooth W: 25 mm; Body length: 2200 mm (1/88 ratio), body width: 500 mm (1/20) (Stumpf et al. 2021).

*Palaeobates polaris* Stensio, 1921 (Hybodontiformes, Acrodontidae), tooth L: 15 mm estimated body length of ca. 1000 mm (1/67 ratio) (Scheyer et al. 2014).

*Polyacrodus twitchetti* De Blanger, 2005 (=*Polyacrodus claveringensis* Stensiö, 1932) (Hybodontiformes, Acrodontidae), Lower Triassic; tooth L: 1.1 mm, body height 30 mm, body length: 110 mm (preserved), 180 mm estimated (De Blanger 2005).

*Hamiltonichthys mapesi* (Hybodontiformes), Pennsylvanian; tooth length: 6 mm, scale length: 0.45-0.5 mm (1/600 ratio); body length 300 mm (1/50 ratio) body width: 30 mm (1/5) (Maisey 1989).

Based on these Hybodontiformes body fossils, for *Acrodus* sp. and *Polyacrodus* sp., the body length is assumed 50 to 88 times of tooth length, body height is assumed 5 to 20 times of tooth length.

Ctenacanthiformes

The ctenacanthiform body fossils reported in the literature listed below were used in size estimations.

*Dracopristis hoffmanorum* (body length 207 cm, trunk height 50 cm, first dorsal spine 57 cm (27% of body height), tooth height: 2.5 cm, tooth length: 2.5 cm) (tooth height/body length: 1/83 ratio) (body height/length Ratio 1/4 ratio) (Hodnett et al. 2021).

Based on the body proportions of *Dracopristis hoffmanorum*, *Ctenacanthus ishii*, the body length is assumed 83 times of tooth length, body height is assumed 20 times of tooth length.
*Helicoprion* Karpinsky, 1899 estimated body length 1000–1500 mm in the Early Triassic (Scheyer et al. 2014). According to Gayford et al. (2024) and references therein, the body length is approximately 12.5 times the tooth whorl diameter. Herein tooth whorl diameter of *Sinohelicoprion* is assumed to be 6 times the tooth height. The body height of *Sinohelicoprion, Helicampodus*, and *Parahelicampodus* is assumed as 15 times the tooth height. The body length of *Sinohelicoprion* is assumed as 75 times the tooth height.

<u>Dermal scales</u>
We herein assume for chondrichthyan taxa with only dermal scale preservation, the body length is 600 times the dermal scale size, and the body height is 60 times the body length. This assumption has been made based on the scaling relationship between body size and scale size in *Hamiltonichthys mapesi* mentioned above, which represents the only body fossil with documented scale sizes that we know of. However, extant sharks show different scaling between the dermal scale size and body length (e.g., Raschi & Mucisk 1984; Dillon et al. 2017), and the dermal scale size differs along the body (e.g., Ankhelyi et al. 2018). Hence, there is no single formula fitting for all and additional aspects further complicates accurate size estimations.

**Temnospondyls:** The body length of Temnospondyls were assumed to be four times the median skull length. This ratio is assumed based on the reconstruction of *Paracyclotosaurus davidi* in Schoch & Milner (2000, fig. 91).

Our body size estimates for chondrichthyans and other vertebrates are intended only to place taxa into broad size categories by using a consistent assumption, not to provide precise or realistic body lengths. Although we applied linear scaling, true tooth–body allometries are likely non-linear and taxon-specific, and developing robust allometric equations would require dedicated future study. Because PFIM uses discrete size classes rather than continuous measurements, moderate estimation errors have limited impact on our results, except in rare cases where large taxa might be underestimated into smaller categories.

**Echinoderms:** Body size proxies included test diameter in echinoids, cup size in crinoids, arm span in ophiuroids, and body length and height in holothuroids. Because these groups are commonly preserved as disarticulated ossicles, taxa lacking articulated specimens were assigned to the medium size class (10–100 mm), with assumed dimensions of 60 mm for crinoids and ophiuroids and 50 mm for echinoids and holothuroids. Size estimates for *Archaeocidaris ladina* (the Dolomites) and *Archaeocidaris* sp.

(Türkiye) were based on published upper and lower size limits for the genus, using test diameter only and excluding spine length.

**Sponges:** If there were no report of body size of a sponge, for instance if it was described from a thin section, the body size was assigned to a size category. Regarding the predators of the sponges (arthropods, crustaceans, fish) and their size range (small to medium), we have assigned two size categories. Hexactinellid sponges were assigned to be small (1–10 mm). Calcaracea was assigned to medium (10–100 mm).

### Inference of community composition from the fossil record

Inferred community composition from the fossil record is not fully comparable to living communities in a modern ecological sense. This is largely because the fossil record is subject to several filters, such as fossilization, preservation, and sampling. For instance, soft bodied fossils are largely absent in the fossil record, and certain skeletal groups can disappear from the record as their shell is more prone to dissolution due to less stable mineralogy (aragonite).

Our inferences on community composition can also be distorted by sampling. For instance, certain groups might be studied more than others (e.g., foraminifers or conodonts due to their utility in biostratigraphy), or certain layers may be less studied as they contain fewer or poorly preserved fossils, or due to physical difficulties in extracting fossils from them.

The fossil record also presents an additional complexity; time averaging, the co-existence of fossils in the same rock formation even if they may not have co-existed in time. Hence, Fossil communities represent elements of marine communities averaged over fairly long periods of time, that is way beyond the animal lifespan and potentially longer than species duration. Time averaging can also increase the chance of finding more taxa in the rock record, as the longer the time span recorded in a formation, the higher probability of organisms being recorded in the fossil record. Time averaging can be either due to low sedimentation rates or vertical mixing by burrowing animals.

Size sorting or lateral mixing (allochtonous elements) can also cause the composition of a sample to differ from the original community composition, by elimination of species existing in a community or addition of species from adjacent assemblages. As such, a fossil community is more comparable to a modern meta-community, as the rock layers may record different sets of communities that migrate, or are laterally connected.

Modern ecological studies show that food web structure and properties can differ depending on sampling effort (Goldwasser & Roughgarden 1997) and spatial extent (Wood et al. 2015). In addition to the sampling and spatial biases, environmental change during the target time interval (environmental bias such as anoxia) and loss of information due to non-fossilization (preservational bias) can further bias reconstruction of food webs in fossil communities. One way to overcome potential sampling biases is to reconstruct food webs with trophic species (i.e., species that have the identical consumers and resources); however, trophic species may not overcome preservational bias, since soft-bodied taxa rarely fossilise so the tropho-species these organisms represent will still be neglected. Nevertheless, Shaw et al. (2021) showed that selective loss of soft-bodied tropho-species does not result in significantly different food web metric values compared to random loss, but non-fossilization (node loss) can have distinct effects on food web metrics, albeit that are predictable and consistent.

Although inferred fossil communities should not be considered exact counterparts of living communities, they are representative of a living community and can inform us about the temporal change in ecological processes and communities' response, if sampling and fossilization do not differ significantly through time. To decrease the potential impacts of sampling and fossilization biases, we have collected datasets from the best sampled sections across the Permian–Triassic boundary, that i) have a continuous rock record, ii) are fossiliferous both in the late Permian and in the Early Triassic, and iii) do not indicate significant changes in taphonomic settings (i.e., differential preservation of skeletal or soft bodied groups). Hence, the sampling effort or taphonomic settings in the studied sections are not assumed to significantly differ across the Permian–Triassic boundary.

### The Paleo Food Web Inference Model (PFIM)

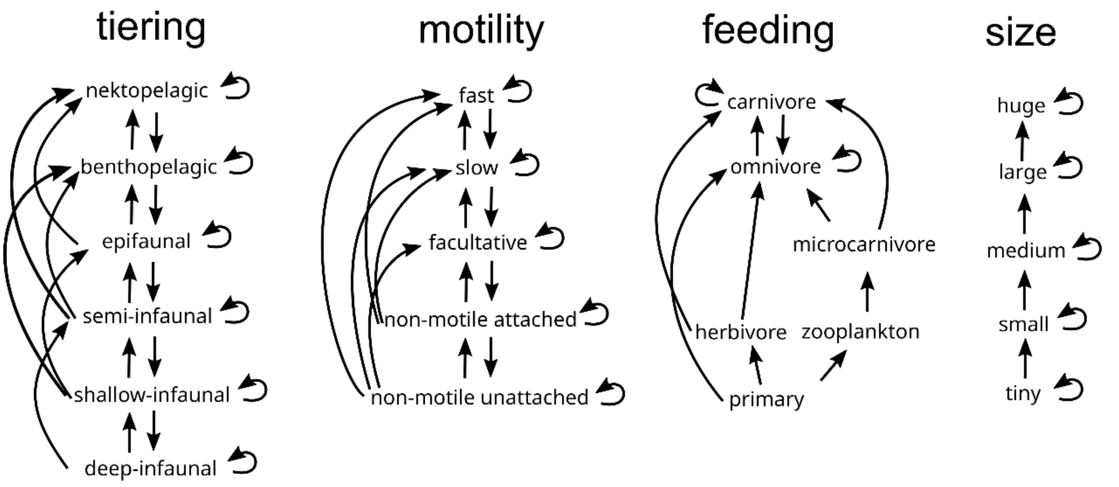


**Figure S9.** Feeding rules to reconstruct food webs using PFIM.

**Table S3.** Food web metrics.

| Metric type | Metric | Description |
|---|---|---|
| Network structure | Connectance | Proportion of possible links in a network that actually occur. Calculated as the number of links (L) divided by number of species (S) squared (i.e. L/S^2). |
| | Size | Number of nodes (species) in a network |
| | Network generality | Standard deviation of normalized in-degree of nodes (see node position metrics below) |
| | Network vulnerability | Standard deviation of normalized out-degree of nodes (see node position metrics below) |
| | Mean trophic level | Mean trophic level of nodes (see node position metrics below) |
| | Maximum trophic level | Maximum trophic level of nodes (see node position metrics below) |
| | Motif: Linear chains | Number of motifs (3-node subnetworks) describing simple linear chains, normalized to S^2. |
| | Motif: Omnivory | Number of motifs describing predators preying on two taxa at different trophic levels, normalized to S^2. |
| | Motif: Apparent competition | Competition between two prey with a common predator. Calculated as the number of motifs describing predators preying on two species, normalized to S^2. |
| | Motif: Direct competition | Competition between two predators with a common prey. Number of motifs describing two predators sharing a prey species, normalized to S^2. |
| Node position | Generality (normalized in-degree) | How many resource nodes a consumer node has. Normalized to the mean number of links per node for that network. |
| | Vulnerability (normalized out-degree) | How many consumer nodes a resource node has. Normalized to the mean number of links per node for that network. |
| | Trophic level | 1 + the weighted mean of the trophic levels of its resources. Primary producers are assumed to have trophic level of 1. |

### Supplementary Results

**Table S4.** The total species diversity, surviving and originating taxa, the extinction and origination percentages in each time-interval.

| interval | location | diversity | survivor from pre-extinction | survivor from extinction interval | originated | extinction percent of pre-extinction taxa | extinction percent of extinction interval taxa | in bin originated percentage |
|---|---|---|---|---|---|---|---|---|
| pre-extinction | Dolomites | 363 | NA | NA | NA | NA | NA | NA |
| extinction interval | Dolomites | 54 | 29 | NA | 25 | 92.01 | NA | 46.30 |
| post-extinction | Dolomites | 68 | 7 | 13 | 61 | 98.07 | 75.93 | 89.71 |
| pre-extinction | Turkiye | 173 | NA | NA | NA | NA | NA | NA |
| extinction interval | Turkiye | 17 | 13 | NA | 4 | 92.49 | NA | 23.53 |
| post-extinction | Turkiye | 79 | 10 | 1 | 69 | 94.22 | 94.12 | 87.34 |
| pre-extinction | Meishan | 460 | NA | NA | NA | NA | NA | NA |
| extinction interval | Meishan | 166 | 87 | NA | 79 | 81.09 | NA | 0.48 |
| post-extinction | Meishan | 70 | 19 | 41 | 51 | 95.87 | 75.30 | 0.73 |
| pre-extinction | Tibet | 129 | NA | NA | NA | NA | NA | NA |
| extinction interval | Tibet | 115 | 41 | NA | 74 | 68.22 | NA | 64.35 |
| post-extinction | Tibet | 45 | 9 | 28 | 36 | 93.02 | 75.65 | 80.00 |
| pre-extinction | Kashmir | 38 | NA | NA | NA | NA | NA | NA |
| extinction interval | Kashmir | 33 | 11 | NA | 22 | 71.05 | NA | 66.67 |
| post-extinction | Kashmir | 66 | 4 | 11 | 62 | 89.47 | 66.67 | 93.94 |
| pre-extinction | Greenland | 24 | NA | NA | NA | NA | NA | NA |
| extinction interval | Greenland | 18 | 4 | NA | 14 | 83.33 | NA | 77.78 |
| post-extinction | Greenland | 95 | 1 | 4 | 94 | 95.83 | 77.78 | 98.95 |
| pre-extinction | Russia | 11 | NA | NA | NA | NA | NA | NA |
| extinction interval | Russia | 23 | 0 | NA | 23 | 100.00 | NA | 100.00 |
| post-extinction | Russia | 28 | 0 | 11 | 28 | 100.00 | 52.17 | 100.00 |

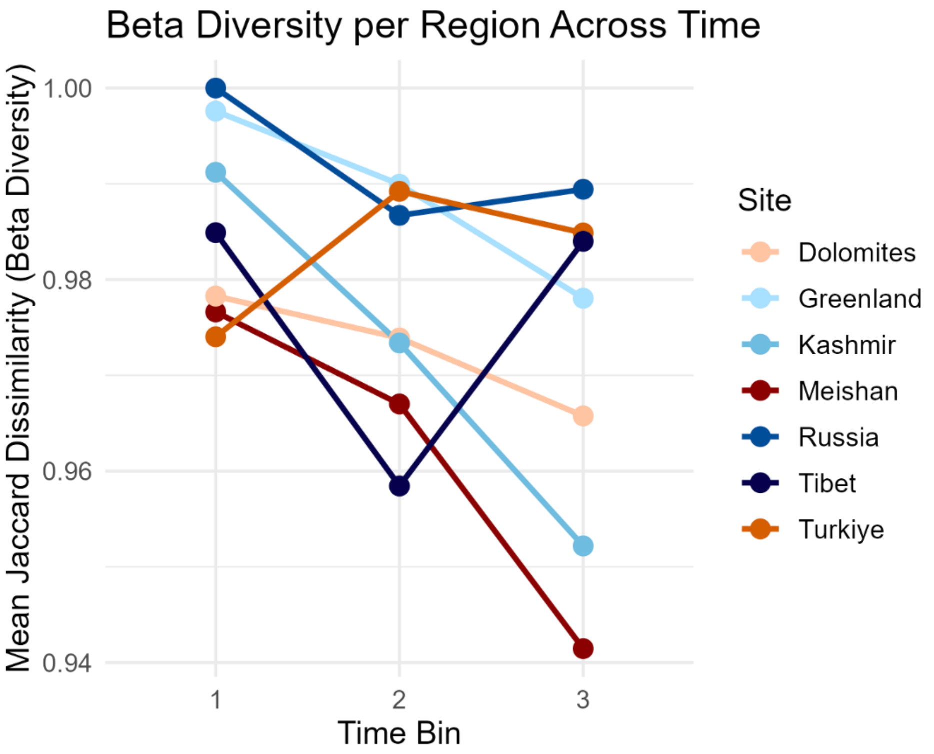


**Figure S10.** Mean beta diversity (Jaccard Dissimilarity) through time. Each data point corresponds to the mean Jaccard dissimilarity of a given location relative to the other six locations in the same time bin.

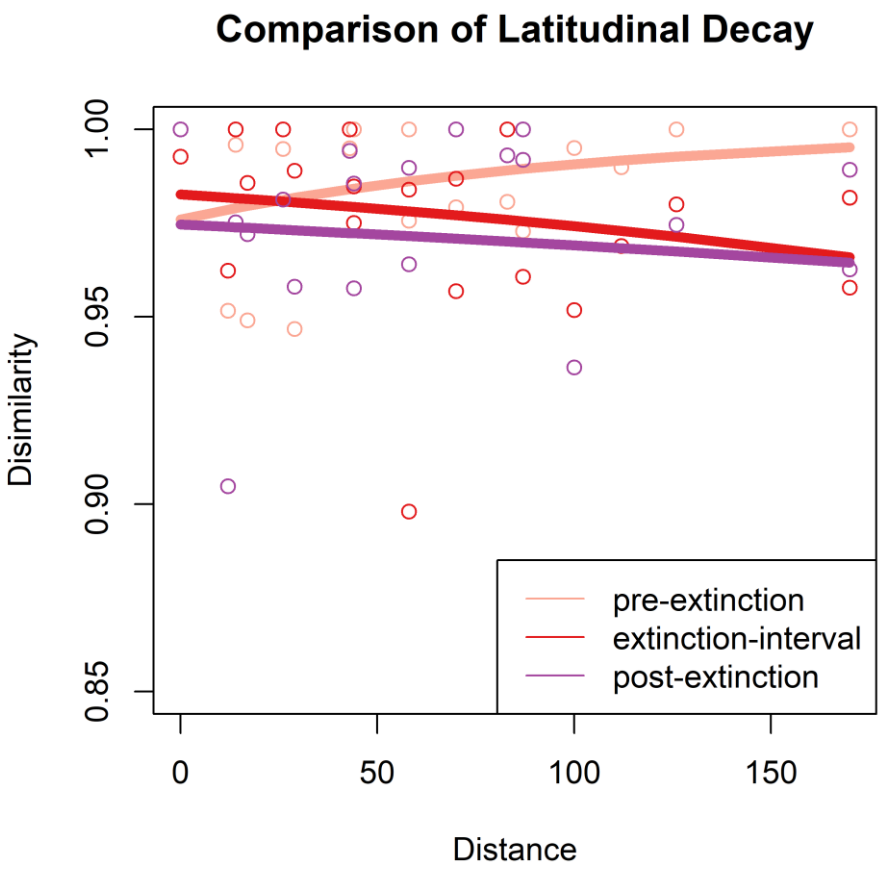


**Figure S11.** Jaccard Dissimilarities with latitudinal distance in three time bins. Distant communities show lower dissimilarity during and after the extinction, due to high cosmopolitanism. Exponential regression models were fitted using decay.model function in betapart R package (Baselga & Orme 2012) which uses Levenberg-Marquardt Nonlinear Least-Squares Algorithm.

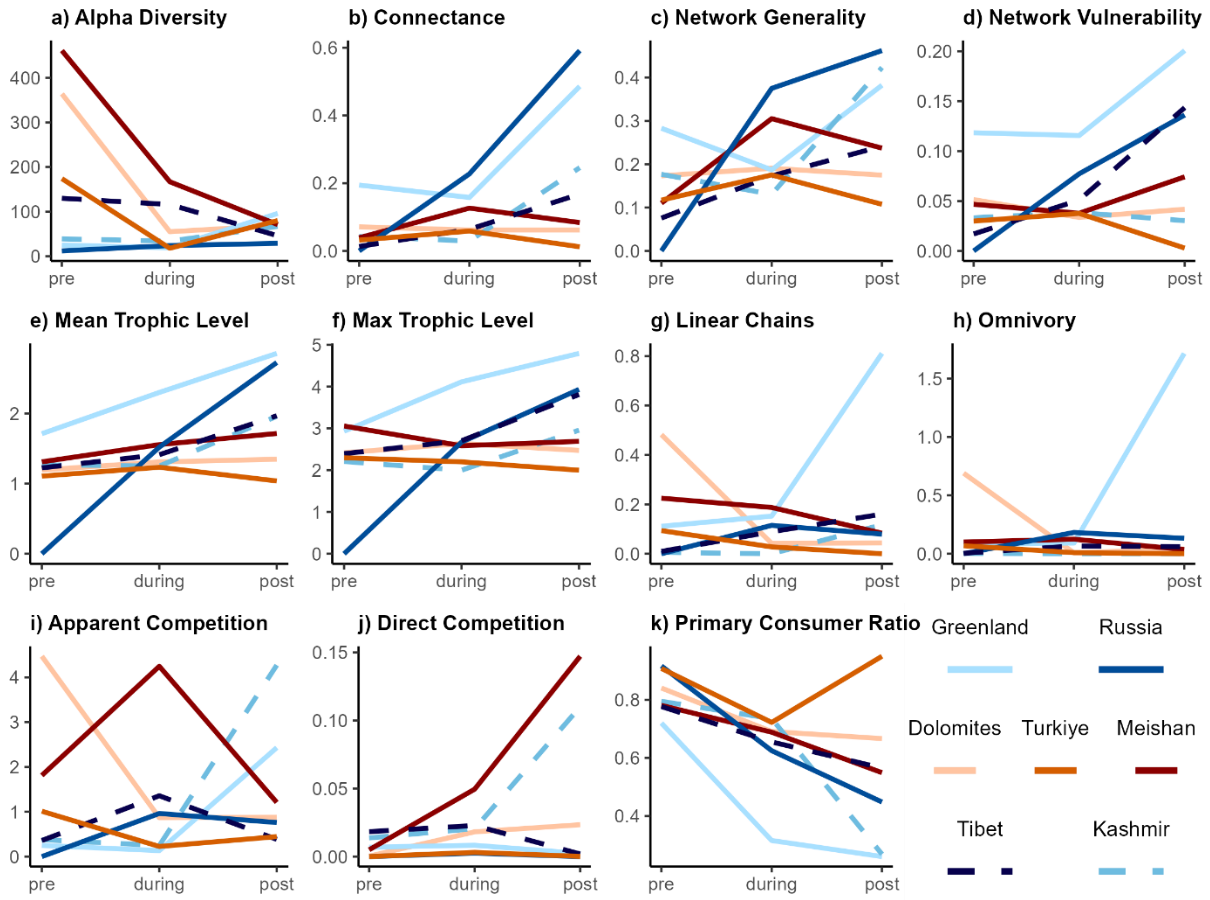


**Figure S12.** Food web metrics of communities across the PTME, <u>remodelled excluding primary producer node</u> to eliminate the effects of temporal decline in primary consumers on the direct competition metric. The model indicates that direct competition among predators increased in Kashmir and low-latitude communities (Meishan, Dolomites, Türkiye) but declined in Tibet and Greenland across the PTME. Low latitudinal sites coloured in red tones, high latitude sites are coloured in blue tones.

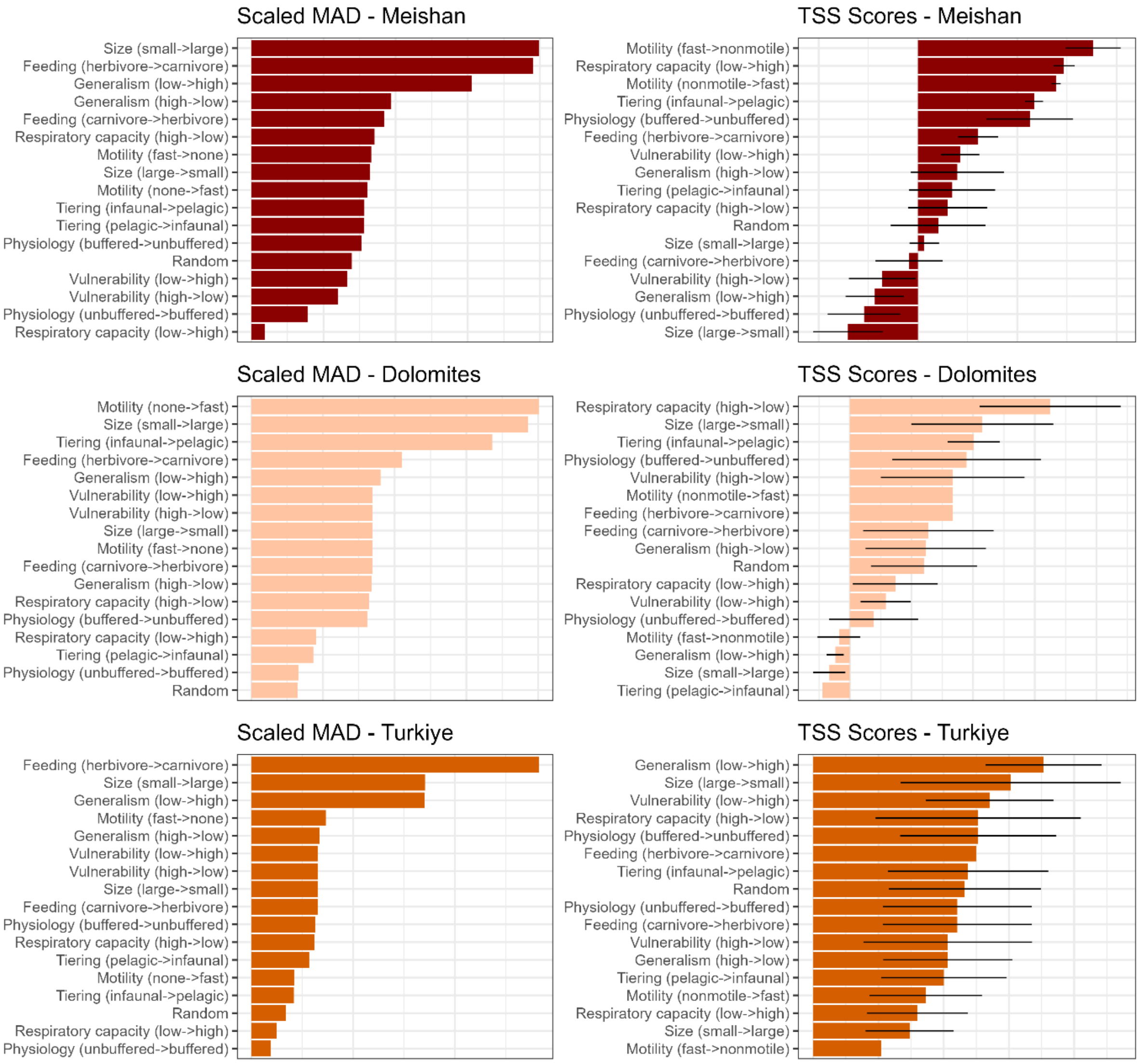


**Figure S13.** Extinction selectivity simulations from pre-extinction interval to mass extinction interval in tropical communities under random scenario and 8 ecological or physiological traits targeting in two directional trait removal.

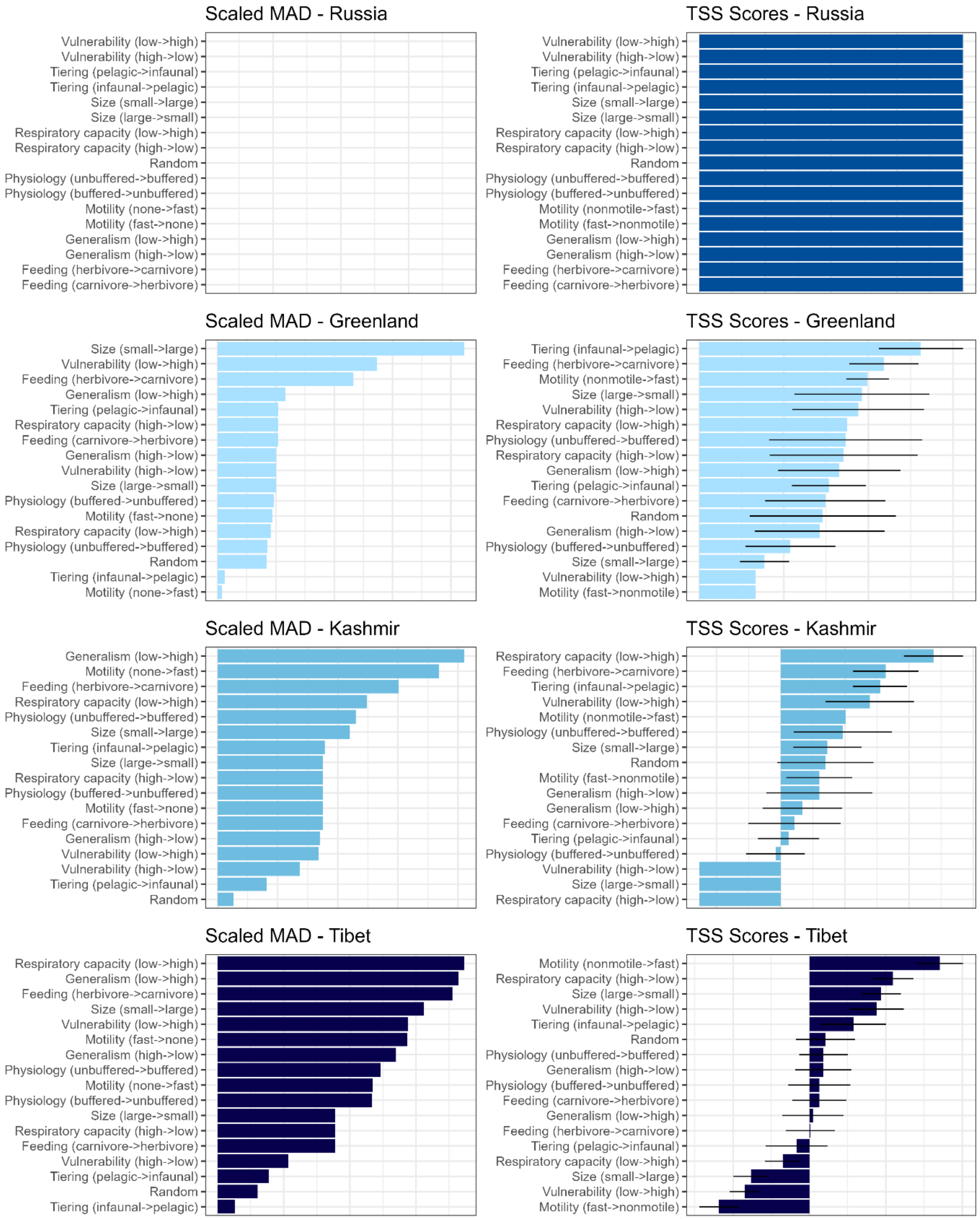


**Figure S14.** Extinction selectivity simulations from pre-extinction interval to mass extinction interval in mid to high latitude communities under random scenario and 8 ecological or physiological traits targeting in two directional trait removal. In Russia, no species survives during the extinction interval, hence different scenarios appear equally likely.

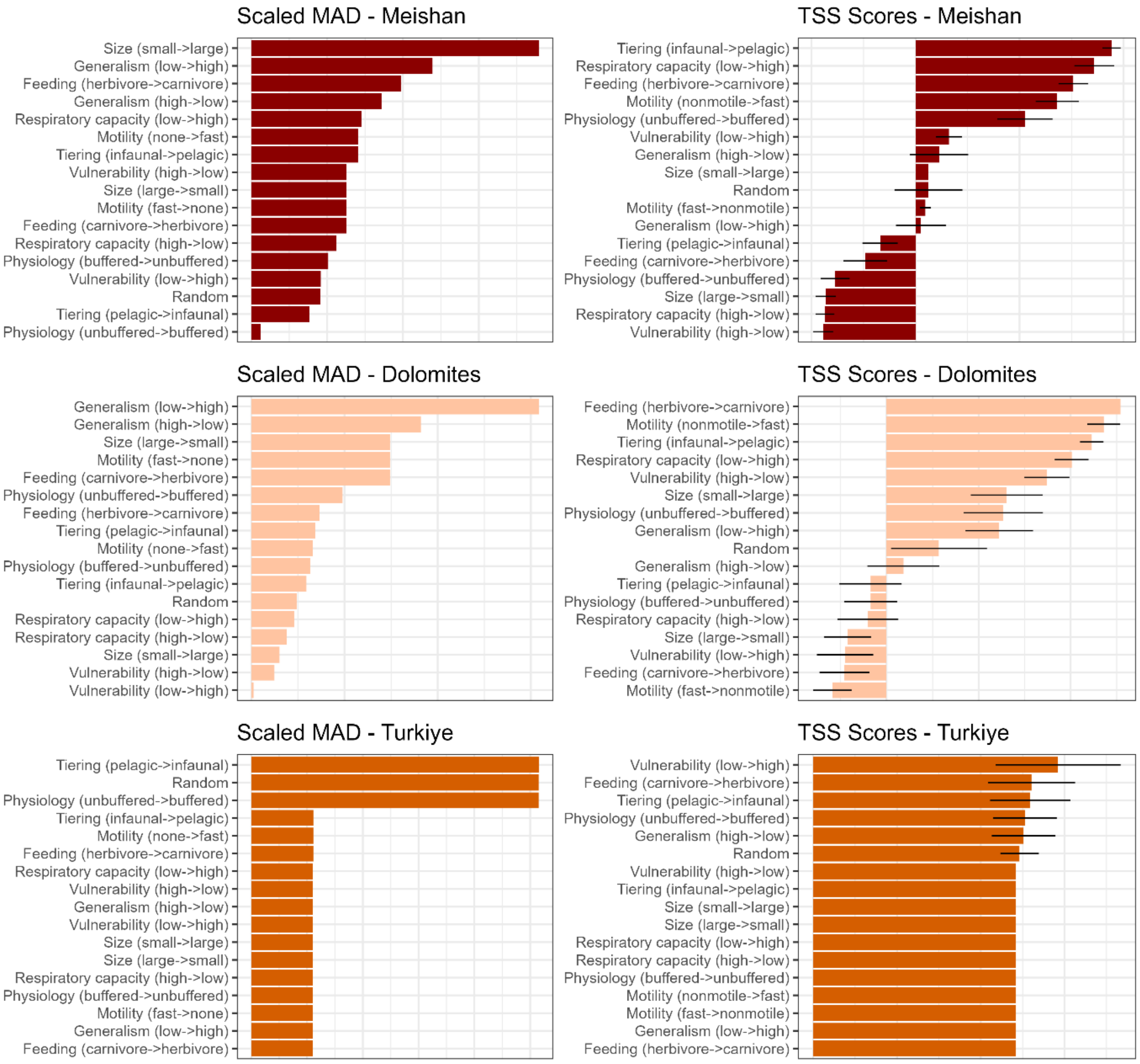


**Figure S15.** Extinction selectivity simulations from mass extinction interval to post-extinction interval in tropical communities under random scenario and 8 ecological or physiological traits targeting in two directional trait removal. In Türkiye, only a single species survives during the extinction interval, hence different scenarios appear equally likely.

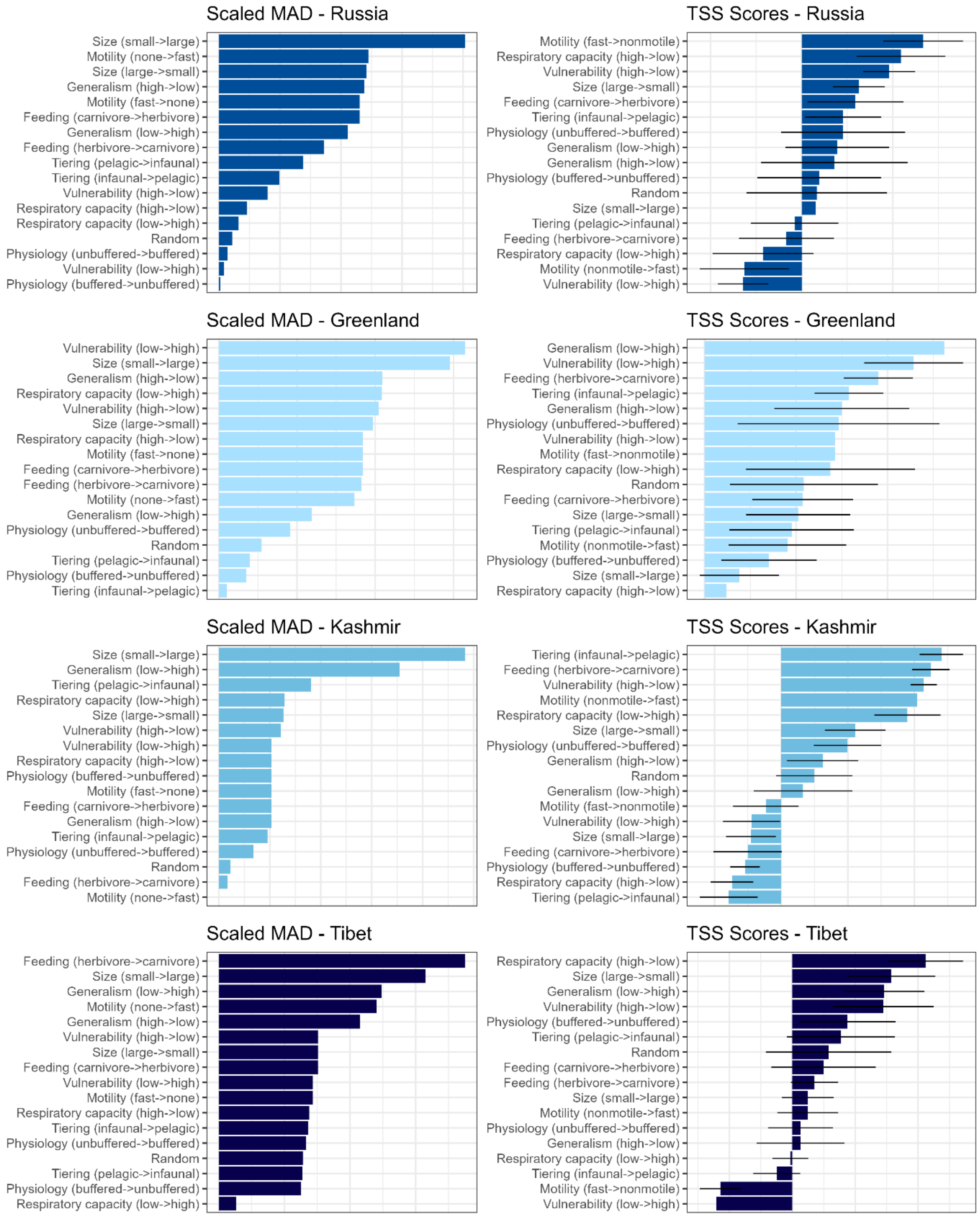


**Figure S16.** Extinction selectivity simulations from mass extinction interval to post-extinction interval in mid to high latitude communities under random scenario and 8 ecological or physiological traits targeting in two directional trait removal.

#### Climate Simulations

Climate simulations with EcoGENIE indicate a global increase in sea surface temperatures (SST) across the PTME, with the greatest rise at mid- to high latitudes (45°–70°) and the smallest at latitudes above 75°NS (Figure 5A). Benthic oceanic temperatures also increased globally, peaking on the continental shelves. pH decreased globally by 0.6–0.7 across all studied regions. Although slight deoxygenation (<0.00006 mol/kg) occurred in both surface and benthic waters, with the highest impact at mid- to high latitudes, surface waters did not reach dysoxic levels (<0.00009 mol/kg), and benthic conditions remained within tolerable limits at all the studied localities. Plankton biomass and primary productivity increased across the PTME in most parts of the Palaeotethys and Neotethys (Dolomites, Türkiye, Tibet, and Kashmir; Figure 5D). The particulate organic carbon (POC) export flux, a proxy for food availability for filter and deposit feeders, declined only in Greenland and Russia for the modelled communities.

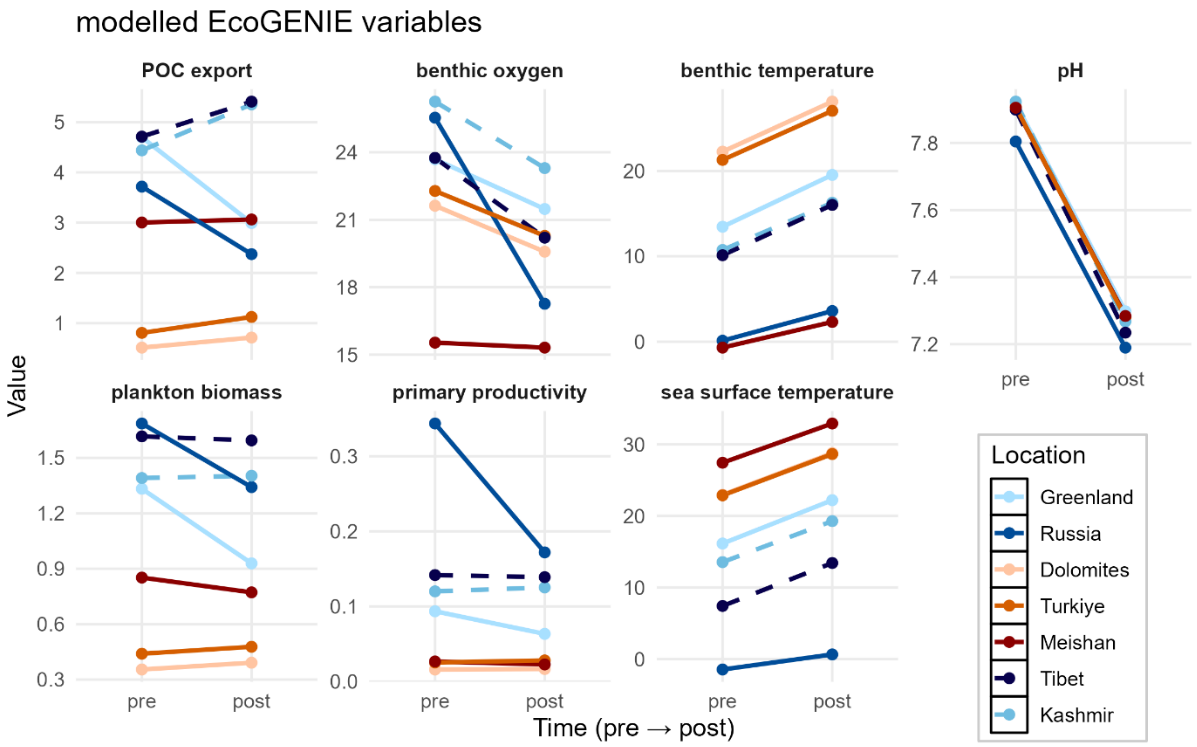


**Figure S17.** Modelled EcoGENIE variables at each location pre- and post-extinction intervals. POC export ($10^{11}$ C mol/year), benthic $O_2$ ($10^{-5}$ mol/kg), benthic temperature (°C), pH (pH units SWS), plankton biomass (total Carbon mmol/$m^3$), primary productivity measured as chlorophyll biomass (mg chl/$m^3$), sea surface temperature (°C).

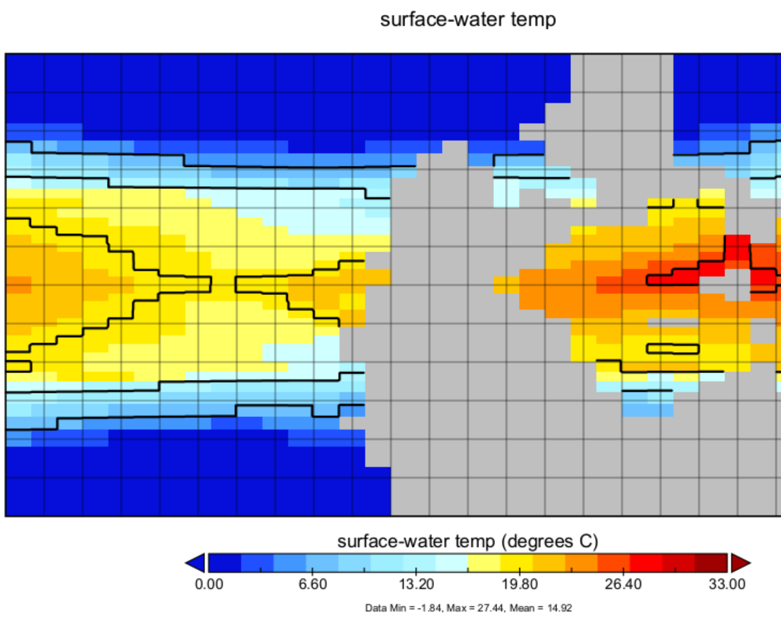


**Figure S18.** SST (x2 pCO2).

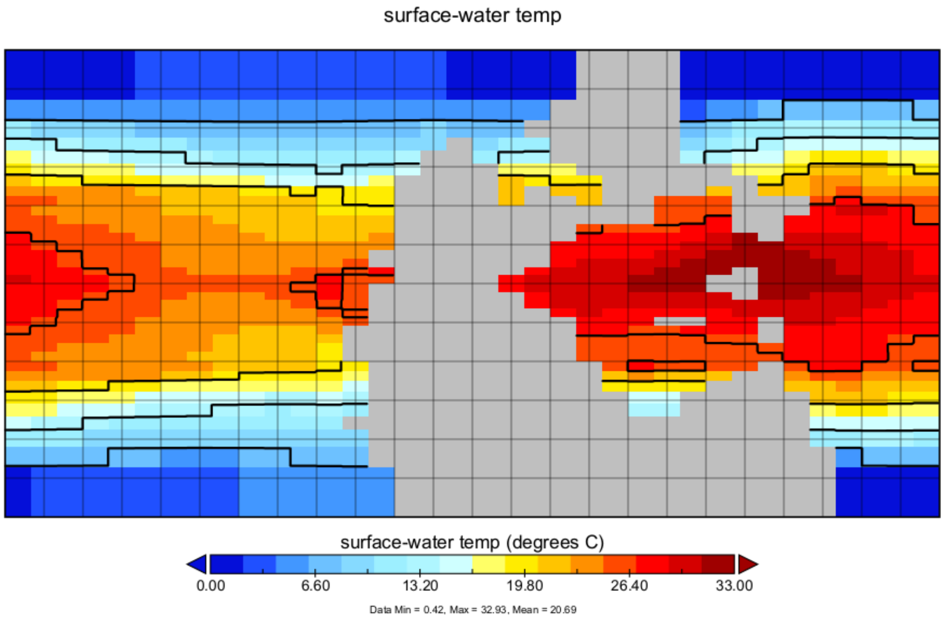


**Figure S19:** SST (x10 pCO2).

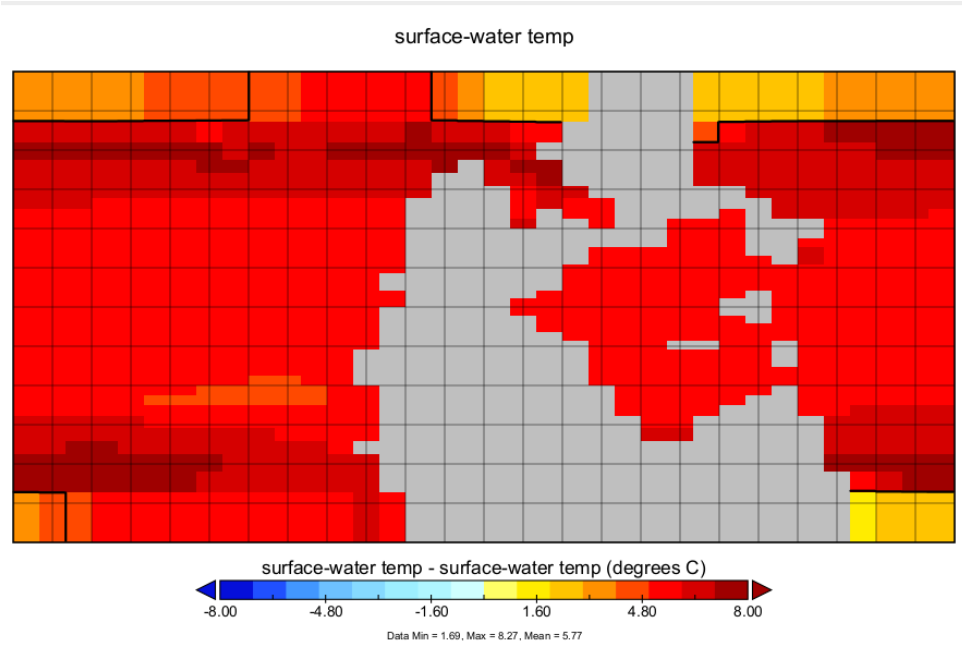


**Figure S20.** SST change (from x2 to x10 pCO2).

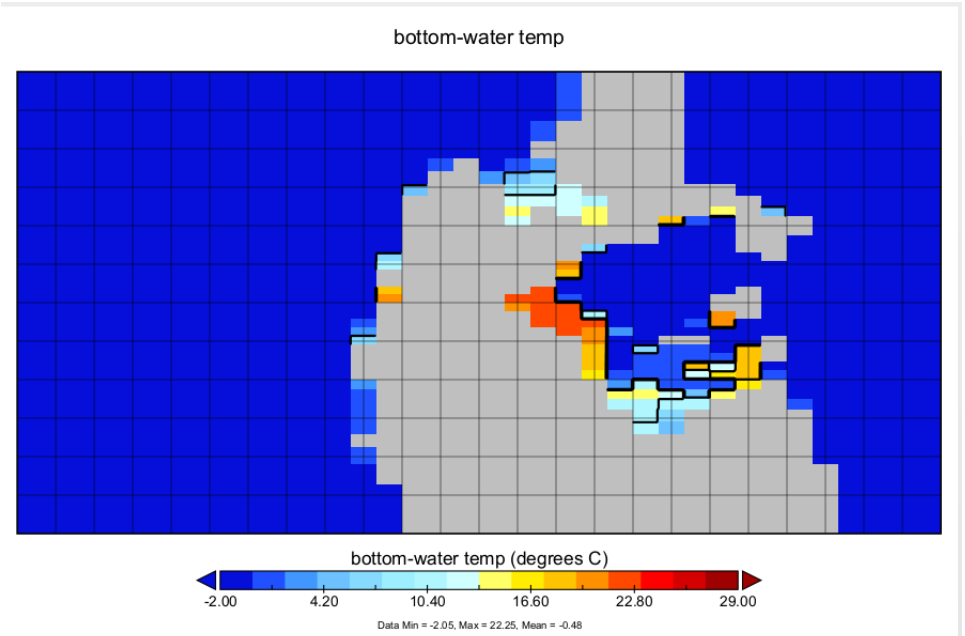


**Figure S21.** Bottom water temperature (x2 pCO2).

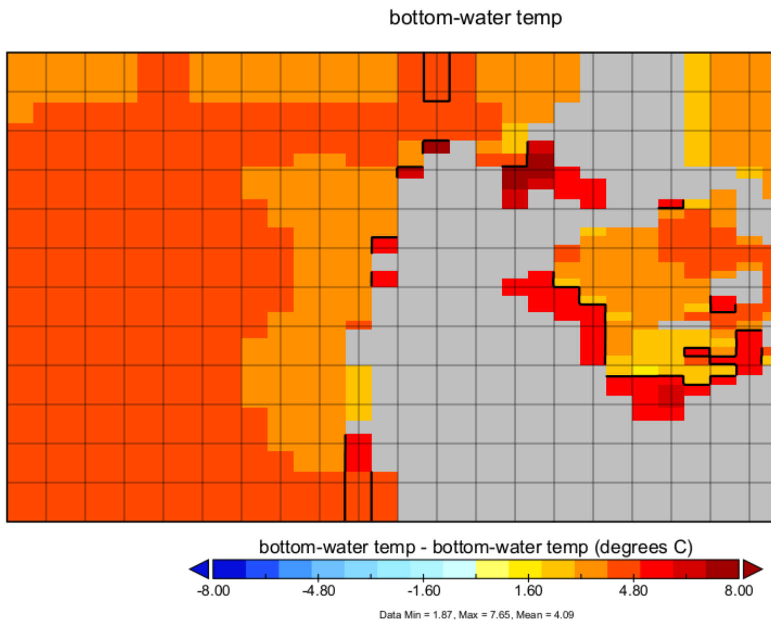


**Figure S22.** Bottom water temperature change (from x2 to 10x pCO2).

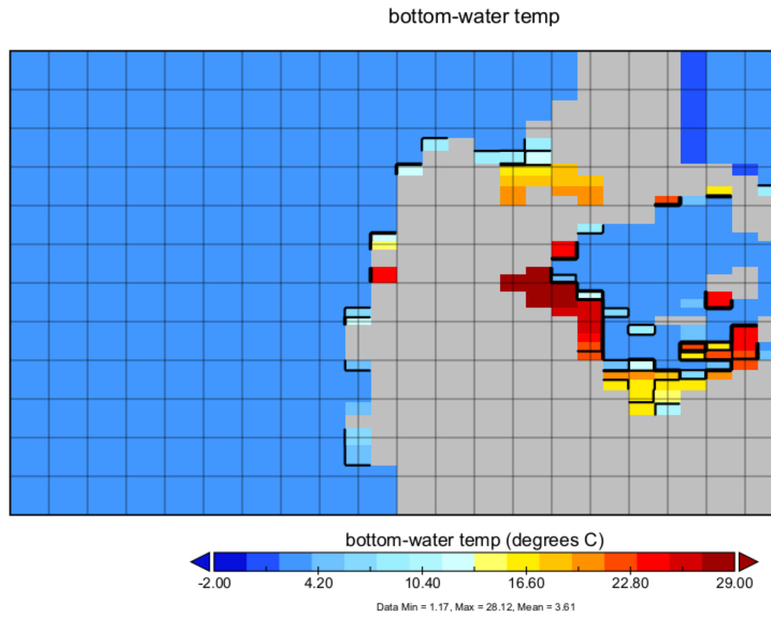


**Figure S23.** Bottom water temperature (x10 pCO2).

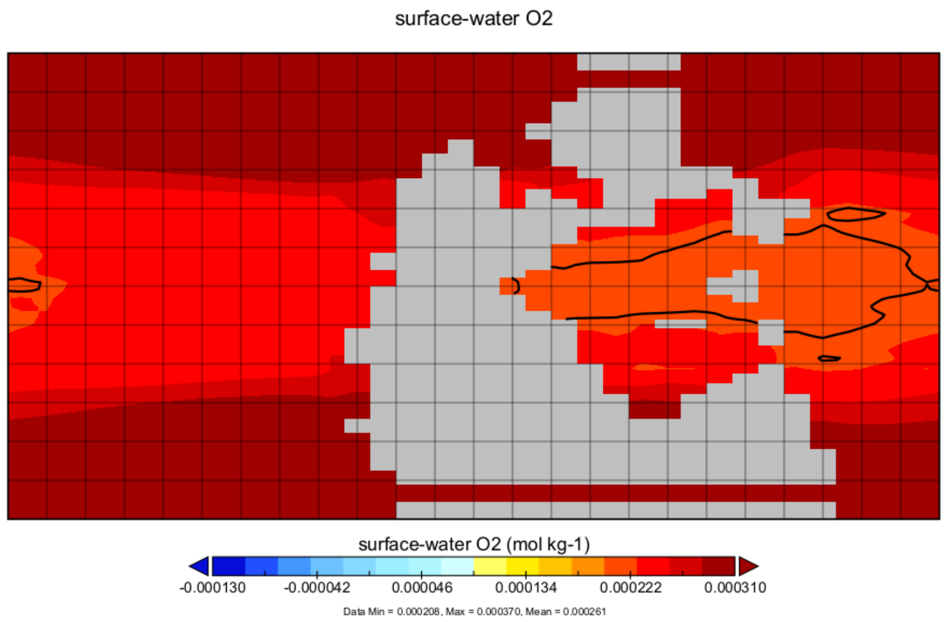


**Figure S24.** Surface water O2 (x2 pCO2).

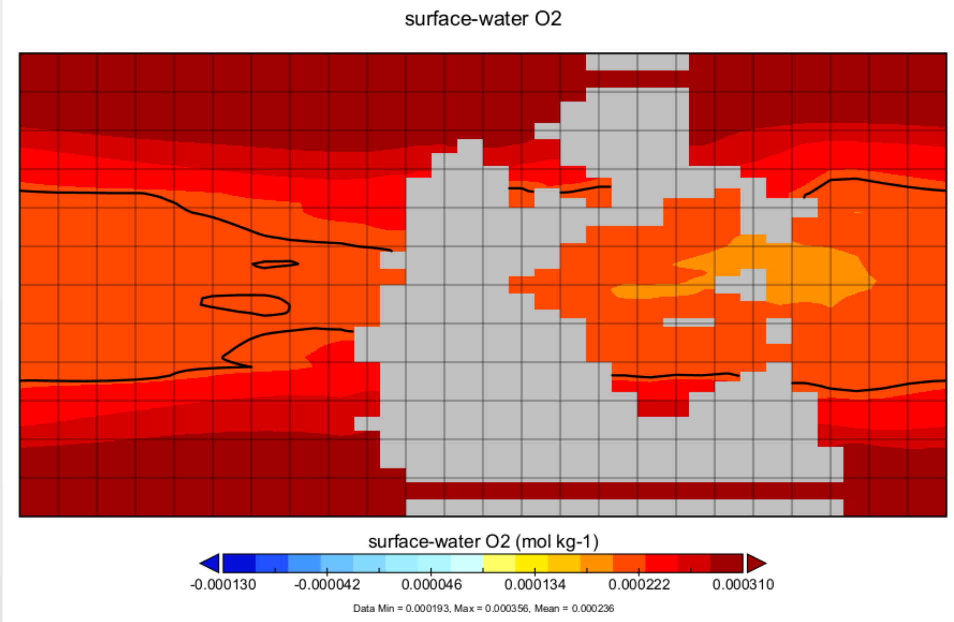


**Figure S25.** Surface water O2 (x10 pCO2).

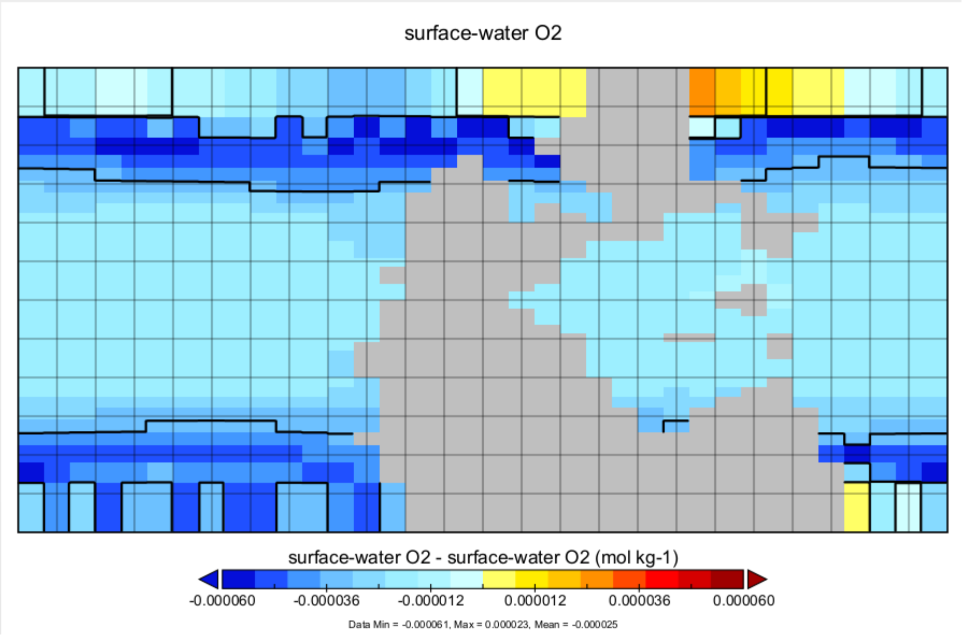


**Figure S26.** Surface water O2 change (from x2 to 10x pCO2).

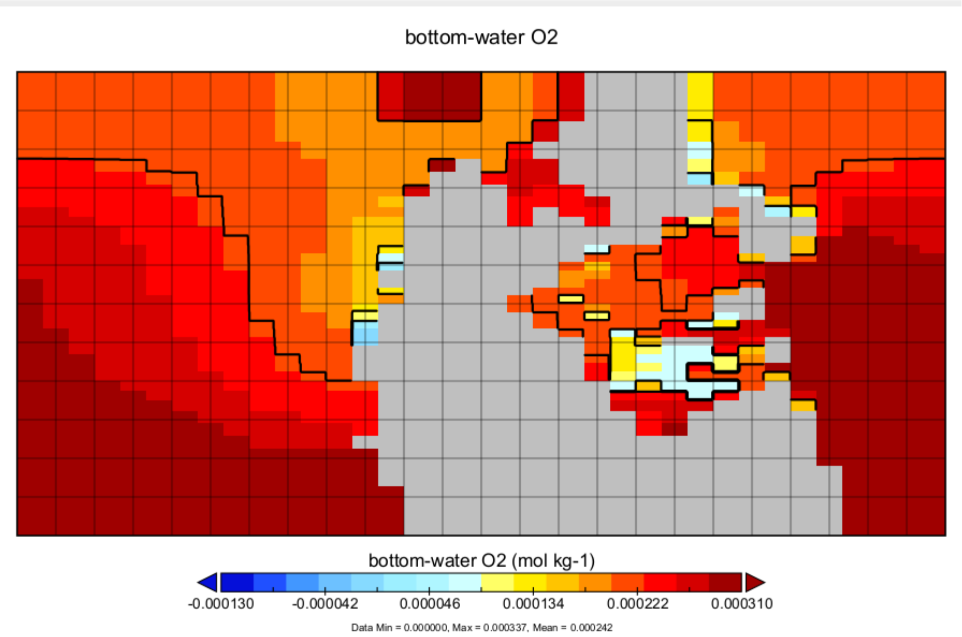


**Figure S27.** Bottom water O2 (x2 pCO2).

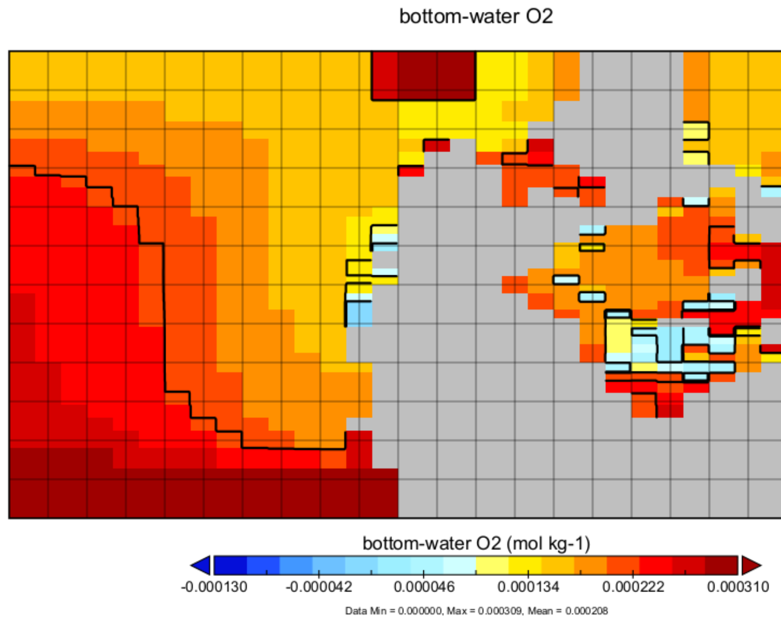


**Figure S28.** Bottom water O2 (x10 pCO2).

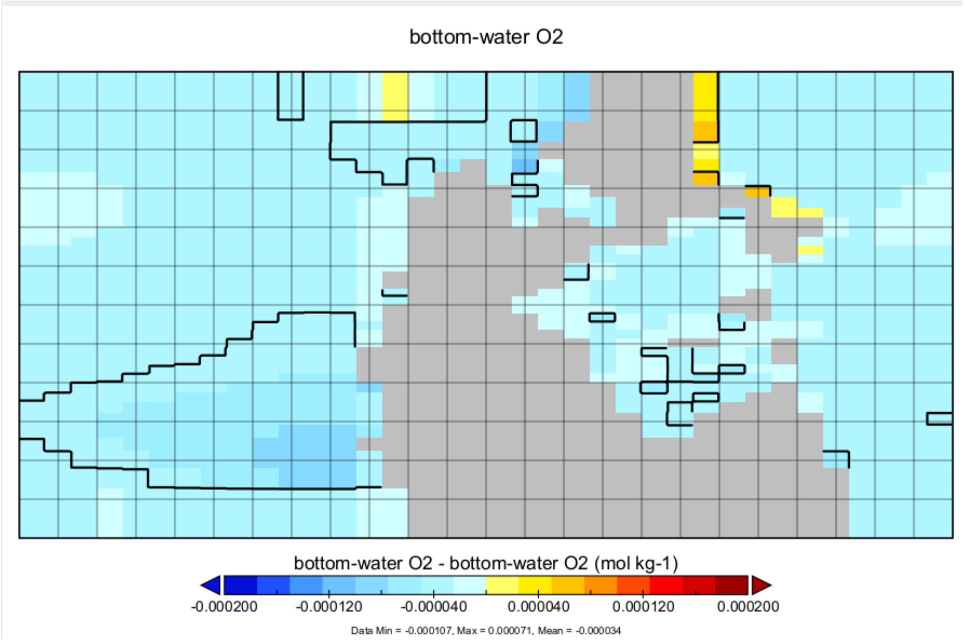


**Figure S29.** Bottom water O2 change (from x2 to 10x pCO2).

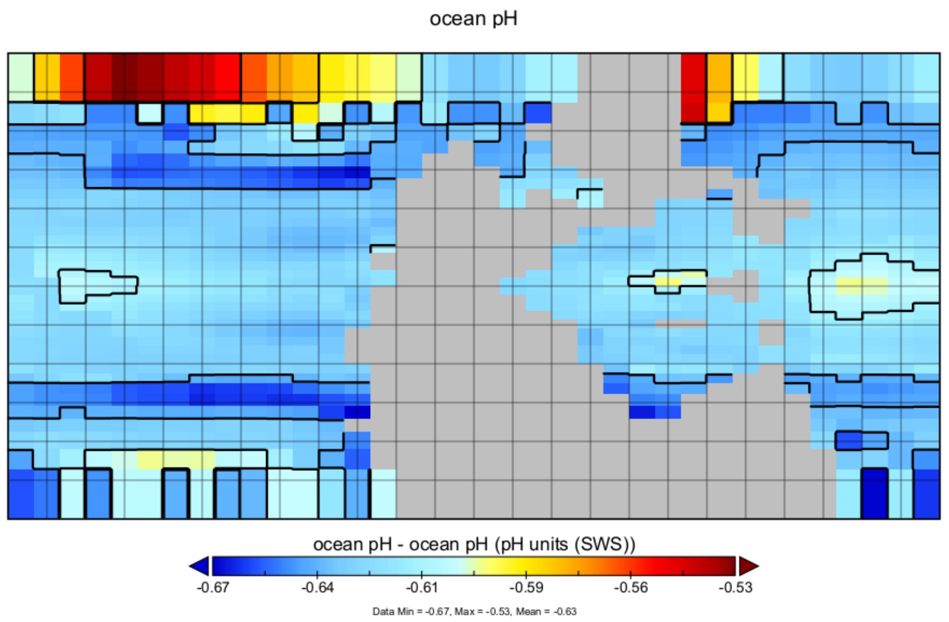


**Figure S30.** pH change (from x2 to 10x pCO2).

### Acknowledgements

Mónica Alejandra Gómez Correa, Andrej Ernst, Hans Hagdorn, Jeffrey Thompson, Ben Thuy, Colin Sumrall, Marie-Béatrice Forel are greatly acknowledged for their identifications of taxa in Türkiye dataset. Erin Dillon is greatly acknowledged for the discussion and giving insights on dermal scales and body size estimation in chondrichthyans. William Foster is acknowledged for providing the dataset that was used by Foster et al. (2022). We acknowledge Luigi Romani (MNHN), Herwig Prinoth (Museum Ladin), Qijian Li (NIGPAS), Haipeng Xu (Nanjing University), Imelda Hausmann (SNSB-BSPG), Carlo Romano (UZH), Torsten Scheyer (UZH), Alex Pohle and Borhan Bagherpour (Ruhr-Universität Bochum) for helping to acquire literature. NERC is acknowledged for the funding (NE/X015025/1).

Karapunar, B., Hagdorn, H., Ernst, A., Thompson, J., Thuy, B., Sumrall, C., Gómez Correa, M. A., Gürsoy, M., Forel, M.-B., Buchwald, S., Frank, A., Kosun, E., Erdem, B., Dunhill, A., & Foster, W. J. (in preparation). Restructuring of marine communities across the Permian–Triassic mass extinction in tropical paleolatitudes — new findings from Türkiye.

Karapunar, B., Wang, X., Frank, A. B., Gürsoy, M., Buchwald, S. Z., Gómez Correa, M. A., Liu, Z., Xu, X., Meng, L., Demir, D., Koşun, E., & Foster, W. J. (2025). Environmental and ecological changes across the Permian–Triassic transition in Türkiye: integrating virtual outcrop models and new fieldwork data. The Depositional Record. *EarthArXiv* *https://doi.org/10.31223/X5KQ9B*.

Karapunar, B., Wolniewicz, A., Romano, C., Ozsvart, P., Rochin-Banaga, H., Kustatscher, E., Buchwald, S. Z., Galasso, F., Davis, D., Lopez-Arbarello, A., Prinoth, H., Bernardi, M., & Foster, W. J. (2025). New insights into the extinction and recovery of marine vertebrates across the Permian-Triassic mass extinction event in the Dolomites, Southern Alps, Italy. *bioRxiv*, 2025.2008.2023.671916. https://doi.org/10.1101/2025.08.23.671916

Kelley, P., Kowalewski, M., & Hansen, T. A. (2003). *Predator-prey interactions in the fossil record* (Vol. 20). Kluwer Academic / Plenum Publishers.

Klapper, G., & Philip, G. M. (1971). Devonian conodont apparatuses and their vicarious skeletal elements. *Lethaia*, *4*(4), 429–452.

Knight, J. B. (1941). Palaeozoic Gastropod Genotypes. *Geological Society of America Special Papers*, *32*, 1–510.

Knoll, A. H., Bambach, R. K., Payne, J. L., Pruss, S., & Fischer, W. W. (2007). Paleophysiology and end-Permian mass extinction. *Earth and Planetary Science Letters*, *256*, 295–313.

Kocsis, Á. T., & Raja, N. B. (2023). chronosphere: Evolving Earth System Variables. R package version 0.6.1. Published on 2023-08-17. doi: 10.32614/CRAN.package.chronosphere.

Kocsis, A. T., Raja, N. B., Williams, S., & Dowding, E. M. (2021). rgplates: R interface for the GPlates Web Service and Desktop Application R package version 0.6.1. Published on 2025-09-11. doi: https://zenodo.org/records/17100022.

Kogan, I. (2011). Remains of *Saurichthys* (Pisces, Actinopterygii) from the Early Triassic Wordie Creek Formation of East Greenland. *Bulletin of the Geological Society of Denmark*, *59*, 93–100.

Kolar-Jurkovšek, T., Chen, Y., Jurkovšek, B., Poljak, M., Aljinović, D., & Richoz, S. (2017). Conodont biostratigraphy of the Early Triassic in eastern Slovenia. *Paleontological Journal*, *51*(7), 687–703.

Korchagin, O. (2011). Foraminifers in the global stratotype (GSSP) of the Permian-Triassic boundary (Bed 27, Meishan, South China). *Stratigraphy and Geological Correlation*, *19*(2), 160–172.

Kozur, H., Mostler, H., & Rahimi-Yazd, A. (1975). Beiträge zur Mikrofauna permotriadischer Schichtfolgen. Teil II: Neue Conodonten aus dem Oberperm und der basalen Trias von Nord- und Zentraliran. *Geologische Paläontologische Mitteilungen Innsbruck*, *5*(3), 1–23.

Kummel, B. (1969). Ammonoids of the Late Scythian (Lower Triassic). *Bull. Mus. Comp. Zool.*, *137*, 311–702.

Kummel, B., & Teichert, C. (1970). Stratigraphy and Paleontology of the Permian-Triassic Boundary Beds, Salt Range and Trans-Indus Ranges, West Pakistan. *Stratigraphic Boundary Problems: Permian and Triassic of West Pakistan, reprint*, *33*, 1–110.

Kutygin, R., Kilyasov, A., & Biakov, A. (2023). The first record of the goniatite genus *Paramexicoceras* in the Changhsingian deposits of the Upper Permian in Northeastern Asia. Doklady Earth Sciences,

Lehman, J. P. (1952). Étude complémentaire des poissons de l'éotrias de Madagascar. *Kungl. Svenska vetenskapsakademiens handlingar, Ser. IV*, *2*(6).

Leighton, L. R. (2003). Predation on brachiopods. In P. Kelley, M. Kowalewski, & T. A. Hansen (Eds.), *Predator-prey interactions in the fossil record* (pp. 215–237). Kluwer Academic / Plenum Publishers.

Leonardi, P. (1935). Il Trias inferiore delle Venezie. *Memorie degli Istituti de Geologia e Mineralogia dell´Università di Padova*, *11*, 1–136.

Leonhard, I., Shirley, B., Murdock, D. J., Repetski, J., & Jarochowska, E. (2021). Growth and feeding ecology of coniform conodonts. *PeerJ*, *9*, e12505.

Lerosey-Aubril, R., & Angiolini, L. (2009). Permian trilobites from Antalya Province, Turkey, and enrollment in Late Palaeozoic trilobites. *Turkish Journal of Earth Sciences*, *18*(3), 427–448.

Li, W.-Z., & Shen, S.-Z. (2008). Lopingian (Late Permian) brachiopods around the Wuchiapingian-Changhsingian boundary at the Meishan Sections C and D, Changxing, South China. *Geobios*, *41*(2), 307–320.

Liao, Z.-t. (1984). New genus and species of Late Permian and earliest Triassic brachiopods from Jiangsu, Zhejiang and Anhui Provinces. *Acta Palaeontologica Sinica*, *23*(3), 276–284.

Liu, G., & Wang, Q. (1994). New material of *Sinohelicoprion* from Changxing, Zhejiang Province. *Vertebrata Palasiatica*, *32*(4), 244–248.

Liu, X., & Wei, F. (1988). A new Saurichthyid from the Upper Permian of Zhejiang, China. *Vertebrata Palasiatica*, *26*(2), 77–89.

Lyu, Z., Orchard, M. J., Golding, M. L., Henderson, C. M., Chen, Z.-Q., Zhang, L., Han, C., Wu, S., Huang, Y., & Zhao, L. (2021). Lower Triassic conodont biostratigraphy of the Guryul Ravine section, Kashmir. *Global and Planetary Change*, *207*, 103671.

Maisey, J. G. (1989). *Hamiltonichthys mapesi*, g. & sp. nov. (Chondrichthyes, Elasmobranchii), from the Upper Pennsylvanian of Kansas. *AMERICAN MUSEUM NOVITATES*, *2931*, 1–42.

Mangum, C. P. (2010). Invertebrate blood oxygen carriers. In R. Terjung (Ed.), *Comprehensive physiology* (pp. 1097–1135). https://doi.org/https://doi.org/10.1002/cphy.cp130215

Marcoux, J., Baud, A., Krystyn, L., & Monod, O. (1986). Guide book part 2, Western Taurides, Antalya – Seydisehir – Isparta – Antalya. Late Permian and Triassic in Western Turkey, Field Workshop 1986,

Margulis, L., & Chapman, M. J. (2009). *Kingdoms and domains: an illustrated guide to the phyla of life on Earth*. Academic Press.

Marrama, G., Lombardo, C., Tintori, A., & Carnevale, G. (2017). Redescription of '*Perleidus*'(Osteichthyes, Actinopterygii) from the early Triassic of northwestern Madagascar. *Rivista Italiana di Paleontologia e Stratigrafia*, *123*(2), 219–242.

Marzola, M., Mateus, O., Milàn, J., & Clemmensen, L. B. (2018). A review of Palaeozoic and Mesozoic tetrapods from Greenland. *Bulletin of the Geological Society of Denmark*, *66*, 21–46.

McClintock, J. B. (1994). Trophic biology of Antarctic shallow-water echinoderms. *Marine Ecology Progress Series*, 191–202.

Mckinney, F. K., Taylor, P. D., & Lidgard, S. (2003). Predation on bryozoans and its reflection in the fossil record. In P. Kelley, M. Kowalewski, & T. A. Hansen (Eds.), *Predator-prey interactions in the fossil record* (pp. 239–261). Kluwer Academic / Plenum Publishers.

Mei, S., Zhang, K., & Wardlaw, B. R. (1998). A refined succession of Changhsingian and Griesbachian neogondolellid conodonts from the Meishan section, candidate of the global stratotype section and point of the Permian–Triassic boundary. *Palaeogeography, Palaeoclimatology, Palaeoecology*, *143*(4), 213–226.

Merla, G. (1930). La fauna del calcare a Bellerophon della regione Dolomitica. *Memorie dell' Istituto Geologico della R. Università di Padova*, *9*, 1–221.

Mette, W., & Roozbahani, P. (2012). Late permian (changsingian) ostracods of the Bellerophon Formation at Seis (Siusi)(dolomites, Italy). *Journal of Micropalaeontolgy*, *31*(1), 73–87.

Middlemiss, C. S. (1910). A revision of the Silurian-Trias sequence in Kashmir. *Records of the Geological Survey of India*, *40*.

Miklukho-Maklay, K. V. (1954). Foraminifery verkhnepermskikh otlozhenii severnogo Kavkaza - Foraminifers from the Late Permian deposits of the northern Caucasus. *Trudy VSEGEI, Gosgeoltekhizdat*, *1*, 1–163.

Monarrez, P. M., Heim, N. A., & Payne, J. L. (2021). Mass extinctions alter extinction and origination dynamics with respect to body size. *Proceedings of the Royal Society B*, *288*(1960), 20211681.

Muromtseva, V. A. (1984). *Permian Marine Deposits and Bivalve Mollusks of the Soviet Arctic - Пермские морские отложения и двустворчатые моллюски Советской Арктики*. Nedra.

Mutter, R. J. (2005). Re-assessment of the genus *Helmolepis* Stensiö 1932 (Actinopterygii: Platysiagidae) and the evolution of platysiagids in the Early-Middle Triassic. *Eclogae Geologicae Helvetiae*, *98*(2), 271–280.

Mutter, R. J., Cartanyà, J., & Basaraba, S. A. (2008). New evidence of *Saurichthys* from the Lower Triassic with an evaluation of early saurichthyid diversity. *Mesozoic fishes*, *4*, 103–127.

Mutter, R. J., & Neuman, A. G. (2008). New eugeneodontid sharks from the Lower Triassic Sulphur Mountain Formation of Western Canada. *Geological Society, London, Special Publications*, *295*, 9–41.

Nakazawa, K. (1981). Permian and Triassic Bivalves of Kashmir. *Palaeontologia Indica, new series*, *46*, 87–122.

Nakazawa, K., & Kapoor, H. (1981). The upper Permian and lower Triassic faunas of Kashmir. *Palaeontologia Indica, new series*, *46*, 1–191.

Nakazawa, K., Kapoor, H. M., Ishii, K.-i., Bando, Y., Okimura, Y., & Tokuoka, T. (1975). The upper Permian and the lower Triassic in Kashmir, India. *Memoirs of the Faculty of Science, Kyoto University. Series of geology and mineralogy*, *42*(1), 1–106.

Neri, C., & Posenato, R. (1985). New biostratigraphical data on uppermost Werfen Formation of western Dolomites (Trento Italy). *Geologisch-Palaontologische Mitteilungen Innsbruck*, *14*(3), 87–107.

Newell, N. D. (1955). Permian Pelecypods of East Greenland. *Meddelelser om Grønland*, *110*(4), 1–36.

Nicoll, R. S., Metcalfe, I., & Cheng-Yuan, W. (2002). New species of the conodont Genus *Hindeodus* and the conodont biostratigraphy of the Permian–Triassic boundary interval. *Journal of Asian Earth Sciences*, *20*(6), 609–631.

Nielsen, E. (1936). Some few preliminary remarks on Triassic fishes from East Greenland. *Meddelelser om Grønland*, *112*(3), 1–55.

Nielsen, E. (1942). Studies on Triassic Fishes from East Greenland: 1. *Glaucolepis* and *Boreosomus*. *Meddelelser om Grønland*, *138*, 1–394.

Nielsen, E. (1949). Studies on Triassic Fishes from East Greenland 2: *Australosomus* and *Birgeria*. *Meddelelser om Grønland*, *146*(1), 1–149.

Nielsen, E. (1952). A preliminary note on *Bobasatrania groenlandica*. *Medd. fra Dansk Geol. Forening. København*, *12*, 197–204.

Noé, S. (1987). Facies and paleogeography of the marine upper Permian and Permian-Triassic boundary in the Southern Alps (Bellerophon Formation, Tesero Horizon). *Facies*, *16*, 89–142.

Nützel, A. (2021). Gastropods as Parasites and Carnivorous Grazers: A Major Guild in Marine Ecosystems. In K. De Baets & J. W. Huntley (Eds.), *The Evolution and Fossil Record of Parasitism: Identification and Macroevolution of Parasites* (pp. 209–229). Springer. https://doi.org/doi.org/10.1007/978-3-030-42484-8_6

Nützel, A., & Karapunar, B. (2023). On Triassic *Murchisonia*-like gastropods—surviving the end-Permian extinction to become extinct in the Late Triassic. *Acta Palaeontologica Polonica*, *68*, 1–21.

Nybelin, O. (1977). Studies on Triassic Fishes from East Greenland III. On *Helmolepis gracilis* Stensiö. *Meddelelser om Grønland*, *200*(2), 1–14.

Ogilvie Gordon, M. M. (1927). Das Gröderner- Fassa- und Enneberggebiet in den Südtiroler Dolomiten Geologische Beschreibung mit besonderer Berücksichtigung der Überschiebungserscheinungen 3. Teil: Paläontologie. *Abhandlungen der Geologischen Bundesanstalt*, *24, Heft 2*, 1–89.

Orchard, M. J., Nassichuk, W. W., & Rui, L. (1994). Conodonts from the lower Griesbachian *Otoceras latilobatum* bed of Selong, Tibet and the position of the Permian—Triassic boundary. *Canadian Society of Petroleum Geologists*, *17*, 823–843.

Pan, H.-z. (1982). Triassic marine gastropods from SW China. *Bulletin of the Nanjing Institute of Geology and Paleontology*, *4*, 153–188.

Pan, H.-Z., & Shen, S.-Z. (2008). Late Permian (Lopingian) Gastropods from the Qubuerga Formation at the Qubu Section in the Mt. Everest (Qomolangma) Region, Southern Tibet (Xizang), China. *Journal of Paleontology*, *82*(5), 1038–1042.

Pasini, M. (1985). Biostratigrafia con i foraminiferi del limite formazione a Bellerophon formazione di Werfen fra Recoaro e La Val Badia (Alpi Meridionali). *Rivista Italiana di Paleontologia e Stratigrafia*, *90*(4), 431–510.

Pattison, J., & Stemmerik, L. (1996). Upper Permian foraminifera from East Greenland. *Bulletin Grønlands Geologiske Undersøgelse*, *171*, 73–90.

Pavlova, E., Manankov, I., Morozova, I., Solovjeva, M., Suetenko, O., & Bogoslovskaya, M. (1991). Permian invertebrates of southern Mongolia. *Transactions*, *40*, 173.

Peel, J. S. (1984). Autecology and systematics of a new Silurian anomphalid gastropod from western North Greenland. *Rapp. Grönlands geol. Unders.*, *121*, 77–87.

Perri, M. C., & Andraghetti, M. (2020). Permian-Triassic Boundary and Early Triassic Conodonts from the Southern Alps, Italy. *Rivista Italiana di Paleontologia e Stratigrafia*, *93*(3), 291–328.

Perri, M. C., & Farabegoli, E. (2003). Conodonts across the Permian–Triassic boundary in the Southern Alps. *Courier Forschungsinstitut Senckenberg*, *245*, 281–313.

Perry, S. F., Lambertz, M., & Schmitz, A. (2019). *Respiratory biology of animals: evolutionary and functional morphology*. Oxford University Press.

Petsios, E., Farrar, L., Tennakoon, S., Jamal, F., Portell, R. W., Kowalewski, M., & Tyler, C. L. (2023). *The Ecology of Biotic Interactions in Echinoids: Modern Insights into Ancient Interactions*. Cambridge University Press.

Plummer, H. J. (1930). Calcareous Foraminifera in the brownwood shale near Bridgeport, Texas; Foraminifera of the Cisco Group of Texas. *The University of Texas Bulletin*, *3019*, 1–81.

Posenato, R. (1985). Un'Associazione oligotipica a *Neoschizodus ovatus* (Goldfuss) della formazione di Werfen (Triassico inf-Dolomiti). *Atti 38 Simposio di Ecologia e Paleoecologia delle Communita bentoniche, Catania*, *3*, 141–153.

Posenato, R. (1992). *Tirolites* (Ammonoidea) from the Dolomites, Bakony and Dalmatia: taxonomy and biostratigraphy. *Eclogae Geologicae Helvetiae*, *85*(3), 893–929.

Posenato, R. (1998). The gen. *Comelicania* Frech, 1901 (Brachiopoda) from the Southern Alps: morphology and classification. *Rivista Italiana di Paleontologia e Stratigrafia*, *104*(1).

Posenato, R. (2001). The athyridoids of the transitional beds between Bellerophon and Werfen formations (uppermost Permian, Southern Alps, Italy). *Rivista Italiana di Paleontologia e Stratigrafia*, *107*(2), 197–226.

Posenato, R. (2009). Survival patterns of macrobenthic marine assemblages during the end-Permian mass extinction in the western Tethys (Dolomites, Italy). *Palaeogeography, Palaeoclimatology, Palaeoecology*, *280*(1-2), 150–167.

Posenato, R. (2010). Marine biotic events in the Lopingian succession and latest Permian extinction in the Southern Alps (Italy). *Geological Journal*, *45*(2-3), 195–215.

Posenato, R. (2011). Latest Changhsingian orthotetid brachiopods in the Dolomites (Southern Alps, Italy): ecological opportunists at the peak of the end-Permian mass extinction. *Journal of Paleontology*, *85*(1), 58–68.

Posenato, R. (2016). Systematics of lingulide brachiopods from the end-Permian mass extinction interval. *Rivista Italiana di Paleontologia e Stratigrafia*, *122*(2), 85–108.

Posenato, R. (2019). The end-Permian mass extinction (EPME) and the Early Triassic biotic recovery in the western Dolomites (Italy): state of the art. *Bollettino della Società Paleontologica Italiana*, *58*(1), 11–34.

Posenato, R., Holmer, L. E., & Prinoth, H. (2014). Adaptive strategies and environmental significance of lingulid brachiopods across the late Permian extinction. *Palaeogeography, Palaeoclimatology, Palaeoecology*, *399*, 373–384.

Posenato, R., Pelikán, P., & Hips, K. (2005). Bivalves and brachiopods near the Permian-Triassic boundary from the Bükk Mountains (Bálvány-North section, northern Hungary). *Rivista Italiana di Paleontologia e Stratigrafia*, *111*(2).

Posenato, R., & Prinoth, H. (1999). Discovery of *Paratirolites* from the Bellerophon Formation (Upper Permian, Dolomites, Italy). *Rivista Italiana di Paleontologia e Stratigrafia*, *105*(1).

Posenato, R., & Prinoth, H. (2004). Orizzonti a nautiloidi ea brachiopodi della Formazione a Bellerophon (Permiano Superiore) in Val Gardena (Dolomiti). *Geo. Alp*, *1*, 71–85.

Posenato, R., Sciunnach, D., & Garzanti, E. (1996). First report of *Claraia* (bivalvia) in the servino formation (Lower Triassic) of the western Orobic Alps, Italy. *Rivista Italiana di Paleontologia e Stratigrafia*, *102*(2), 201–210.

Prinoth, H., & Posenato, R. (2007). Late Permian Nautiloids from the Bellerophon Formation of the Dolomites (Italy). *Palaeontographica Abteilung A Paläozoologie, Stratigraphie*, *282*(1-6), 135–165.

Prinoth, H., & Posenato, R. (2023). Bivalves from the Changhsingian (upper Permian) Bellerophon Formation of the Dolomites (Italy): ancestors of Lower Triassic post-extinction benthic communities. *Papers in Palaeontology*, *9*(2), e1486.

Raschi, W. G., & Musick, J. A. (1984). Hydrodynamic aspects of shark scales. *Special Report in Applied Marine Science and Ocean Engineering*(272), 182.

Rauzer-Chernousova, D. M. V., A. Y.; Glebovskaya, E. M.; Grozdilova, L. P.; Lipina, O. A.; Suleimanov, I. S.; Tchernysheva, N. E. (1948). *Stratigraphy and Foraminifera of the Lower Carboniferous of the Russian Platform and Fore-Urals -Стратиграфия и фораминиферы нижнего карбона Русской платформы и Приуралья. Akademya Nauk SSSR Trudy Instituta Geologicheskikh Nauk. 62 (ser. geol. no. 19): 1-260 (in Russian).*

Reed, F. R. C. (1931). New fossils from the Productus Limestones of the Salt Range, with notes on other species. *Memoirs of the Geological Survey of India, Palaeontologia Indica*, *17*, 1–56.

Reichel, M. (1946). Sur quelques foraminifères nouveaux du Permien méditerranéen. *Ecologae Geologicae Helvetiaef*, *38*(2), 524–560.

Reid, C. M., & Tamberg, Y. (2021). Trophic partitioning and feeding capacity in Permian bryozoan faunas of Gondwana. *Palaeontology*, *64*(4), 555–572.

Reitlinger, E. A. (1950). Foraminifera of the Middle Carboniferous deposits of the central part of the Russian Platform (excluding the family Fusulinidae) - Фораминиферы среднекаменноугольных отложений центральной части Русской платформы (исключая сем. Fusulinidae). *Proceedings of the Institute of Geological Sciences, Academy of Sciences of the USSR, Geological Series*, *126*, 1–128.

Richardson, J. (1997). Ecology of articulated brachiopods. *Introduction*, *1*, 441–462.

Richoz, S. (2006). Stratigraphie et variations isotopiques du carbone dans le Permien supérieur et le Trias inférieur de quelques localités de la Néotéthys (Turquie, Oman et Iran). *Mémoires de Géologie (Lausanne)*, *46*, 1–251.

Ridgwell, A., Hargreaves, J., Edwards, N. R., Annan, J., Lenton, T. M., Marsh, R., Yool, A., & Watson, A. (2007). Marine geochemical data assimilation in an efficient Earth System Model of global biogeochemical cycling. *Biogeosciences*, *4*(1), 87–104.

Romano, C., Koot, M. B., Kogan, I., Brayard, A., Minikh, A. V., Brinkmann, W., Bucher, H., & Kriwet, J. (2016). Permian–Triassic Osteichthyes (bony fishes): diversity dynamics and body size evolution. *Biological Reviews*, *91*(1), 106–147.

Ronchi, A., Santi, G., Marchetti, L., Bernardi, M., & Gianolla, P. (2018). First report on swimming trace fossils of fish from the Upper Permian and Lower Triassic of the Dolomites (Italy). Annales Societatis Geologorum Poloniae,

Ros-Franch, S., Márquez-Aliaga, A., & Damborenea, S. E. (2014). Comprehensive database on Induan (Lower Triassic) to Sinemurian (Lower Jurassic) marine bivalve genera and their paleobiogeographic record. *Paleontological Contributions*(8), 1–219. https://doi.org/10.17161/PC.1808.13433

Rui, L., He, J.-w., Chen, C.-z., & Wang, Y.-g. (1988). Discovery of fossil animals from the basal clay of Permian–Triassic boundary in the Meishan area of Changxing, Zhejiang and its significance. *Journal of Stratigraphy*, *12*(1), 48–52.

Sakagami, S. (1981). Upper Permian Bryozoa from Guryul Ravine and the Spur Three kilometres North of Barus. *The Upper Permian and Lower Triassic faunas of Kashmir, Palaeontologia Indica New Series*, *46*, 47–62.

Sakagami, S., Sciunnach, D., & Garzanti, E. (2006). Late Paleozoic and Triassic bryozoans from the Tethys Himalaya (N India, Nepal and S Tibet). *Facies*, *52*(2), 279–298.

Säve-Söderbergh, G. (1935). On the dermal bones of the head in labyrinthodont stegocephalians and primitive reptilia: with special reference to Eotriassic stegocephalians from East Greenland. *Medd Gronl*, *98*, 1.

Schaal, E. K. (2014). *Permian-Triassic Global Change: the Strontium cycle and body size evolution in marine clades*. Stanford University.

Scheyer, T. M., Romano, C., Jenks, J., & Bucher, H. (2014). Early Triassic marine biotic recovery: the predators' perspective. *Plos One*, *9*(3), e88987.

Schoch, R. R., & Milner, A. R. (2000). *Handbuch Der Paläoherpetologie: Encyclopedia of Paleoherpetology Teil 3B Stereospondyli*. Dr. Friedrich Pfeil.

Scotese, C., & Wright, N. (2018). PALEOMAP paleodigital elevation models (PaleoDEMS) for the phanerozoic PALEOMAP project. *EarthByte* *https://www.earthbyte.org/paleodem-resource-scotese-and-wright-2018*.

Scotese, C. R. (2016). PALEOMAP PaleoAtlas for GPlates and the PaleoData Plotter Program. https://www.earthbyte.org/paleomap-paleoatlas-for-gplates/.

Shaw, J. O., Coco, E., Wootton, K., Daems, D., Gillreath-Brown, A., Swain, A., & Dunne, J. A. (2021). Disentangling ecological and taphonomic signals in ancient food webs. *Paleobiology*, *47*(3), 385–401.

Shaw, J. O., Dunhill, A. M., Beckerman, A. P., Dunne, J. A., & Hull, P. M. (2024). A framework for reconstructing ancient food webs using functional trait data. *bioRxiv*, 2024.2001.2030.578036. https://doi.org/10.1101/2024.01.30.578036

Shen, S.-Z., Archbold, N. W., Shi, G.-R., & Zhong-Qiang, C. (2000). Permian brachiopods from the Selong Xishan section, Xizang (Tibet), China Part 1: stratigraphy, Strophomenida, Productida and Rhynchonellida. *Geobios*, *33*(6), 725–752.

Shen, S.-Z., Cao, C., Shi, G., Wang, X., & Mei, S. (2003). Lopingian (Late Permian) stratigraphy, sedimentation and palaeobiogeography in southern Tibet. *Newsletters on Stratigraphy*, *39*(2/3), 157–179.

Shen, S.-Z., Shi, G., & Archbold, N. (2003). Lopingian (late Permian) brachiopods from the Qubuerga Formation at the Qubu section in the Mt. Qomolangma region, southern Tibet (Xizang), China.

Shen, S., Archbold, N. W., Shi, G. R., & Chen, Z.-Q. (2001). Permian Brachiopods from the Selong Xishan Section, Xizang ( Tibet), China. Part 2: Palaeobiogeographical and Palaeontological Implications, Spiriferida, Athyridida and Terebratulida. *Geobios*, *34, 2*, 157–182.

Shen, S., Jin, Y., Zhang, Y., & Weldon, L. (2017). Permian brachiopod genera on type species of China. In J. Y. Rong, Y. G. Jin, S. Z. Shen, & R. B. Zhan (Eds.), *Phanerozoic Brachiopod Genera of China* (pp. 651–881). Science Press.

Shen, S., & Yugan, J. (1999). Brachiopods from the Permian–Triassic boundary beds at the Selong Xishan section, Xizang (Tibet), China. *Journal of Asian Earth Sciences*, *17*(4), 547–559.

Shen, S. Z., Cao, C.-Q., Henderson, C. M., Wang, X.-D., Shi, G. R., Wang, Y., & Wang, W. (2006). End-Permian mass extinction pattern in the northern peri-Gondwanan region. *Palaeoworld*, *15*(1), 3–30.

Sheng, H. B. (1988). New material of Early Permian ammonoids from Xizang and Qinghai. *Professional Papers of Stratigraphy and Palaeontology*, *20*, 76–84.

Sheng, J.-Z., Chen, C.-z., Wang, Y.-g., Rui, L., Liao, Z.-t., Bando, Y., Ishii, K.-i., Nakazawa, K., & Nakamura, K. (1984). Permian-Triassic boundary in middle and eastern Tethys. *北海道大学理学部紀要*, *21*(1), 133–181.

Sheng, J.-z., Chen, C.-z., Wang, Y.-g., Rui, L., Liao, Z.-t., & Jiang, N. (1983). A research in the Permian-Triassic boundary stratotype from Changxing area, Zhejiang. *Journal of Stratigraphy*, *7*(4), 245–257.

Shi, C.-g. (1987). The Changhsingian ostracodes from Meishan, Changxing, Zhejiang. *Stratigraphy and Palaeontology of Systemic Boundaries in China. Permian Triassic Boundary*, *1*, 23–80.

Shi, G., Zhang, Y.-c., Shen, S.-z., & He, W.-h. (2016). Nearshore–offshore–basin species diversity and body size variation patterns in Late Permian (Changhsingian) brachiopods. *Palaeogeography, Palaeoclimatology, Palaeoecology*, *448*, 96–107.

Shigeta, Y., Zakharov, Y. D., Maeda, H., & Popov, A. M. (2009). The lower Triassic system in the Abrek Bay area, South Primorye, Russia. *National Museum of Nature and Science Monographs*, *38*, 1–218.

Shimizu, D. (1981). Upper Permian brachiopod fossils from Guryul Ravine and the Spur three kilometers north of Barus. *Palaeontologica Indica, new series*, *46*, 65–87.

Song, H.-J., Tong, J.-N., Zhang, K.-X., Wang, Q.-X., & Chen, Z. (2007). Foraminiferal survivors from the Permian-Triassic mass extinction in the Meishan section, South China. *Palaeoworld*, *16*(1-3), 105–119.

Song, H., Tong, J., & Chen, Z. (2009). Two episodes of foraminiferal extinction near the Permian–Triassic boundary at the Meishan section, South China. *Australian Journal of Earth Sciences*, *56*(6), 765–773.

Song, H., Tong, J., & He, W. (2006). Latest Permian small foraminiferal fauna at the Meishan section, Zhejiang Province. *Wei ti gu Sheng wu xue bao= Acta Micropalaeontologica Sinica*, *23*(2), 87–104.

Song, H., Wignall, P. B., Tong, J., & Yin, H. (2013). Two pulses of extinction during the Permian–Triassic crisis. *Nature Geoscience*, *6*(1), 52–56.

Song, H., Wu, Y., Dai, X., Dal Corso, J., Wang, F., Feng, Y., Chu, D., Tian, L., Song, H., & Foster, W. J. (2024). Respiratory protein-driven selectivity during the Permian-Triassic mass extinction. *The Innovation*, *5*(3).

Sørensen, A. M., Håkansson, E., & Stemmerik, L. (2008). Upper Permian bryozoans of central East Greenland. *Bulletin of the Geological Society of Denmark*, *56*, 39–51.

Spath, L. F. (1930). The Eotriassic Invertebrate fauna of East Greenland. *Meddeleser om Gronland*, *83*(1), 1–90.

Spath, L. F. (1935). Additions to the Eotriassic Invertebrate fauna of East Greenland. *Meddeleser om Gronland*, *98*(1), 1–115.

Stache, G. (1877). Beiträge zur Fauna der Bellerophonkalke Südtirols, Nr. 1. Cephalopoden und Gastropoden. *Jahrbuch der kaiserlich-königlichen geologischen Reichsanstalt*, *27*(Heft 3), 271–318.

Stache, G. (1878). Beiträge zur Fauna der Bellerophonkalke Südtirols. Nr. 2. Pelecypoden und Brachiopoden. *Jahrbuch der kaiserlich-königlichen geologischen Reichsanstalt*, *28*(Heft 1), 93–168.

Stemmerik, L., Bendix-Almgreen, S. E., & Piasecki, S. (2001). The Permian–Triassic boundary in central East Greenland: past and present views. *Bulletin of the Geological Society of Denmark*, *48*(2), 159–167.

Stensiö, E. A. (1932). Triassic Fishes from East Greenland: Collected by the Danish Expeditions in 1929-31. *Meddelelser om Grønland*, *83*(3), 1–305.

Strank, A. (1983). New stratigraphically significant Foraminifera from the Dinantian of Great Britain. *Palaentology*, *26*(2), 435–442.

Studies, N. G. I. f. S. (2024). *Panoply netCDF, HDF and GRIB data viewer (Version 5.5)*. In https://www.giss.nasa.gov/tools/panoply/

Stumpf, S., López-Romero, F. A., Kindlimann, R., Lacombat, F., Pohl, B., & Kriwet, J. (2021). A unique hybodontiform skeleton provides novel insights into Mesozoic chondrichthyan life. *Papers in Palaeontology*, *7*(3), 1479–1505.

Surlyk, F., Bjerager, M., Piasecki, S., & Stemmerik, L. (2017). Stratigraphy of the marine Lower Triassic succession at Kap Stosch, Hold with Hope, North-East Greenland. *Bulletin of the Geological Society of Denmark*, *65*, 87–123.

Sweet, W. C. (1970). Permian and Triassic conodonts from a section at Guryul Ravine, Vihi district, Kashmir. *The University of Kansas, Paleontological Contributions - Paper 49*, 1–10.

Sweet, W. C., & Donoghue, P. C. (2001). Conodonts: past, present, future. *Journal of Paleontology*, *75*(6), 1174–1184.

Taraz, H., Golshani, F., Nakazawa, K., Shimizu, D., Bando, Y., Ishii, K.-i., Murata, M., Okimura, Y., Sakagami, S., Nakamura, K., & Tokuoka, T. (1981). The Permian and the Lower Triassic Systems in Abadeh region, central Iran. *Mem. Fac. Sci., Kyoto Univ., Ser. Geol. Mineral.*, *47*, 61–133.

Taylor, P. D., & Vinn, O. (2006). Convergent morphology in small spiral worm tubes ('*Spirorbis*') and its palaeoenvironmental implications. *Journal of the Geological Society*, *163*(2), 225–228.

Teichert, C., & Kummel, B. (1972). Permian-Triassic boundary in the Kap Stosch area, East Greenland. *Bulletin of Canadian Petroleum Geology*, *20*(4), 659–675.

Terrill, D. F., Jarochowska, E., Henderson, C. M., Shirley, B., & Bremer, O. (2022). Sr/Ca and Ba/Ca ratios support trophic partitioning within a Silurian conodont community from Gotland, Sweden. *Paleobiology*, *48*(4), 601–621.

Thompson, J. R., Posenato, R., Bottjer, D. J., & Petsios, E. (2019). Echinoids from the Tesero Member (Werfen Formation) of the Dolomites (Italy): implications for extinction and survival of echinoids in the aftermath of the end-Permian mass extinction. *PeerJ*, *7*, e7361.

Thompson, M. L. (1942). New genera of Pennsylvanian fusulinids. *American Journal of Science*, *240*(6), 403–420.

Tommasi, A. (1882). *Il Trias inferiore delle nostre Alpi coi suoi giacimenti metalliferi: il Pizzo dei tre Signori*. Casa editrice dottor Francesco Vallardi.

Tommasi, A. (1895). La fauna del Trias inferiore nel versante meridionale delle Alpi. *Palaeontographia Italica: Memorie di paleontologia*, *1*, 43–76.

Tommasi, A. (1899). Alcuni fossili nuovi nel trias inferiore delle nostre Alpi. *Rendiconti del Reale. Istituto Lombardo di Scienze e Lettere*, *32*, 771–774.

Tozer, E. T. (1967). A standard for Triassic time. *Geological Survey of Canada Bulletin*, *156*, 1–103.

Tozer, E. T. (1994). Canadian Triassic ammonoid faunas. *Geological Survey of Canada Bulletin*, *467*, 1–663.

Trümpy, R. (1969). Lower Triassic ammonites from Jameson Land (East Greenland). *Meddelelser om Grønland*, *168*(2), 77–116.

Tscherdynzew, W. (1914). Zur Foraminiferen-Fauna der permischen Ablagerungen des östlichen Teils des europäischen Russlands. *Trudy Obshchestva Estestvoispytatelei pri Imperatorskom Kazanskom Universitete*, *46*(5), 1–88.

Twitchett, R. (1997). *Palaeoenvironments of the Lower Triassic of the Dolomites, Northern Italy* University of Leeds].

Twitchett, R. J. (1996). The resting trace of an acorn-worm (Class: Enteropneusta) from the Lower Triassic. *Journal of Paleontology*, *70*(1), 128–131.

Twitchett, R. J. (1999). Palaeoenvironments and faunal recovery after the end-Permian mass extinction. *Palaeogeography, Palaeoclimatology, Palaeoecology*, *154*, 27–37.

Twitchett, R. J., & Barras, C. G. (2004). Trace fossils in the aftermath of mass extinction events. In D. McIlroy (Ed.), *The Application of Ichnology to Palaeoenvironmental and Stratigraphic Analysis* (Vol. Publ. 228, pp. 397–418). Geol. Soc. Lond., Spec.

Twitchett, R. J., Looy, C. V., Morante, R., Visscher, H., & Wignall, P. B. (2001). Rapid and synchronous collapse of marine and terrestrial ecosystems during the end-Permian biotic crisis. *Geology*, *29, no. 4*, 351–354.

Twitchett, R. J., & Wignall, P. B. (1996). Trace fossils and the aftermath of the Permo-Triassic mass extinction: evidence from northern Italy. *Palaeogeography, Palaeoclimatology, Palaeoecology*, *124*, 137–151.

Uchman, A., Lebanidze, Z., Beridze, T., Kobakhidze, N., Lobzhanidze, K., Makadze, D., Khutsishvili, S., Chagelishvili, R., Koiava, K., & Khundadze, N. (2022). Revision of the trace fossil *Megagrapton* Książkiewicz, 1968 with focus on *Megagrapton aequale* Seilacher, 1977 from the lower Eocene of the Lesser Caucasus in Georgia.

Vachard, D., & Krainer, K. (2022). Calcareous algae and foraminifers across the Permian-Triassic boundary interval (uppermost Bellerophon Formation and basal Werfen Formation) in the Dolomites (South Tyrol-Trentino, Italy); Calcareous algae and foraminifers across the Permian-Triassic boundary interval (uppermost Bellerophon Formation and basal Werfen Formation) in the Dolomites (South Tyrol-Trentino, Italy). *Palaeontographica Abteilung A: Palaeozoology – Stratigraphy*, *324*(1-6), 1–173.

Vannier, J., Abe, K., & Ikuta, K. (1998). Feeding in myodocopid ostracods: functional morphology and laboratory observations from videos. *marine biology*, *132*(3), 391–408.

Vissarionova, A. Y. K., G.D.; Lipina, O.A.; Morozova, V.G.; Rauser-Chernousova, D.M.; Suleimanov, I.S.; Shamov, D.F.; Shcherbovich, S.F. (1949). *Foraminifera of the Upper Carboniferous and Artinskian deposits of the Bashkir Cis-Urals. Trudy Instituta Geologicheskikh Nauk. Vol. 105, Geological series (No. 35).* . Publishing House of the Academy of Sciences of the USSR.

Waagen, W. (1879-1887). *Salt-Range Fossils: Productus-Limestone Fossils*. Geological Survey Office.

Wang, L., Wignall, P. B., Sun, Y., Yan, C., Zhang, Z., & Lai, X. (2017). New Permian-Triassic conodont data from Selong (Tibet) and the youngest occurrence of *Vjalovognathus*. *Journal of Asian Earth Sciences*, *146*, 152–167.

Wang, N., Jin, F., Wang, W., & Zhu, X. (2007). Actinopterygian fishes from the Permian-Triassic boundary beds in Zhejiang and Jiangxi Provinces, South China And fish mass extinction, recovery and radiation. *Vertebrata Palasiatica*, *45*(4), 307–329.

Wang, N., Zhu, X., Jin, F., & Wang, W. (2007). Chondrichthyan microremains under Permian-Triassic boundary both in Zhejiang and Jiangxi provinces, China-Fifth report on the fish sequence study near the Permian-Triassic boundary in South China. *Vertebrata Palasiatica*, *45*(1), 13–36.

Wang, N. C., & Liu, H. T. (1981). Coelacanth fishes from the marine Permian of Zhejiang, South China. *Vertebrata Palasiatica*, *19*(4), 305–&.

Wang, Y.-g. (1984). Earliest triassic ammonoid fauna from Jiangsu and Zhejiang and their bearing on the definition of Permo-Triassic boundary. *Acta Palaeontologica Sinica*, *23*(3), 257–269.

Wang, Y., Shen, S., Cao, C., Henderson, C., Zhu, G.-p., Liu, J.-m., & Xu, Y.-l. (2004). Re-Study on the Wuchiapingian-Changsingian Boundary Section at Meishan, Changxing, Zhejiang Province. *Journal of Stratigraphy*, *28*(1), 27–34.

Wang, Y., Shen, S., Cao, C., Wang, W., Henderson, C., & Jin, Y. (2006). The Wuchiapingian–Changhsingian boundary (Upper Permian) at Meishan of Changxing County, South China. *Journal of Asian Earth Sciences*, *26*(6), 575–583.

Ward, B. A., Wilson, J. D., Death, R. M., Monteiro, F. M., Yool, A., & Ridgwell, A. (2018). EcoGEnIE 1.0: plankton ecology in the cGEnIE Earth system model. *Geoscientific Model Development*, *11*(10), 4241–4267.

Ward, P., Barord, G. J., Schauer, A., & Veloso, J. (2023). Comparative trophic levels of phragmocone-bearing cephalopods (nautiloids, ammonoids, and sepiids). *Integr Comp Biol*, *63*(6), 1285–1297.

Waterhouse, J. B. (1983). A Late Permian lyttoniid fauna from northwest Thailand.

Waterhouse, J. B. (2002). Classification within Productidina and Strophalosiidina (Brachiopoda). *Earthwise*, *5*, 1–60.

Waterhouse, J. B. (2004). Permian and Triassic stratigraphy and fossils of the Himalaya in northern Nepal. *Earthwise*, *6*, 1–288.

Wei, F. (1977). On the occurrence of platysomid in the Changsing (Changxing) limestone of Zhejiang. *Acta Palaeontologica Sinica*, *16*(2), 293–298.

Wei, H., Zhang, X., & Qiu, Z. (2020). Millennial-scale ocean redox and δ13C changes across the Permian–Triassic transition at Meishan and implications for the biocrisis. *International Journal of Earth Sciences*, *109*(5), 1753–1766.

Wignall, P. B., & Hallam, A. (1992). Anoxia as a cause of the Permian/Triassic mass extinction: facies evidence from northern Italy and the western United States. *Palaeogeography, Palaeoclimatology, Palaeoecology*, *93*(1-2), 21–46.

Wignall, P. B., & Newton, R. (2003). Contrasting Deep-water Records from the Upper Permian and Lower Triassic of South Tibet and British Columbia: Evidence for a Diachronous Mass Extinction. *Palaios*, *18*, 153–167.

Wignall, P. B., & Twitchett, R. J. (2002). Permian-Triassic sedimentology of Jameson Land, East Greenland: incised submarine channels in an anoxic basin. *Journal of the Geological Society of London*, *159*, 691–703.

Wilson, J., Monteiro, F., Schmidt, D., Ward, B., & Ridgwell, A. (2018). Linking marine plankton ecosystems and climate: a new modeling approach to the warm early Eocene climate. *Paleoceanography and Paleoclimatology*, *33*(12), 1439–1452.

Wittenburg, P. v. (1908a). Beiträge zur Kenntnis der Werfener Schichten Südtirols. *Geologische und Palaeontologische Abhandlungen*, *8, Heft 5*, 251–289.

Wittenburg, P. v. (1908). Einige neue Fossilien aus den Werfener Schichten Südtirols. *Neues Jahrbuch für Mineralogie etc.*, *1*, 16–21.

Wittenburg, P. v. (1908b). Neue Beiträge zur Geologie und Paläontologie der Werfener Schichten Südtirols, mit Berücksichtigung der Schichten von Wladiwostok. *Centralblatt für Mineralogie, Geologie und Paläontologie, Jahrgang 1908*, *2*, 67–89.

Wood, S. A., Russell, R., Hanson, D., Williams, R. J., & Dunne, J. A. (2015). Effects of spatial scale of sampling on food web structure. *Ecology and Evolution*, *5*(17), 3769–3782.

Wu, H., Zhang, Y., Stubbs, T. L., Chen, A., Zhai, P., & Sun, Y. (2023). Wuchiapingian (Lopingian, late Permian) brachiopod fauna from Guangdong Province, southeastern China: systematics and contribution to the Lopingian recovery. *Journal of Paleontology*, *97*(1), 112–139.

Wu, W.-S. (1975). The coral fossils from Qomolangma Feng Region. *A report of Scientific expedition in the Mount Jolmo Lungma region (1966^ 1968). Palaeontology, Fasc. I, Sci. Press, Beijing*, 83–113.

Wu, Y., Chu, D., Tong, J., Song, H., Dal Corso, J., Wignall, P. B., Song, H., Du, Y., & Cui, Y. (2021). Six-fold increase of atmospheric p $CO_2$ during the Permian–Triassic mass extinction. *Nature Communications*, *12*(1), 2137.

Xu, G., & Grant, R. E. (1994). Brachiopods near the Permian-Triassic boundary in south China.

Xu, H.-P., Cao, C.-Q., Yuan, D.-X., Zhang, Y.-C., & Shen, S.-Z. (2018). Lopingian (late Permian) brachiopod faunas from the Qubuerga Formation at Tulong and Kujianla in the Mt. Everest area of southern Tibet, China. *Rivista Italiana di Paleontologia e Stratigrafia*, *124*(1).

Yadrenkin, A., Biakov, A., Kutygin, R., & Kopylova, A. (2020). New findings and stratigraphic distribution of foraminifera from Permian–Triassic boundary deposits in the Southern Verkhoyansk Region. *Russian Journal of Pacific Geology*, *14*(5), 447–459.

Yang, J. Z., & Xia, F. S. (1975). Bryozoan fossils from the Mount Qomolangma Region. *A report of scientific expedition in the Mount Jolmo Lungma Region (1966-1968)*, *1*, 39–70.

Yang, L., Dai, X., Liu, X., Feng, Y., Jiang, S., Wang, F., Song, H., Tian, L., & Song, H. (2024). Foraminiferal extinction and size reduction during the Permian-Triassic transition in southern Tibet. *Journal of Earth Science*, *35*(6), 1799–1809.

Yang, Z., Yang, F., & Wu, S. (1996). The ammonoid *Hypophiceras* fauna near the Permian-Triassic boundary at Meishan section and in South China: stratigraphic significance. In H. Yin (Ed.), *The Palaeozoic-Mesozoic boundary candidates of global stratotype section and point of the Permian-Triassic boundary* (pp. 49–56). China University of Geosciences Press.

Yang, Z., Yin, H., Wu, S., Yang, F., Ding, M., & Xu, G. (1987). Permian-Triassic boundary stratigraphy and fauna of South China. *PRC Ministry of Geology and Mineral Resources, Geological Memoirs, Series*, *2*(6), 1–378.

Yin, H. (1996). *The Palaeozoic-Mesozoic boundary, candidates of Global Stratotype Section and Point of the Permian-Triassic boundary*. China University of Geoscience Press.

Yin, H., Wu, S., Ding, M., Zhang, K., Tong, J., Yang, J., & Lai, X. (1996). The Meishan section, candidate of the Global Stratotype Section and Point of the Permian-Triassic boundary. . In H.-f. Yin (Ed.), *The Palaeozoic-Mesozoic Boundary Candidates of Global Stratotype Section and Point of the Permian-Triassic Boundary* (pp. 31–48). China University of Geosciences Press.

Yochelson, E. L., & Hongfu, Y. (1985). Redescription of Bellerophon asiaticus Wirth (early Triassic: Gastropoda) from China, and a survey of Triassic Bellerophontacea. *Journal of Paleontology*, *59*(5), 1305–1319.

Yuan, D.-X., Zhang, Y.-C., & Shen, S.-Z. (2018). Conodont succession and reassessment of major events around the Permian-Triassic boundary at the Selong Xishan section, southern Tibet, China. *Global and Planetary Change*, *161*, 194–210.

Yuan, Z., Xu, G.-H., Dai, X., Wang, F., Liu, X., Jia, E., Miao, L., & Song, H. (2022). A new perleidid neopterygian fish from the Early Triassic (Dienerian, Induan) of South China, with a reassessment of the relationships of Perleidiformes. *PeerJ*, *10*, e13448.

Zakharov, Y. D. (2002). Ammonoid succession of Setorym River (Verkhoyansk area) and problem of Permian-Triassic boundary in Boreal Realm. *Journal of Earth Science*, *13*(2), 107–123.

Zakharov, Y. D., Biakov, A., & Horacek, M. (2014). Global correlation of basal Triassic layers in the light of the first carbon isotope data on the Permian-Triassic boundary in Northeast Asia. *Russian Journal of Pacific Geology*, *8*(1), 1–17.

Zatoń, M., Niedźwiedzki, G., Blom, H., & Kear, B. P. (2016). Boreal earliest Triassic biotas elucidate globally depauperate hard substrate communities after the end-Permian mass extinction. *Scientific Reports*, *6*(1), 36345.

Zatoń, M., Niedźwiedzki, G., Rakociński, M., Blom, H., & Kear, B. P. (2018). Earliest Triassic metazoan bioconstructions from East Greenland reveal a pioneering benthic community in the immediate aftermath of the end-Permian mass extinction. *Global and Planetary Change*, *167*, 87–98.

Zhang, C., Bucher, H., & Shen, S.-Z. (2017). Griesbachian and Dienerian (early Triassic) ammonoids from Qubu in the Mt. Everest area, southern Tibet. *Palaeoworld*, *26*(4), 650–662.

Zhang, K., Lai, X., Tong, J., & Jiang, H. (2009). Progresses on study of conodont sequence for the GSSP section at Meishan, Changxing, Zhejiang Province, South China. *Acta Palaeontologica Sinica*, *48*(3), 474–486.

Zhang, K., Tong, J., Shi, G. R., Lai, X., Yu, J., He, W., Peng, Y., & Jin, Y. (2007). Early triassic conodont–palynological biostratigraphy of the Meishan D section in Changxing, Zhejiang province, South China. *Palaeogeography, Palaeoclimatology, Palaeoecology*, *252*(1-2), 4–23.

Zhang, K., Yin, H., Tong, J., Jiang, H., & Luo, G. (2013). China "Golden Spikes": Global standard sections and points research. In C. A. o. S. Nanjing Institute of Geology and Palaeontology (Ed.), *三叠系下三叠统印度阶全球标准层型剖面和点位 [GSSP of the Induan Stage]* (pp. 281–319). Zhejiang University Press.

Zhang, M. (1976). A new species of helicoprionid shark from Xizang. *Chinese Journal of Geology*, *11*(4), 332–336.

Zhang, S., & Jin, Y. G. (1976). Late Paleozoic brachiopods from the Mount Jolmo Lungma region. *A report of scientific expedition in the Mount Jolmo Lungma Region (1966-1968)*, *2*, 159–242.

Zhao, J., Liang, X.-l., & Zheng, Z. (1978). Late Permian cephalopods of south China. *Palaeontologia Sinica*, *154*(12), 1–194.

Zhao, J., Sheng, J.-z., Yao, Z.-q., Liang, X.-l., Chen, C.-z., Rui, L., & Liao, Z.-t. (1981). The Changhsingian and Permian-Triassic boundary of South China. *Bull Nanjing Inst Geol Paleont Acad Sinica*, *2*, 1–85.

Zhao, X., & Tong, J. (2010). Two episodic changes of trace fossils through the Permian-Triassic transition in the Meishan cores, Zhejiang Province. *Science China Earth Sciences*, *53*(12), 1885–1893.

Zhuravlev, A. V., Plotitsyn, A. N., & Gruzdev, D. A. (2019). Carbon Isotope Ratios in the Apatite-Protein Composites of Conodont Elements—Palaeobiological Proxy. In *Processes and phenomena on the boundary between biogenic and abiogenic nature* (pp. 749–764). Springer.
